# Mutant IDH uncouples p53 from target gene regulation to disable tumor suppression

**DOI:** 10.1101/2024.09.30.615916

**Authors:** Cole Martin, William B. Sullivan, Jacqueline Brinkman, Deena Scoville, Jossie J Yashinskie, Sha Tian, Riccardo E. Mezzadra, Yu-Jui Ho, Richard P. Koche, Timour Baslan, Jesse Raab, David Corcoran, Lydia W.S. Finley, Scott W. Lowe, John P. Morris

## Abstract

p53 prevents tumor initiation and progression via transcriptional regulation of target gene networks. Here, we find that cancer-associated mutations in isocitrate dehydrogenase (IDH) can uncouple p53 activity from tumor suppression by perturbing chromatin states that determine target gene expression. Mutant IDH impairs tumor regressions and promotes the outgrowth of cancer cells with transcriptionally active, wild-type p53 in a mouse model of liver cancer where restoration of p53 activity results in tumor clearance. Mutant IDH alters p53 target gene expression through the oncometabolite 2-hydroxyglutarate (2-HG), an inhibitor of alpha-ketoglutarate (αKG)-dependent chromatin remodeling enzymes, without preventing p53 accumulation or global genomic binding. Rather, mutant IDH alters chromatin accessibility landscapes that dictate target gene expression, resulting in disabled upregulation of targets that execute tumor suppression. Specifically, mutant IDH disrupts the expression of pro-apoptotic p53 targets that enable p53-dependent tumor regressions, including the death ligand receptor Fas. Pharmacological inhibition of mutant IDH in TP53 wild-type cholangiocarcinoma cells, a tumor type where p53 and IDH mutations are mutually exclusive, potentiates p53 target gene expression and sensitizes cells to Fas ligand and chemotherapy-induced apoptosis. Therefore, we implicate the disruption of p53 target gene regulation as a reversible, oncogenic feature of cancer-associated IDH mutations.

**SIGNIFICANCE:** We find that chromatin states altered by cancer-associated IDH mutations intersect with transcriptional regulation of p53 target genes. This reversible interaction may represent a strategy to reinvigorate latent tumor suppression in IDH mutant, p53 wild-type tumors.

## INTRODUCTION

p53 is a sequence-specific transcription factor that acts broadly across tissues to constrain cancer initiation and progression (1). The distribution of cancer-associated mutations across its coding sequence and the dissection of its conserved functional domains strongly implicates the ability of p53 to regulate gene expression as an essential effector of tumor suppression. In support of this, TP53 is not only the most frequently mutated gene in cancer, but most alterations occur within its DNA binding domain (2,3). Furthermore, expression of transcriptionally incompetent p53 phenocopies the acceleration of tumor development observed upon genetic deletion in mice (4,5). While the capacity to regulate gene expression is therefore critical for its tumor suppressive function, how p53 activity culminates in the expression of target gene programs that impede tumorigenesis and how these networks are disabled in the absence of inactivating mutations remains incompletely understood.

p53 controls a vast set of direct and indirect target genes that regulate pleiotropic cellular programs implicated in tumor suppression. These programs include cell cycle arrest and senescence, apoptosis and other forms of cell death, metabolism, and programs of cellular differentiation (1,6,7). However, specific target genes and their downstream phenotypic modules are frequently dispensable for tumor suppression or fail to fully phenocopy p53 loss when inactivated (4,5,8). Furthermore, p53 target gene regulation is tissue and cell type-dependent with a limited number of universally regulated genes (6,9–11). This suggests that p53-mediated tumor suppression depends on mechanisms that selectively coordinate the regulation of target genes downstream of p53 transcription factor activity (2,12,13).

Metabolic rewiring that alters the abundance of metabolites that influence epigenetic states is emerging as one such regulatory paradigm (14,15). For example, we have recently demonstrated that p53 activity leads to the accumulation of the metabolite alpha-ketoglutarate (αKG) (16), which acts as both an intermediate in cellular metabolism and as a substrate for chromatin remodeling enzymes involved in gene regulation through histone and DNA modifications (17). αKG is implicated as an effector of p53 and increasing its levels in cancer cells recapitulates features of chromatin accessibility, αKG-dependent enzyme activity, and gene expression observed in response to restoration of wild-type p53 activity. Conversely, elevating levels of succinate, a competitive inhibitor of αKG-dependent enzymes, blunts the ability of p53 to slow tumor progression (16).

Like p53 inactivation, impaired αKG-dependent enzyme function is implicated in tumor development (18) and often corresponds with cancer progression following loss of p53 function (16,19). Cancers acquire loss-of-function mutations in αKG-dependent histone demethylases and DNA-modifying proteins (20,21) as well as mutations in metabolic regulators that cause the accumulation of competitive inhibitors of αKG-dependent enzymes (17). The most common of these mutations occurin isocitrate dehydrogenase (IDH) 1 and 2, which are cytoplasmic and mitochondrial enzymes, respectively, that normally catalyze conversion of isocitrate to αKG (22). Cancer-associated IDH1/2 mutations result in neomorphic activity that converts αKG into the oncometabolite R-2-hydroxyglutarate (2-HG). In turn, 2-HG inhibits αKG-dependent enzymes, resulting in altered chromatin states and gene expression programs that are implicated in tumorigenesis (17,18). Thus, p53 activity is responsible for increasing αKG levels and promoting the function of αKG-dependent enzymes, which are inhibited by oncometabolites, such as 2-HG, during tumor progression. Here, we test the hypothesis that cancer-associated IDH mutations could subvert p53-dependent tumor suppression by inhibiting αKG-driven tumor suppressive programs.

We find that mutant IDH can disable p53-dependent tumor suppression by perturbing control of p53 target gene expression. Mutant IDH acts through 2-HG to alter chromatin states that connect p53 activity to expression of direct and indirect target genes. Altered p53 target gene expression by mutant IDH is observed in both mouse and human liver cancer cells, with specific conservation of the ability of 2-HG to impair p53-dependent expression of the cell death receptor Fas. Thus, our work demonstrates that oncogenic IDH mutations can contribute to tumorigenesis by disrupting the expression of p53 target genes that execute critical features of tumor suppression and suggests that normalization of p53 transcriptional networks may be a therapeutic benefit of inhibiting mutant IDH in TP53 wildtype tumors.

## RESULTS

### Cancer-associated IDH mutants uncouple p53 from liver tumor suppression

IDH1/2 mutations that produce 2-HG are frequent in glioblastoma, acute myeloid leukemia, chondrosarcoma, and cholangiocarcinoma, where they are implicated in cancer initiation and progression (23). Remarkably, p53 and both IDH1 and 2 mutations display significant mutual exclusivity across multiple cholangiocarcinoma sequencing studies (24,25) (Fig. S1), indicating a potential genetic interaction between wild-type p53 and mutant IDH in liver tumorigenesis. To test if mutant IDH can influence p53 function in liver cancer cells, we introduced IDH2 mutants that result in accumulation of 2-HG into primary cancer cells derived from liver tumors induced by delivery of oncogenic <u>N</u>ras^G12D^ and linked expression of a doxycycline inducible (“on dox”), tetracycline response element regulated <u>p</u>53 <u>sh</u>RNA into adult hepatocytes (NP^sh^ cells, Fig. S2A, **Fig. 1A**) (26). This model allows for the restoration of wild-type p53 in liver cancer cells that progressed due to doxycycline-dependent (“ON DOX”) suppression of p53. Accordingly, removing doxycycline in vitro (“OFF DOX”) results in the rapid accumulation of p53 protein, corresponding with an increase in the cellular αKG:succinate ratio (Fig. S2B and C). As expected, constitutive expression of mutant IDH2^R140Q^ and IDH2^R172K^ in NP^sh^ cells resulted in 2-HG production, which was then potentiated by p53 restoration following the withdrawal of doxycycline, likely reflecting the increased levels of its substrate, αKG, induced by p53 activity (**Fig. 1B**). This highlights the NP^sh^ model as a relevant setting to test the effect of 2-HG, a potent, competitive inhibitor of αKG, on p53-dependent cell fate.

**Figure 1.**
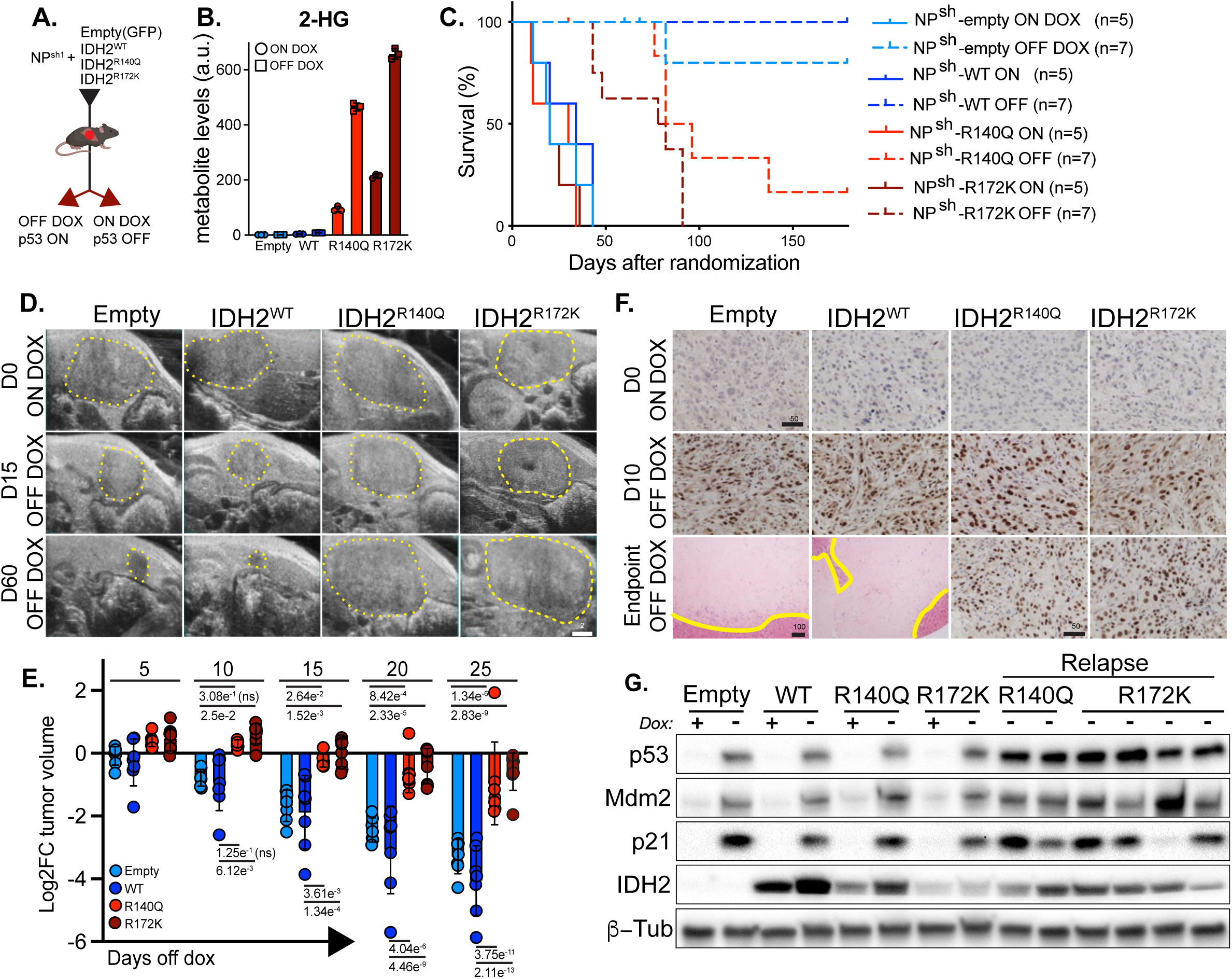
Mutant IDH disables p53 triggered liver tumor suppression. **a**. Schematic of delivery of IDH2 allelic series to NP^sh^-1 cells and *in vivo* analysis of liver tumor growth in response to p53 restoration following withdrawal of doxycycline (dox) chow. Mouse image from Biorender.com. **b**. GC-MS measurement of cellular 2-HG levels in NP^sh^-1 expressing empty vector, IDH2^WT^, IDH2^R140Q^, or IDH2^R172K^ on dox and 8 days after dox withdrawal (n=3 wells per condition). **c**. Survival curve of mice bearing orthotopic tumors derived from injection of NP^sh^-1 empty vector, IDH2^WT^, IDH2^R140Q^, or IDH2^R172K^ expressing cells from randomization into on dox and dox withdrawal groups, monitored for up to 6 months. **d.** Representative, longitudinal ultrasound imaging of orthotopic tumors generated from transplant of NP^sh^-1 cells expressing empty vector, IDH2^WT^, IDH2^R140Q^, or IDH2^R172K^ following withdrawal of doxycycline chow. Dashed lines outline tumor boundaries. **e.** Relative volume of indicated NP^sh^-1 orthotopic tumors following withdrawal of dox chow. **f.** Representative immunohistochemistry of p53 in indicated orthotopic tumors following withdrawal of doxycycline chow. Yellow lines indicate boundary of acellular tumor remnants in NP^sh^-1-empty vector and IDH2^WT^ transplants from H&E staining. **g**. Western blot of p53, Mdm2, and p21/Cdkn1a in parent NP^sh^-1 cells expressing empty vector, IDH2^WT^, IDH2^R140Q^, or IDH2^R172K^ on dox versus 8 days of p53 restoration by dox withdrawal and primary cell lines generated from relapsed R140Q and R172K expressing tumors. Scale bars: d: 2mM; f: 100 uM (H&E) or 50uM (p53 IHC). Significance in e calculated with one way ANOVA with Tukey correction for multiple comparisons.

p53 reactivation in established liver tumors resulting from orthotopic transplant of NP^sh^ cells triggers potent tumor suppression and rapid regressions in immune competent mice (26). To test if mutant IDH alters the effect of p53 on cell fate during liver tumor maintenance, we first tracked tumor size upon p53 restoration in control tumors generated from the injection of NP^sh^ cells expressing either GFP or GFP-linked wild-type (WT) IDH2 (NP^sh^-empty, NP^sh^-IDH2^WT^) as controls versus cells expressing GFP-linked mutant IDH2^R140Q^ or IDH2^R172K^ (NP^sh^-IDH2^R140Q^, NP^sh^-IDH2^R172K^). Doxycycline withdrawal from mice bearing control tumors resulted in precipitous regressions within 10-15 days and prolonged survival over 6 months of monitoring that corresponded with complete elimination of tumor cells (**Fig. 1C-E**). In contrast, expression of mutant IDH2^R140Q^ and IDH2^R172K^ significantly blunted initial tumor regressions following doxycycline withdrawal and led to frequent tumor regrowth after a period of stasis (IDH2^R140Q^, 6/7 mice, IDH2^R172K^ 7/7 mice versus IDH2^WT^, 0/7 mice, Vector, 1/7 mice, **Fig. 1C-E**, Fig. S2D). The effect of mutant IDH on tumor growth was specific to the setting of p53 restoration, as neither tumor growth nor survival in dox-fed (ON DOX) mice was altered by the expression of either IDH2^R140Q^ or IDH2^R172K^ (**Fig. 1C**, Fig. S2D). Tumor regressions triggered by p53 restoration were similarly attenuated by expression of mutant IDH2^R140Q^ and IDH2^R172K^ in orthotopic tumors generated by injection of an independently-derived NP^sh^ cell line (Fig S2E).

While expression of mutant IDH2 disabled tumor regressions following doxycycline withdrawal, this did not correspond with a failure of p53 restoration, as its protein levels were indistinguishable between IDH WT and IDH mutant cells *in vivo* (**Fig. 1F**). Remarkably, p53 remained strongly expressed in relapsed tumors, indicating a failure of p53-triggered tumor clearance without doxycycline independent reactivation of the p53 shRNA (IDH2^R140Q^ 2/6 relapsed tumors, IDH2^R172K^ 4/7 relapsed tumors, **Fig. 1F**, Fig. S2F). Similarly, mutant IDH did not prevent p53 accumulation or induction of well-established p53 targets Cdkn1a/p21 and Mdm2 *in vitro* upon doxycycline withdrawal, which also remained strongly expressed in cell lines from relapsed tumors that maintained high levels of p53 and elevated 2-HG levels (**Fig. 1G**, Fig. S2G). Thus, consistent with both the mutual exclusivity of TP53 and IDH1/2 mutations in human cholangiocarcinoma (Fig. S1) and the reported accumulation of WT p53 in mutant IDH disease (27), cancer-associated IDH mutations can disable p53-triggered liver tumor elimination and establish a cell state where otherwise tumor-suppressive levels of p53 protein are tolerated during liver tumor progression.

### Mutant IDH disconnects p53 activity from target gene expression

Given the key role that transcriptional regulation plays in p53-mediated tumor suppression asked if mutant IDH effects p53-dependent gene expression in NP^sh^ cells. Bulk RNA sequencing (RNA-SEQ) was performed on NP^sh^-IDH2^WT^ and NP^sh^-IDH2^R172K^ cells maintained in doxycycline (ON DOX, p53 off) versus 8 days after dox withdrawal (OFF DOX, p53 on). To determine if gene expression changes were dependent on sustained oncometabolite production, all cells were pretreated for 7 days with either DMSO control or AG-221 (29), an inhibitor of mutant IDH2 that effectively reduced 2-HG levels (Fig. S3A and B).

As illustrated by principal component analysis, mutant IDH enforced a 2-HG-dependent effect on gene expression in NP^sh^ cells (PC2) (**Fig. 2A**). Consistent with a functional interaction between mutant IDH and transcriptional regulation by p53, 2-HG resulted in a remarkable perturbation of p53 target gene expression. p53 activity in NP^sh^-IDH2^WT^ cells resulted in 2,235 differentially expressed genes (1574 upregulated, fold change≥2; 661 downregulated, fold change≤-2, adjP<0.05, **Fig. 2B**). Mutant IDH significantly altered expression of ∼20% of p53 regulated genes (fold change≥+/-2, p_adj_<0.05, 454/2235) in a largely reversible, 2-HG dependent fashion, with expression of ∼55% of significantly altered genes rescued by blocking 2-HG (252/454) (**Fig. 2B and C**). Of note, mutant IDH predominantly led to blunted upregulation of and repression of p53 targets, although some displayed enhanced activation or repression (Supplementary Table 1). Thus, mutant IDH interferes with p53-dependent gene expression, despite robust accumulation of p53 protein.

**Figure 2.**
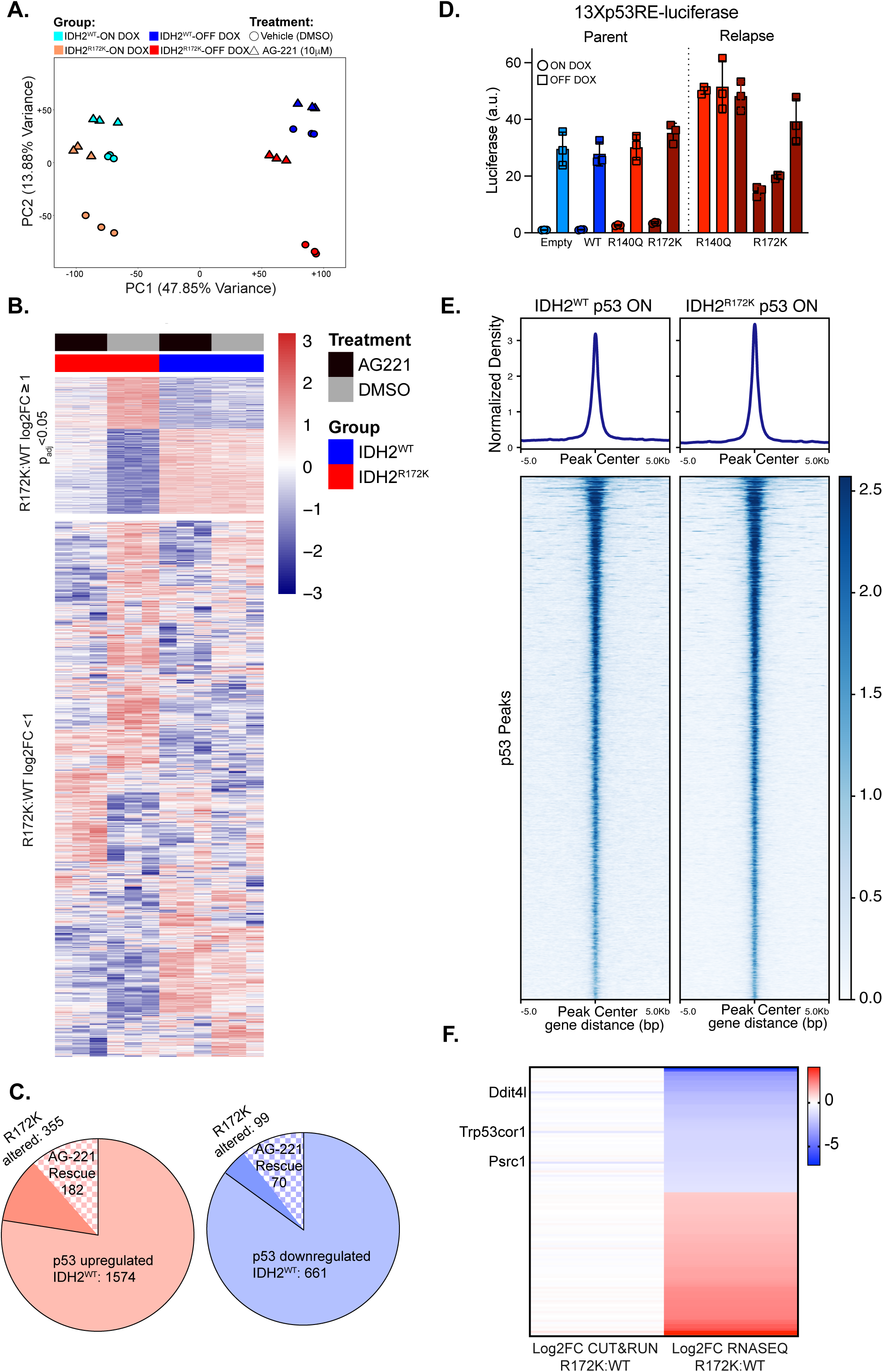
Mutant IDH uncouples p53 activity from target gene regulation. a. Principal component analysis of gene expression (RNA-SEQ) measured in NP^sh^-1-IDH2^WT^ and NP^sh^-1-IDH2^R172K^ cells following p53 restoration (on dox versus off dox) treated with AG-221. b. Heatmap depicting Z-scored expression of genes differentially expressed between NP^sh^-1- IDH2^WT^ ON DOX cells (p53 off) and NP^sh^-1-IDH2^WT^ OFF DOX cells (p53 on). The heatmap shows expression of these p53-regulated genes in NP^sh^-1-IDH2^WT^ OFF DOX cells (p53 on) and NP^sh^-1- IDH2^R172K^ OFF DOX cells (p53 on) treated with either DMSO or AG-221 (n=3 per condition). Top cluster represents genes differentially expressed in off dox NP^sh^-1-IDH2^WT^ versus NP^sh^-1- IDH2^R172K^ cells by greater than 2-fold, p_adj_<0.05). c. Quantification of p53 up and downregulated targets differentially expressed greater than 2-fold, p_adj_<0.05 in the setting of mutant IDH and reversed by treatment with AG-221 (IDH2^WT^ versus AG221 treated IDH2^R172K^ p_adj_>0.05). d. Relative p53 reporter activity (13Xp53RE-luciferase) in parent NP^sh^-1 empty vector, IDH2^WT^, IDH2^R140Q^, or IDH2^R172K^ and NP^sh^-1-IDH2^R140Q^ and NP^sh^-1-IDH2^R172K^ relapse cells (n=3 wells per condition). e. Profile and tornado plots of p53 peaks identified in NP^sh^-1-IDH2^WT^ and NP^sh^-1- IDH2^R172K^ cells 8 days after dox withdrawal by CUT&RUN (average of n=3 replicates per condition). f. Average fold change of p53 binding magnitude (CUT&RUN) associated with target genes up or downregulated by RNA-SEQ in NP^sh^-IDH2^R172K^ versus NP^sh^-IDH2^WT^ cells (fold change≥+/-2, p_adj_<0.05).

Transcriptional regulation by p53 is predicated on nuclear accumulation and transactivation of direct target genes following response element engagement (13,28). While mutant IDH altered p53-triggered gene expression in NP^sh^ cells, it neither prevented nuclear p53 accumulation nor its ability to stimulate response element-driven reporter gene activity in parental cell lines or primary cells derived from relapsed tumors (Fig. S3C, **Fig. 2D**) (29). Therefore, in this setting of oncogenic stress, mutant IDH perturbs the regulation of p53 target gene networks without disabling p53 stability or its capacity to interact with response elements and the basal transcriptional machinery necessary for target transactivation (30).

Next, we tested if changes in p53 chromatin occupancy could account for altered target gene regulation by mutant IDH. We mapped global p53 binding through cleavage under targets and release using nuclease profiling (“CUT&RUN” (31)) following doxycycline withdrawal in NP^sh^- IDH2^WT^ and NP^sh^-IDH2^R172K^ cells. This profiling approach was robust, as we observed enrichment in signal at high-confidence p53 target genes (e.g. p21/Cdkn1a) that were similarly induced upon p53 restoration in both IDH WT and mutant cells (Fig. S3D), as well as a significant enrichment of p53 and p53 family motifs in identified peaks (Fig. S3E). Consistent with the ability of p53 to engage its response element in reporter gene experiments, profiling of p53 binding in NP^sh^- IDH2^R172K^ and NP^sh^-IDH2^WT^ cells revealed neither a reduction in the total number of peaks observed nor a change in their average magnitude (Fig. S3F, **Fig. 2E**). Thus, while mutant IDH resulted in greater than 2-fold differential expression of 97 of 931 candidate direct p53 targets identified in NP^sh^-IDH2^WT^ cells, here defined by at least one p53 peak within 10 kb of the gene body (32), this was not reflected by commensurate change in p53 binding (Fig. S3G). Of putative direct targets altered by mutant IDH by greater than 2-fold, only 3 displayed a significant difference in p53 binding (Ddit4l, Trp53cor1, Psrc1, **Fig. 2F**, Fig. S3G). Therefore, mutant IDH perturbs p53 dependent transcriptional regulation of target genes in mouse liver cancer cells without preventing its accumulation or disrupting its profile of genomic binding, indicating that mutant IDH alters determinants of p53 target gene expression downstream of chromatin engagement.

### Mutant IDH perturbs chromatin states that determine p53 target gene expression

Cancer-associated IDH mutations are linked to altered epigenetic states through inhibition of αKG-dependent chromatin remodelers, including histone demethylases and DNA modifying enzymes, resulting in 2-HG-dependent patterns of gene expression (33). As cellular chromatin states can determine p53 target gene expression (13,34), we asked if the perturbed regulation of p53-dependent genes we observed corresponded with changes in histone methylation and DNA modifications influenced by mutant IDH. Expression of mutant IDH2^R172K^ in NP^sh^ cells resulted in increased levels of histone methylation regulated by αKG-dependent demethylase activity (e.g. H3K4me^3^, H3K9me^2^, H3K9me^3^, and H3K36me^3,^ Fig. S4A) (35). Of note, expression of mutant IDH maintained elevated levels of H3K4me^3^ and H3K9me^2^ following p53 restoration, two marks that were reduced following p53 reactivation in IDH WT cells in a fashion consistent with an increased αKG:succinate ratio (Fig. S2B, Fig. S4A). Similarly, p53 restoration increased levels of 5-hmC, the product of the αKG-dependent Tet enzymes (36), in the setting of WT IDH, but this effect blunted by mutant IDH (Fig. S4B).

To determine if perturbed levels of αKG dependent chromatin modifications are functionally linked to altered p53 target gene regulation by mutant IDH, we restored p53 activity in NP^sh^-IDH2^WT^ control cells treated with JIB-04 (37), an inhibitor of Jumonji-C αKG dependent demethylases that increased histone methylation in a similar fashion as mutant IDH (**Fig. 3A**). Inhibiting global levels of histone demethylation was sufficient to blunt upregulation of p53 direct and indirect target genes disabled by 2-HG (e.g., Ephx1 and P2ry2, respectively, **Fig. 3B and C**), implicating the inhibition of αKG dependent chromatin modifying enzymes by mutant IDH in altering both direct and indirect p53 target gene expression.

**Figure 3.**
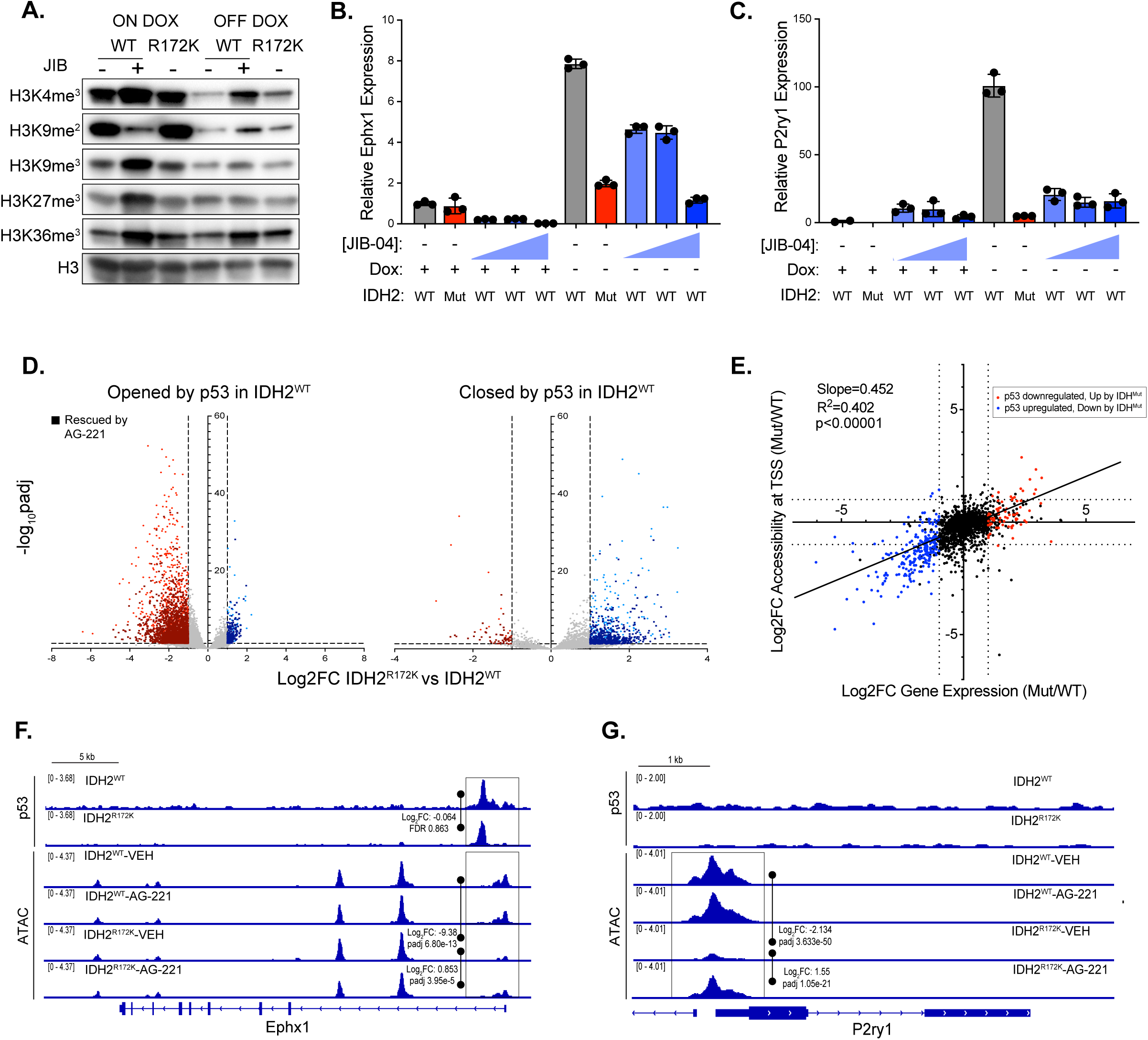
Mutant IDH perturbs chromatin landscapes that connects p53 activity to direct and indirect target gene expression. a. Western blot of H3K4me3, H3K9me2, H3K9me3, H3K27me3, and H3K36me3 in NP^sh^-1- IDH2^WT^ cells treated with 96 hours of 1 uM JIB-04, and NP^sh^-1-IDH2^R172K^ cells, on and 8 days off dox. b,c. qRT-PCR of Ephx1, b, and P2ry1, c, in NP^sh^-1-IDH2^WT^ cells treated with 96 hours of vehicle (grey bars) or JIB-04 (0.2, 0.5 and 2 uM, blue bars) and vehicle treated NP^sh^-1-IDH2^R172K^ cells (red bars) on and off dox for 8 days. Expression was normalized to 36B4 and presented relative to NP^sh^-1-IDH2^WT^ cells on dox (n=3 wells per condition). d. Volcano plots depicting the accessibility (ATAC-SEQ) of p53 dependent opened (left) and closed (right) loci in NP^sh^-1-IDH2^WT^ versus NP^sh^-1-IDH2^R172K^. Highlighted points indicate differentially accessible peaks (fold change>2, p_adj_<0.05). Squares denote loci rescued by AG-221 in NP^sh^-1-IDH2^R172K^. e. Correlation of log2FC gene expression and log2FC TSS associated accessibility magnitude in NP^sh^-1-IDH2^WT^ versus NP^sh^-1-IDH2^R172K^ cells 8 days after p53 restoration (dox off). Blue/Red genes are up/downregulated by dox withdrawal in NP^sh^-1-IDH2^WT^ and down/upregulated in NP^sh^-1-IDH2^R172K^ (fold change≥+/-2, p_adj_<0.05). f.g. Average p53 CUT&RUN and ATAC-SEQ treated with AG-221 (n=3 each condition) profiles of the Ephx1 locus, f, and P2ry1 locus, g, in NP^sh^-1-IDH2^WT^ versus NP^sh^-1-IDH2^R172K^ cells 8 days after p53 restoration.

Inhibition of αKG-dependent, chromatin-modifying enzymes by mutant IDH results in altered chromatin accessibility landscapes that connect to gene expression (38). As remodeled chromatin accessibility corresponds with p53 activation and target gene expression (16,39), we asked if mutant IDH altered accessibility landscapes associated with p53 restoration in NP^sh^ cells through transposase-accessible chromatin with sequencing (ATAC-SEQ, Fig. S3A). As observed in RNA-SEQ profiling, principal component analysis of the ATAC-SEQ data revealed 2-HG-dependent perturbations of global chromatin accessibility (PC2) (Fig. S4C). Mirroring its alteration of p53- dependent gene expression, mutant IDH reduced both gains and losses in chromatin accessibility induced by p53 activity in a 2-HG-dependent fashion. Approximately 14% of peaks opened upon p53 restoration were perturbed by mutant IDH (fold change≥+/-2, p_adj_<0.05, 3164/23022 peaks; 2500/3164, ∼79% rescued), and similarly ∼16% of peaks closed by p53 were perturbed by mutant IDH (fold change≥+/-2, p_adj_<0.05, 783/4801 peaks; 617/783, ∼79% rescued, **Fig. 3D**, Fig. S4D).

Altered chromatin accessibility extended to transcriptional start sites (TSS) of p53 target genes in a fashion that correlated directly with their expression (**Fig. 3E**). Thus, mutant IDH enforced a 2- HG-dependent disconnect between p53 activity and TSS accessibility at subsets of both directly bound targets (e.g. Ephx1, **Fig. 3F**) and indirect targets (e.g. P2ry1, **Fig. 3G**), with others, including well-established targets such as Mdm2, unchanged by 2-HG (Fig. S4E). Taken together, these data demonstrate that mutant IDH influences αKG-dependent chromatin landscapes that selectively determine the expression of p53 direct and indirect targets downstream of p53 transcription factor activity.

### Mutant IDH disables p53 triggered tumor regressions by inhibiting Fas expression and Fas-mediated apoptosis

p53 transcriptionally regulates an array of cell fates, including cell cycle arrest, senescence, and apoptosis, that are contextually involved in tumor suppression (4,40,41). Despite its profound effect on p53 target gene expression, mutant IDH did not prevent reduced proliferation or emergence of senescence associated beta-galactosidase (SA β–gal) activity in response to p53 restoration *in vitro* (Fig. S5A-C). Ki67 labeling revealed similarly reduced proliferation of NP^sh^ control versus mutant IDH expressing cells in tumors 10 days after doxycycline withdrawal (Fig. S5D). Therefore, in the context of NP^sh^ liver cancer cells, the ability of mutant IDH to disable p53 dependent tumor suppression is not due to disabled proliferative arrest or the initiation of senescence.

On the other hand, gene set enrichment analysis (GSEA) revealed enrichment of several pathways implicated as tumor suppressive effectors of p53 in NP^sh^-IDH2^WT^ versus NP^sh^-IDH2^R172K^ cells upon p53 restoration, including function of the spliceosome (42), DNA damage repair (43), and apoptosis (44) (Fig. S6A). This included attenuated expression of mediators of extrinsic apoptosis, e.g., the death receptor Fas (45), as well as genes involved in intrinsic apoptosis execution including Pmaip1/Noxa (46), Caspase 7 (47), and Parp10 (48) (Fig. S6A). Upregulation of these genes by p53 was reduced by mutant IDH in a largely 2-HG dependent fashion, with p53 dependent expression of Fas, Caspase-7, and Parp10 rescued by inhibiting mutant IDH and only Pmaip1 remaining suppressed (**Fig. 4A and B**, Fig. S6B). Consistent with the ability of mutant IDH to perturb chromatin states associated with p53 target gene regulation, decreased expression of pro-apoptotic genes corresponded with reduced TSS associated chromatin accessibility for both directly bound (Fas, Pmaip1) and indirect targets (Casp7, Parp10) (Fig. S6C).

**Figure 4.**
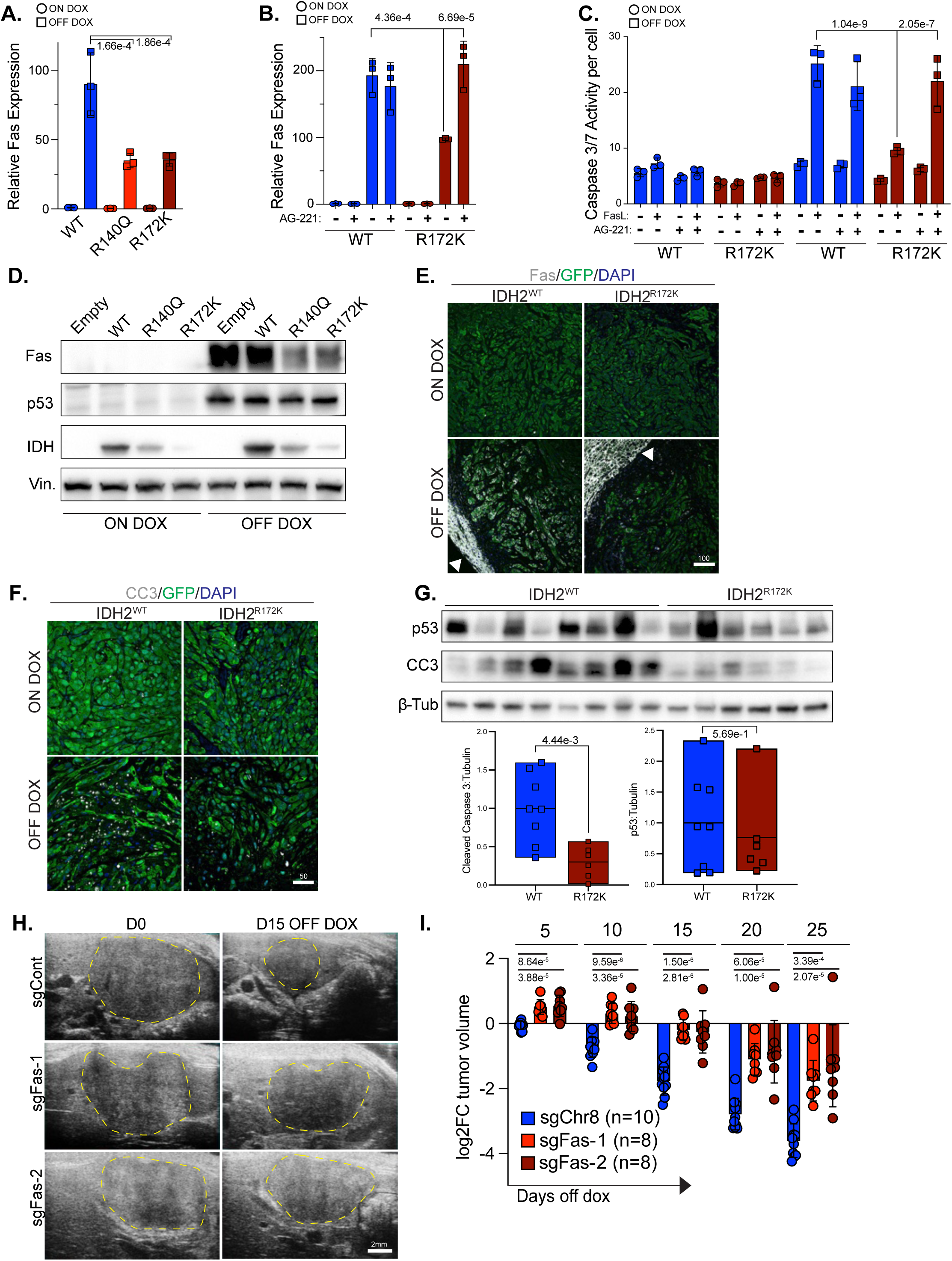
Mutant IDH disconnects p53 from liver tumor suppression by disabling Fas expression and Fas dependent apoptosis. **a.** qRT-PCR of Fas in NP^sh^-1-IDH2^WT^, IDH2^R140Q^, and IDH2^R172K^ cells on and off dox for 8 days normalized to 36b4. **b**. qRT-PCR of Fas in NP^sh^-1-IDH2^WT^ and NP^sh^-1-IDH2^R172K^ cells on and off dox for 8 days treated with AG-221 normalized to 36b4 (n=3 wells per condition). **c**. Cleaved caspase 3/7 activity measured by Caspase-Glo 3/7 luminescence assay in NP^sh^-1-IDH2^WT^ and NP^sh^-1-IDH2^R172K^ cells treated with AG-221, on dox or off dox for 8 days, treated with FasL (n=3 wells per condition) **d.** Western blot for Fas in NP^sh^-1-IDH2^Empty^, NP^sh^-1-IDH2^WT^, NP^sh^-1-IDH2^R140Q^, and NP^sh^-1-IDH2^R172K^ cells on and off dox for 8 days. **e, f**. Representative co-immunofluorescent staining of Fas, **e**, or cleaved caspase 3 (CC3), **f**, and GFP in orthotopic NP^sh^-1-IDH2^WT^ and NP^sh^- 1-IDH2^R172K^ tumors on dox and 10 days after withdrawal of dox. **g**. Western blot (top) of p53 and cleaved caspase 3 in lysates from orthotopic NP^sh^-1-IDH2^WT^ and NP^sh^-1-IDH2^R172K^ tumors on dox and 10 days after withdrawal of dox (β**-**Tub=β**-**Tubulin). Quantification (bottom) of CC3 and p53 western blots normalized to β**-**Tubulin. **h**. Representative longitudinal ultrasound imaging of orthotopic tumors generated from transplant of NP^sh^-1 cells expressing sgChr8, sgFas-1, and sgFas-2 on dox and 15 days after withdrawal of dox. Dashed lines outline tumor boundaries. **i**. Relative volume of indicated orthotopic tumors following withdrawal of dox chow. a,b,c,i significance calculated with one way ANOVA with Tukey correction for multiple comparisons. g. significance calculated by students t-test. Scale bars e, 100µM, f, 50µM, h, 2mM.

Therefore, we next asked if uncoupling of p53 activity from expression of pro-apoptotic target genes altered apoptotic sensitivity in NP^sh^ cells *in vitro* and *in vivo*. p53 restoration alone in NP^sh^- IDH2^WT^ cells did not result in increased apoptosis, as measured by caspase 3/7 activity (**Fig. 4C**). However, consistent with elevated Fas expression, p53 restoration led to increased sensitivity to apoptosis triggered by treatment with Fas ligand (FasL), an effect that was blocked by mutant IDH in a 2-HG-dependent fashion (**Fig. 4C**). Accordingly, mutant IDH reduced the upregulation of Fas protein by p53 *in vitro* (**Fig. 4D**) and *in vivo* (**Fig. 4E**), which was consistent with reduced levels of cleaved caspase 3 following p53 reactivation as visualized by immunofluorescence and measured in tumor lysates (**Fig. 4F and G**).

p53 restoration in orthotopic NP^sh^ tumors in immune competent hosts results in recruitment and activation of CD8+ T-cells characterized by increased FasL expression that are critical for tumor regressions (26). Therefore, we nominated reduced levels of Fas, and thus decreased sensitivity to extrinsic FasL, as a possible mechanism whereby mutant IDH blunts p53-triggered tumor suppression. Accordingly, sgRNA-mediated knockout of Fas in NP^sh^-1 cells not only prevented FasL-mediated apoptosis *in vitro* (Fig. S6D and E), but also phenocopied the effect of mutant IDH on p53 induced tumor regressions in orthotopic liver transplants (**Fig 4H**, I). Therefore, by inhibiting p53-dependent Fas expression, mutant IDH permits tolerance of p53 activation *in vivo* that would otherwise result in the sensing of extrinsic apoptotic cues and the clearance of tumor cells in response to p53 activation.

### Inhibiting mutant IDH expands p53 target gene expression and sensitizes human cholangiocarcinoma cells to FasL and chemotherapy-induced apoptosis

Given the ability of mutant IDH to disconnect p53 activity from target gene expression in NP^sh^ mouse liver cancer cells and the mutual exclusivity of p53 and IDH1/2 mutations in human cholangiocarcinoma (**Fig. S1**), we next tested if mutant IDH intersects with p53-dependent gene regulation in human cholangiocarcinoma cells. To this end, we identified two cholangiocarcinoma cell lines with wild-type TP53 and oncogenic IDH mutations, RBE (IDH1^R132S^) and SNU-1079 (IDH1^R132C^), and treated these cells with the Mdm2 inhibitor nutlin to activate p53 in the setting of mutant IDH1 inhibition by the clinically approved inhibitor Ivosidenib (AG-120) (49,50). Of note, since IDH inhibitors have been shown to rewire chromatin states over weeks to change cell fate (51), AG-120 was applied for at least 21 days prior to analysis. Over this time frame 2-HG was persistently reduced (**Fig. 5A**), resulting in decreased histone methylation consistent with increased αKG-dependent demethylase activity (**Fig. 5B**). Treatment with the Mdm2 inhibitor nutlin resulted in p53 accumulation and increased expression of p53 target genes Mdm2 and p21 regardless of 2-HG levels (**Fig. 5C**), indicating that like in the NP^sh^ model above, mutant IDH does not prevent p53 accumulation or interfere with its ability to regulate all target genes in human cholangiocarcinoma cells. Of note, inhibiting mutant IDH did not lead to an increase in p53 transcript levels (52) (**Fig. S7A**).

**Figure 5.**
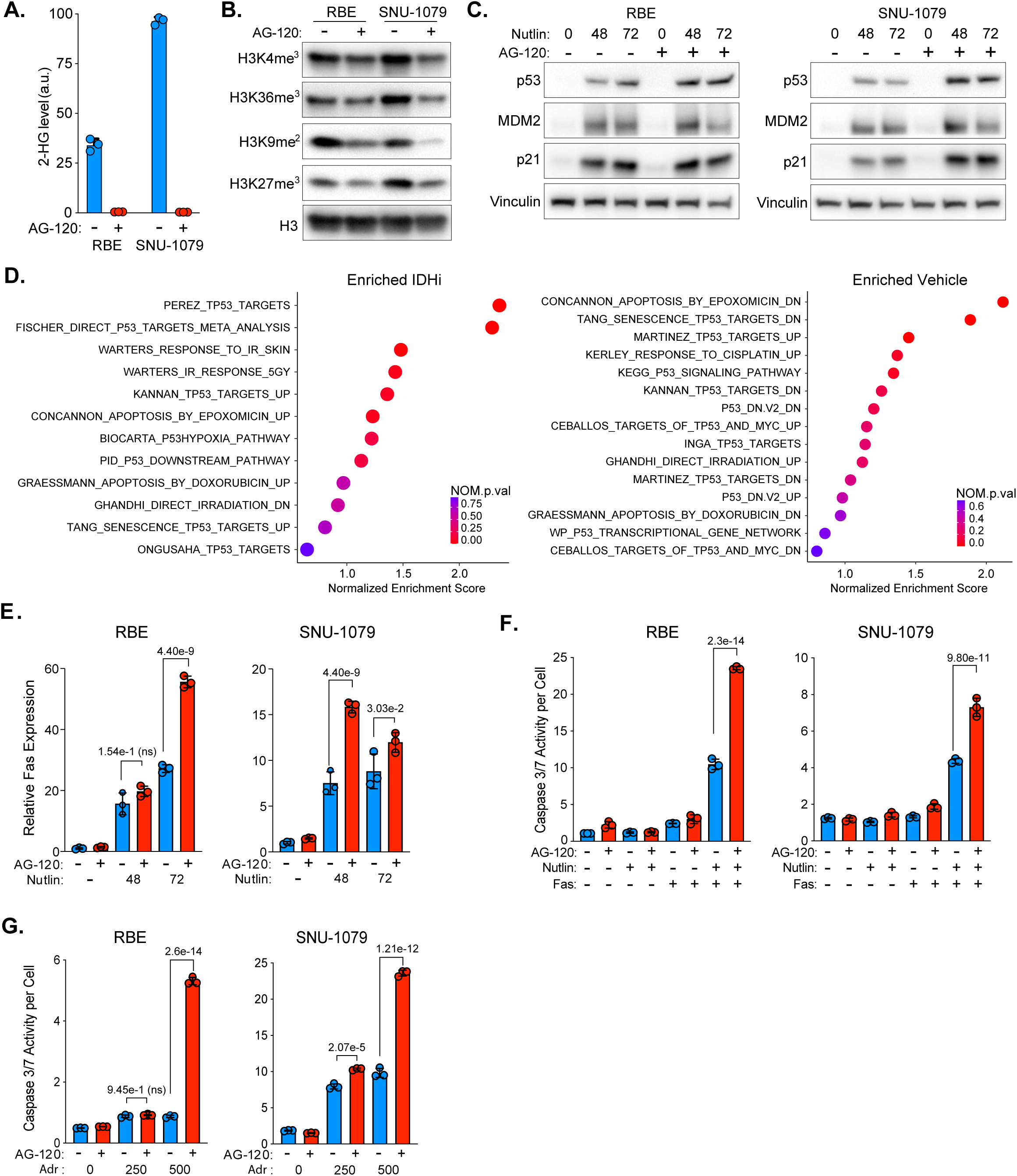
Inhibiting mutant IDH in cholangiocarcinoma cells increases p53 dependent gene expression and sensitizes to Fas and chemotherapy induced apoptosis. **a.** GC-MS measurement of cellular 2-HG levels in RBE and SNU-1079 cholangiocarcinoma cells treated with AG-120 for 21 days (n=3 wells per condition). **b**. Western blot of histone 3 methylation in RBE and SNU-1079 cells after at least 21 days of AG-120 treatment. **c**. Western blot of p53, MDM2, and CDKN1A/p21 in RBE (left) and SNU-1079 (right) cells pretreated with AG-120 and given 5 uM nutlin for 48 and 72 hours. **d**. RNA-SEQ GSEA analysis results of indicated p53 activity associated gene sets enriched in 5 uM nutlin treated RBE cells pretreated with AG-120 (left) versus vehicle (right). **e**. qRT-PCR of FAS in RBE (left) and SNU-1079 (right) pre-treated with AG-120 and 5 uM nutlin for 48 and 72 hours normalized to ACTB (n=3 wells per condition). **f**. Cleaved caspase 3/7 activity measured by Caspase-Glo 3/7 luminescence assay in RBE (left) and SNU-1079 (right) cells treated with AG-120, 5 uM nutlin, and FasL (n=3 wells per condition). **g**. Cleaved caspase 3/7 activity measured by Caspase-Glo 3/7 luminescence assay in RBE (left) and SNU-1079 (right) cells treated with AG-120 and indicated Adriamycin concentrations (n=3 wells per condition). e,f,g significance calculated with one way ANOVA with Tukey correction for multiple comparisons.

To test if mutant IDH acts through 2-HG to alter the transcriptional response to p53 activation in human cholangiocarcinoma cells, we performed RNA-SEQ comparing nutlin and AG-120-treated RBE cells to RBE cells treated with nutlin alone. This expression profiling approach revealed expanded p53-regulated gene expression upon inhibition of mutant IDH. Gene set enrichment analysis (GSEA) of biological hallmarks revealed enrichment of the p53 pathway signature in nutlin and AG-120 treated cells, while gene sets reflecting pathways suppressed by p53 activity remained enriched in nutlin and vehicle treated cells (e.g. MYC and E2F targets, **Fig. S7B**). We extended this analysis to a set of curated gene lists reflecting genes upregulated in TP53 wild-type cells and tissues exposed to genotoxic stressors as well as genes reflecting decreased p53 activity (e.g. p53 deletion or expression of negative regulators of p53 targets such as MYC, **Supplemental Table 2**). Consistent with perturbation of genes up and downregulated by p53 in mouse liver cancer cells, reducing 2-HG levels in human cholangiocarcinoma cells led to increased expression of gene signatures associated with p53 activation. Similarly, genes downregulated by p53 or derepressed by p53 inactivation were enriched in vehicle treated cells where 2-HG levels remained high (**Fig. 5D**). Thus, mutant IDH interferes with p53 target gene regulation in cholangiocarcinoma cells, suggesting active antagonism of p53-dependent gene expression by mutant IDH that persists through tumor development and can be pharmacologically reversed.

Next, we tested if perturbed regulation of p53-dependent gene expression in cholangiocarcinoma cells extended to specific pro-apoptotic targets and resulted in altered sensitivity to apoptotic cues. As in NP^sh^ cells, mutant IDH inhibited both Fas expression in response to nutlin treatment and apoptosis in response to FasL in both RBE and SNU-1079 cells (**Fig. 5E and F**). Furthermore, inhibition of mutant IDH increased the sensitivity of both RBE and SNU-1079 cells to the DNA damaging agent Adriamycin, which corresponded with increased apoptosis (**Fig. 5G**, **Fig. S7C and D**). Thus, mutant IDH perturbs p53-triggered expression of the Fas death receptor in conserved fashion, and reversibly blunts p53-dependent sensitivity to apoptosis triggered by FasL or chemotherapeutic agents that result in p53 accumulation.

## DISCUSSION

Transcriptional regulation is essential for tumor suppression by p53 (7). Here, we demonstrate that chromatin states established by cancer-associated IDH mutations intersect with the control of p53 target gene expression, resulting in uncoupling of p53 activity from gene regulatory networks that result in tumor suppression. Mutant IDH acts through 2-HG, a competitive inhibitor of αKG-dependent enzymes, to perturb chromatin accessibility landscapes that determine the magnitude of expression of both direct and indirect p53 targets. Pharmacological inhibition of αKG-dependent chromatin remodeling enzymes phenocopies the effect of mutant IDH on selective p53 target gene expression, implicating αKG-dependent enzyme function as a determinant of p53-dependent transcriptional regulation. Mutant IDH impairs expression of pro-apoptotic p53 targets including the death receptor Fas, which facilitates sensing of extrinsic cell death cues, leading to tolerance of p53 activity that would otherwise result in cell death. The ability of mutant IDH to selectively perturb p53 target gene expression extends to human cholangiocarcinoma cells that reflect mutual exclusivity between p53 and IDH1/2 mutations. Taken together, this study establishes perturbation of p53 target gene regulation as a reversible, oncogenic feature of mutant IDH that persists through tumor development and could be exploited to induce latent p53 anti-tumor effects in IDH mutant, p53 WT tumors.

p53 is implicated in the control of hundreds to thousands of direct and indirect transcriptional targets in a context-dependent fashion (6). However, the molecular logic that connects p53 activity to target gene regulation remains incompletely understood. Current models suggest that the transcriptional output of p53 is regulated at multiple levels to establish its target gene networks, including its degree and duration of accumulation (53,54), selective target gene binding (55,56), as well as the local chromatin context at target genes (34,57). Our study demonstrates that while mutant IDH does not prevent p53 accumulation or DNA binding, it results in remodeling of chromatin accessibility landscapes directly associated with expression of specific targets. Critically, pharmacological inhibition of αKG-dependent demethylases that mimics the effect of mutant IDH on histone methylation is sufficient to recapitulate the effect of mutant IDH on p53 target gene expression. This is consistent with a role for αKG and αKG-dependent chromatin regulation as a key effector arm of p53 that is disabled by mutant IDH through 2-HG. Thus, our work supports a “feed-forward” model where the increased levels of αKG induced by p53 activity (observed here, (16,58)) reinforce chromatin states that determine target gene expression and ultimately, p53-dependent cell fate. Both the utilization of nutrients that can contribute to αKG levels and αKG-dependent enzyme expression display tissue and cell type dependency (17), which we speculate could contribute to the establishment of context-dependent p53 gene regulatory outputs and potentially the selection for mutations, like those in IDH1/2, that result in inhibition of these enzymes during tumorigenesis.

While we find that mutant IDH can perturb expression of a substantial portion of p53-regulated genes, this intersection with target gene regulation is not universal. For example, expression of well-established direct targets involved in cell cycle arrest (e.g. Cdkn1a) and p53 feedback inhibition (e.g. Mdm2) are not altered by mutant IDH, whereas genes involved in sensing and executing cell death cues (e.g. Fas) are disabled. Functionally, this results in a disconnect between coincident cell fates induced by p53 (59) where, for example, cells undergoing cell cycle arrest and senescence are no longer subject to apoptosis in response to cell-extrinsic FasL. Perturbation of p53 target genes involved in tumor suppression through interpreting cell-extrinsic signals is consistent with the function of mutant IDH as an atypical oncogene (22) that interferes with cellular programs that depend on interactions between cellular compartments. Indeed, sustained inhibition of 2-HG in IDH mutant cholangiocarcinoma cells does not result in reduced proliferation in vitro (60), and IDH inhibitors result in decreased growth of IDH mutant cholangiocarcinoma only in the setting of an intact immune microenvironment (22). While the current work focuses on the ability of mutant IDH to disable p53-dependent sensing of apoptotic cues, our profiling identifies a number of p53 target genes altered by mutant IDH that may contribute to disconnecting other facets of tumor suppression.

IDH1 and 2 are the most commonly altered metabolic enzymes across human cancer (22). Here, we studied the interaction of p53 and mutant IDH specifically in mouse liver cancer and human cholangiocarcinoma cells due to the remarkable mutual exclusivity between p53 and IDH mutation observed in this liver cancer type (**Fig S1**) (24,25). Of note, this pattern of mutual exclusivity is not a universal feature of IDH mutant tumors as p53 and IDH mutation frequently co-occur in other tumors, prominently glioma (61). These tissue dependent configurations of IDH and TP53 mutation may be a function of contextual or temporal roles for p53 target gene regulation during tumor development. For example, IDH mutations are acquired before TP53 mutations in glioma (62) and have been shown to drive features of cancer initiation in neural cells but then become dispensable for maintenance of transformation (63).

In summary, our work demonstrates that the effects of cancer-associated IDH mutations on chromatin states converge on p53 target gene regulation. It provides functional support for a metabolite-sensitive epigenetic mechanism through which p53-dependent increases in cellular αKG promote tumor suppressive transcriptional programs that can be selected against during tumorigenesis via acquisition of mutations that produce 2-HG. In some cancers, such as cholangiocarcinoma, 2-HG permits tolerance of wild-type p53 resulting in mutual exclusive patterns of IDH and p53 mutation. Inhibitors of mutant IDH have shown some therapeutic efficacy, although in the case of cholangiocarcinoma this has largely been limited to partial clinical response and disease stabilization (50). As the production of 2-HG can be effectively inhibited by well-tolerated agents in mutant IDH cholangiocarcinoma and other tumor types, our work suggests that combining inhibitors of mutant IDH with therapies that induce p53 accumulation could represent a strategy to reinvigorate p53-triggered anti-tumor effects, such as extrinsically and chemotherapy-triggered apoptosis, in p53 WT, IDH mutant cancers.

## MATERIALS AND METHODS

### Mouse modeling

All animal experiments were performed under approval of the Memorial Sloan Kettering or University Of North Carolina Institutional Animal Care and Use Committees. 8-10 weeks old C57Bl6N mice were utilized for hydrodynamic tail vein injections and as hosts for orthotopic transplant of primary mouse liver cancer cells as described below.

### Cell culture and treatments

NP^sh^-1, -2, and -3 primary cell lines were derived from liver tumors that formed via hydrodynamic tail vein injection (HTVI) of 8-10 week old C57BL/6N female mice maintained on doxycycline chow (625 mg/kg, Harland Teklad) with the following endotoxin free plasmids combined in a volume of sterile 0.9% NaCl solution corresponding with 10% of mouse weight in grams: (1) 5 μg pT3-EF1a- NrasG12D-IRES-rtTA, (2) 20 μg pT3-TRE-tRFP-shp53, and (3) 5 μg CMV-SB13, resulting in transposon mediated, stable integration of (1) and (2) into hepatocytes (as described in (26)). Mice were maintained on dox chow until tumor development. Tumors were harvested, minced, digested sequentially with 1mg/mL collagenase V (Sigma), 1mg/ml Dispase II (Roche), and in 0.25% trypsin (Gibco), with PBS washes in between each enzyme treatment. Dissociated cells were plated in complete DMEM (10% tetracycline free FBS (Gibco), 1x penicillin–streptomycin) containing 1 ug/mL doxycycline (to maintain expression of the tRFP linked shp53) on collagen-coated plates (PureCol, Advanced Biomatrix, 0.1 mg/ml). Primary cells were passaged until free from contaminating host cells (e.g. cancer associated fibroblasts) as indicated by depletion of tRFP negative cells by flow cytometry. NP^sh^ cells were pre-treated with AG-221 (10uM) for 7 days before beginning indicated experiments. RBE and SNU-1079 human cholangiocarcinoma cell lines (generous gift from S. Kugel Lab) were cultured in RPMI 1640 medium supplemented with 10% FBS (Gibco), 1x penicillin–streptomycin. 5 uM AG-120 (or DMSO) was added for at least 3 weeks as indicated before treatment with nutlin (5uM) or Adriamycin (indicated concentrations). All mouse and human cells were tested frequently for mycoplasma contamination using GeM Mycoplasma Detection Kit (Venor).

### Virus Production and Infection

Retrovirus was produced via co-transfection of cDNA or shRNA cargo with packaging plasmids pCMV-VSVG (Addgene #8454) and pBS-CMV-GagPol (Addgene #35614) into HEK293T cells. Lentivirus was produced via co-transfection of delivery cargo with pMD2.G (Addgene #12259) and psPAX2 (Addgene #12260). Virus containing medium was cleared of debris via centrifugation, filtered through 0.45 uM PES membrane, diluted 1:1 in appropriate culture medium, and mixed with 8 ug/mL polybrene. Cancer cell lines were exposed to virus for two 24- hour periods, grown for 24 hours in normal medium, then subjected to selection via antibiotics. Cells were selected for at least 7 days before beginning experiments. IDH2 cDNAs were cloned into retroviral backbone pMIG (Addgene #9044) as described (51). NP^sh^ cell lines were infected with empty pMIG vector, pMIG-IDH2^WT^, pMIG-IDH2^R140Q^, or pMIG-IDH2^R172K^ then sorted based on GFP expression (BD-FACS-ARIA) and cultured in complete DMEM supplemented with doxycycline. For CRISPR/Cas9-mediated genome editing, NP^sh^ cell lines were transduced to stably express Cas9 with the lentiCas9-blast (Addgene #52962) lentiviral backbone and selected with 10 ug/mL blasticidin. Single guide (sg)RNAs targeting CD95/Fas (sgFas- 1=ACTTCTACTGCGATTCTCC, sgFas-2=CACTTGGTATTCTGGGTCA) or an intergenic region of chromosome 8 (ACATTTCTTTCCCCACTGG) as control were introduced using pUSEPB (16) and selected with 4 ug/mL puromycin.

### Orthotopic tumor studies

For orthotopic transplant of NP^sh^ cells expressing indicated cDNAs or vectors enabling CRISPR/Cas9 editing cells maintained in doxycycline were washed with PBS, collected via trypsinization, and counted following quenching with complete DMEM. Cells were then resuspended in serum-free DMEM combined 1:1 with growth factor reduced matrigel (Corning) and 2.5×10^5^ cells injected into externalized livers of C57Bl6N using a Hamilton syringe fitted with a 31-gauge needle. Host mice were enrolled on dox chow (625 mg/kg, Harlan Laboratories) at least 2 days before transplant and mice were maintained on dox chow until randomized for p53 restoration by switching to conventional feed or maintenance on dox feed. Liver tumor volume was measured by small animal ultrasound (Vevo 2100) beginning 2 weeks post-transplant and mice were randomized into OFF DOX and ON DOX groups when tumors reached between ∼50- 400 mm^3^. Tumor volume was then measured every 5 days and animals were sacrificed at indicated timepoints, or in the case of longitudinal studies, when mice reached MSKCC and UNC IACUC approved humane endpoints for body condition or tumor burden. All mice bearing orthotopic tumors were evaluated daily for indications of distress or specified endpoint criteria. Specifically, mice were immediately euthanized if they showed signs of pain, difficulty breathing, infection, bleeding, weight loss >20% of starting weight, or developed tumors 15 mM in diameter. No tumors analyzed in this study exceeded the approved limit.

### Metabolite Profiling

Mouse and human cells were passaged 48 hours before collection for metabolite profiling under the indicated conditions at ∼70-80% confluency. Fresh medium was exchanged ∼16-18 hour before extraction. Metabolites extraction was performed in 1 ml ice-cold 80% methanol containing 2 μM deuterated 2-hydroxyglutarate (d-2-hydroxyglutaric-2,3,3,4,4-d5 acid, d5-2HG) as an internal standard. Cells in methanol were incubated overnight at −80 °C, and lysates were then scraped and centrifuged at 21,000xg for 20 min to remove insoluble, fixed material. Cleared metabolite extracts were then dried (Genevac EZ-2 Elite) and incubated in 50 μl of 40 mg/ml methoxyamine hydrochloride in pyridine at 30 °C for 2 h. Metabolites were then further derivatized by incubating for 30 minutes at 37 °C following addition of 80 μl MSTFA + 1% TCMS (Thermo Scientific) and 70 μl ethyl acetate (Sigma). Samples were analyzed with an Agilent 7890A gas chromatography system coupled to Agilent 5975C mass selective detector, operated in splitless mode with helium gas flow at 1 ml/min. 1ul of sample was injected onto an HP-5MS column and the gas chromatography oven temperature ramped from 60°C to 290°C over 25 minutes. Peaks representing compounds of interest were extracted and integrated using MassHunter software (Agilent Technologies), and peak area was normalized to the internal standard (d5-2HG) peak area and the protein content of duplicate samples as determined by BCA protein assay (Thermo Scientific). Ions used for quantification of metabolite levels are as follows: αKG, *m/z* 304; succinate, *m/z* 247; d5-2HG *m*/*z* 354; 2HG, *m*/*z* 349.

### Western Blotting

Tumor and cell lysates were extracted using RIPA Buffer (CST). Tumors were mechanically dissociated using Green RINO beads (NextAdvance, GREENR1-RNA) in a bullet blender tissue homogenizer (NextAdvance). Tumor and cell lysates were sonicated with a probe sonicator (Fischer scientific) then cleared via centrifugation at max speed. Protein concentrations were measured using the Pierce BCA Protein Assay (Thermo Fisher) read on a Cytation 5 (BioTek) and equal protein amounts were separated via SDS-PAGE. Proteins were transferred to 0.45 uM Immobilon-P PVDF membranes (Sigma-Aldrich), blocked with 5% milk or BSA in 1x Tris Buffered Saline (TBST), then incubated with primary antibodies in blocking solution overnight at 4°C on a platform rocker. Primary antibodies were used as follows: p53 (1C12) (1:1000, 2524S, Cell Signaling Technologies), Cleaved Caspase-3 (Asp175) (5A1E) (1:500, 9664, Cell Signaling Technologies), Fas (EPR24898-74) (1:1000, ab271016, Abcam), beta-Tubulin (1:5000, 66240-1- Ig, Proteintech), Vinculin (E1E9V) (1:3000, 13901S, Cell Signaling Technologies), IDH2 (1:1000, 15932-1-AP, Proteintech), Mdm2 (1:1000, 51541, Cell Signaling Technologies), p21 (F-5) (1:1000, sc-6246, Santa Cruz Biotechnology), Total Histone H3 (1B1B2) (1:3000, 14269, Cell Signaling Technologies), H3K9me2 (D85B4) (1:1000, 4658T, Cell Signaling Technologies), H3K9me3 (D4W1U) (1:1000, 13969T, Cell Signaling Technologies), H3K36me3 (D5A7) (1:500, 4909T, Cell Signaling Technologies), H3K27me3 (C36B11) (1:2000, 9733T, Cell Signaling Technologies), and H3K4me3 (C42D8) (1:1000, 9751S, Cell Signaling Technologies). After washing, corresponding horseradish peroxidase conjugated secondary antibodies (CST) were applied for one hour at 1:10,000 dilution and proteins were detected using Clarity Western ECL Substrate (BioRad). Secondary antibodies were goat anti-rabbit IgG HRP (1:10,000, 7074S, Cell Signaling Technologies) and horse anti-mouse IgG HRP (1:10,000, 7076S, Cell Signaling Technologies). Images were captured using a ChemiDoc Imaging System (BioRad).

### Histone Enrichment by Acid Extraction

First, cytoplasm was lysed by scraping cells into PBS then resuspending in Triton Extraction Buffer (TES) containing 0.5% Triton X-100 and 0.02% NaN_3_ in PBS supplemented with 1x complete Protease Inhibitor Cocktail tablet (Roche) on ice. Nuclei were isolated via centrifugation then washed once in TES. Nuclei were subjected to acid extraction in 0.2 N HCl overnight at 4°C. Insoluble debris were pelleted and the soluble histone fraction was collected and neutralized with NaOH. Protein concentration was determined using a BCA, and equal protein amounts were separated via SDS-PAGE for Western blotting as described above.

### Cytoplasm-Soluble Nucleoplasm-Chromatin Fractionation

The NE-PER nuclear and Cytoplasmic Extraction Reagent Kit (Thermo Fisher) was used to separate cytoplasm fraction from nuclear fraction according to manufacturer’s protocol with slight modifications. Following cytoplasmic extraction with the Cytoplasmic Extraction Reagent, nuclei were resuspended in 250 mM Sucrose, 10 mM MgCl_2_ containing 1X complete Protease Inhibitor Cocktail (Roche). Resuspended nuclei were floated on top of 350 mM Sucrose, 0.5 mM MgCl_2_ then centrifuged for 10 minutes at 1000xg to remove contaminating cytoplasmic debris. Nuclei were then washed with PBS, pelleted by centrifugation, and resuspended in Nuclear Extraction Reagent. After 30 minutes, the soluble nuclear fraction was isolated by pelleting the insoluble chromatin fraction at 21,000xg for 10 minutes. The insoluble chromatin fraction was washed once with Nuclear Extraction Reagent and resuspended in modified RIPA buffer containing 1.4% Triton-X100. Chromatin was sonicated to shear DNA releasing chromatin-bound proteins and Western blotting was performed as described above.

### qRT-PCR

RNA was extracted from cell lines using RNAeasy Mini Kit (Qiagen) using the manufacturer’s protocol. Reverse transcriptase (iScript, BioRad) synthesized cDNA from 1 ug of RNA measured on a NanoDrop One (Thermo Scientific). A CFX Opus Real-Time PCR System (Bio-Rad) was used for qPCR amplification with SYBR green (SsoAdvanced, Bio-Rad or PowerUp, Thermo Fisher). Primers were designed using MGH Harvard’s PrimerBank with 36B4 (Rplp0) used as a reference gene for mouse samples and beta-Actin (ACTB) for human samples. Mus musculus primer sequences were Fas/CD95: left, TATCAAGGAGGCCCATTTTGC; right, TGTTTCCACTTCTAAACCATGCT; Pmaip1/Noxa: left, GCAGAGCTACCACCTGAGTTC; right, CTTTTGCGACTTCCCAGGCA; Casp7: left, AAGACGGAGTTGACGCCAAG; right, CCGCAGAGGCATTTCTCTTC; Parp10: left, CTCTGTACTTTGAAAACCACCGT; right, GCTCAGTCGGACACCATGTA; Trp53: left, CTAGCATTCAGGCCCTCATC; right, TCCGACTGTGACTCCTCCAT; Cdkn1a: left, CGGTGTCAGAGTCTAGGGGA; right, ATCACCAGGATTGGACATGG; Mdm2: left, TGTCTGTGTCTACCGAGGGTG; right, TCCAACGGACTTTAACAACTTCA; Ddit4l: left, CGGCCAGCATTTCAGAGTTG; right, CAGGGACCAAGACCTTAGAGC; Ephx1: left, GGAGACCTTACCACTTGAAGATG; right, GCCCGGAACCTATCTATCCTCT; P2ry1: left, GAGGTGCCTTGGTCGGTTG; right, CGGCAGGTAGTAGAACTGGAA; 36B4: left, GCTCCAAGCAGATGCAGCA; right, CCGGATGTGAGGCAGCAG. Homo sapiens primer sequences were FAS/CD95: left, AGATTGTGTGATGAAGGACATGG; right, TGTTGCTGGTGAGTGTGCATT; TP53: left, CAGCACATGACGGAGGTTGT; right, TCATCCAAATACTCCACACGC; ACTB: left, CATGTACGTTGCTATCCAGGC; right, CTCCTTAATGTCACGCACGAT.

### Hematoxylin and Eosin (H&E) Staining

Tumors fixed in 10% zinc-buffered formalin overnight at 4°C were paraffin-embedded, sectioned at 5 uM, then deparaffinized and re-hydrated by xyline washes and progressive ethanol dilution series. After a 5-minute water wash, slides were incubated in 100% Mayer’s Hematoxylin Solution (Sigma) followed by 5 minutes in water. Slides were then incubated for 2 minutes in 0.2% Eosin Y (Sigma) in 70% ethanol containing 0.5% acetic acid and washed with water. Following a dehydration series in ethanol and xylenes, tissue was mounted using permount mounting medium (VWR).

### Immunofluorescence

Tumors were fixed overnight in 10% zinc-buffered formalin at 4°C, paraffin-embedded, and sectioned at 5 uM. Sections were deparaffinized and re-hydrated via xylene washes followed by incubation in progressively dilute ethanol. Antigen retrieval was performed in a pressure cooker at max pressure in citrate antigen retrieval buffer (Vector) before blocking in 5% BSA diluted in PBS. Primary antibodies were incubated overnight at 4°C with the following dilutions in blocking buffer: p53 (CM5) (1:300, NCL-L-p53-CM5p, Leica Biosystems), Cleaved Caspase-3 (Asp175) (5A1E) (1:400, 9664, Cell Signaling Technologies), Fas (EPR24898-74) (1:100, ab271016, Abcam), Green Fluorescent Protein (1:500, ab13970, Abcam), and Ki67 (SP6) (1:500, ab16667, Abcam). For immunofluorescence, fluorescently conjugated secondary antibodies were added for one hour at RT by diluting in blocking buffer as follows: Goat anti-Chicken IgY (H+L) Alexa Fluor 488 (1:500, A-11039, Invitrogen), Goat anti-Rabbit IgG (H+L) Alexa Fluor 594 (1:500, A11037, Invitrogen), and Goat anti-Rabbit IgG (Heavy chain) Alexa Fluor 555 (1:500, A27039, Invitrogen). Nuclei were stained with DAPI then images were captured using an FV1000 confocal laser scanning microscope (Olympus). For immunohistochemistry, endogenous peroxidases were quenched with 1% hydrogen peroxide in PBS following antigen retrieval. After blocking and primary antibody incubation, goat HRP anti-rabbit IgG peroxidase (#LS-J1066-50, Vector) was added for 1 hour at RT. DAB peroxidase (HRP) substrate was added (LS-J1075, Vector) for 60 seconds before quenching in PBS. Mayer’s hematoxylin solution (MHS32-1L, Sigma-Aldrich) was added to counterstain for 3 minutes then washed in water. Sections were then dehydrated in ethanol series, dipped in xylenes, then mounted under coverslips with VectaMount (Vector). Images were captured using an inverted microscope (Olympus IX70).

### Quantification of Ki-67 Immunofluorescence

An ImageJ Macro was used to process the images and measure the Ki-67 signal within the nuclei of GFP positive areas of five fields of view across three tumors of each condition. Briefly, a mask was generated from the GFP signal of each slice of tumor. Particle selection was performed on the DAPI staining within the mask followed by watershedding to outline individual nuclei within GFP positive region. The mean Ki-67 signal was measured for each nucleus. A Ki-67 positivity threshold was set from the mean nuclear Ki-67 signal and the average percent Ki-67 nuclei per field of view for each tumor is reported.

### p53 Activity Luciferase Reporter Assay

Lipofectamine 3000 Transfection Reagent (Thermo Fisher) was used to co-transfect NP^sh^ cells with PG13-luc (Addgene, #16442) or MG15-luc (Addgene, 16443) alongside pIS1 (Addgene, #12179) at a 2:1 ratio of Firefly to Renilla vector DNA according to manufacturer’s reverse transfection protocol in a 96-well black sided plate. Cells were transfected for 16 hours before incubating in normal complete DMEM supplemented with or without dox for an additional 48 hours. The Dual-Glo Luciferase Substrate (Promega) was added 1:1 with medium before reading firefly luminescence on a GloMax Luciferase Microplate Reader (Promega) with 0.5 second integration time. Dual-Glo Stop and Glo Substrate (Promega) was then added to measure Renilla luminescence. Separate wells containing no cells were used for background luminescence subtraction. Firefly luminescence was divided by Renilla luminescence to account for transfection efficiency and cell number within each well. Three experiments were performed each with triplicate wells of a 96-well plate.

### Caspase 3/7 Activity Luciferase Assays

NP^sh^ cells grown on dox or at indicated times after dox withdrawal were seeded into 96-well black sided plates with 6 wells per condition. Recombinant mouse Fas ligand (RnD Systems) was added to 100 ug/mL with HA Tag Antibody (RnD Systems) supplemented at 2.5 ng/mL for Fas crosslinking as recommended by manufacturer. Serum concentration was reduced to 1% in DMEM for all conditions for sensitization to Fas-mediated apoptosis. After 24 hours of FasL treatment, Caspase-Glo 3/7 Substrate (Promega) was added 1:1 to medium in triplicate wells and detected using a GloMax Luciferase Microplate Reader (Promega) with 0.5 second integration time. To normalize to cell number, triplicate wells were trypsinized, neutralized with complete medium, then counted using an Attune Nxt Autosampler (Thermo Fisher). Cell concentration was determined by gating all cells to exclude debris and triplicate concentrations were averaged to achieve the cells per well of each condition. Luminescence values were subtracted by background signal collected from wells devoid of cells then divided by the average cells per well to report the Caspase 3/7 Activity per cell for each condition. Caspase 3/7 activity in SNU-1079 and RBE cells was determined as above, except cells were cultured in 10% serum RPMI 1640 medium.

### CellTiter-Glo Cell Viability Assay

RBE and SNU-1079 were pretreated with 5 uM AG-120 for 3 weeks followed by seeding 3000 cells per well into a 96-well black sided plate in triplicate. The day after seeding, cells were treated with a serial dilution of Adriamycin for 72 hours. CellTiter-Glo substrate was added to triplicate wells of each condition at an equivalent volume of medium (75 uL) and luminescence was measured with a GloMax Luciferase Microplate Reader (Promega) with 0.5 second integration time. Luminescence was normalized to additional triplicate wells receiving no Adriamycin to account for differences in seeding between AG-120 and DMSO treated conditions.

### Growth Curves

Growth curves generated by measuring population doubling in NP^sh^-empty, NP^sh^-IDH2^WT^, NP^sh^- IDH2^R140Q^, and NP^sh^-IDH2^R172K^ cells over 9 days. Cells maintained on dox were washed with PBS, collected by trypsinization, counted, and 50,000 cells plated in triplicate in 6 well dishes with and without dox. At 2, 4, 6, and 9 days, cell number was determined by flow cytometry (Guava Technologies EasyCyteHT), at indicated timepoints and 50,000 cells were replated at day 2, 4, and 6. Population doublings for each growth period were calculated with the formula 3.32*(log(final cell number)-log(initial cell number)) and added to determine growth over time.

### Senescence Associated β-Galactosidase Assay

SA-β-galactosidase activity was detected in NP^sh^-empty, NP^sh^-IDH2^WT^, NP^sh^-IDH2^R140Q^, and NP^sh^- IDH2^R172K^ as described in (16). Cells were plated for 48 hours before staining at indicated times on and off dox. For staining, cells were washed with PBS, fixed with 0.5% glutaraldehyde in PBS for 15 min, washed with 1 mM MgCl_2_ in PBS, pH 5.5, and incubated overnight at 37°C in 1 mM MgCl_2_, 1 mg/mL X-Gal (Roche), and 5 mM each potassium ferricyanide (Sigma) and potassium ferrocyanide (Sigma) in PBS, pH 5.5. At least 200 cells were scored to determine SA-β- galactosidase activity from triplicate wells.

### BrdU staining

NP^sh^-empty, NP^sh^-IDH2^WT^, NP^sh^-IDH2^R140Q^, and NP^sh^-IDH2^R172K^ cells were grown for 48 hours at indicated conditions on and off dox and pulsed with 10um Brdu in complete medium for 1 hour. Cells were washed with PBS and then fixed in 4% paraformaldehyde in PBS and then treated with 1M HCl for 30 minutes at 37C to nick DNA. Cells were blocked with 5% BSA in PBS and BrdU was then detected with a mouse anti-Brdu primary antibody (BD Biosciences, Clone B44, 347580) and goat anti-mouse AlexaFluor-488 conjugated secondary (1:500, A-11001, Invitrogen) and nuclei counterstained with DAPI. At least 200 nuclei (DAPI positive) were scored for BrdU from wells in triplicate.

### CUT&RUN

Genomic occupancy of p53 was determined using the Cell Signaling Technology CUT&RUN Assay Kit according to manufacturer’s protocol. Briefly, TrypLE Express (Gibco) was used for cell dissociation followed by treatment with wash buffer and binding to Concanavalin A coated magnetic beads. Bead-coated cells were collected with a magnet and incubated in primary antibody targeting p53 (#53484, CST) or mouse IgG isotype control (#2524, CST) diluted 1:50 in antibody buffer with digitonin rocking overnight at 4°C. After washing in digitonin buffer, secondary anti-mouse antibody (ab46540, Abcam) was added to the bead slurry at 1:100 rocking at 4°C for 1 hour. Protein A/G Micrococcal Nuclease (CST) was added 1:30 for 1 hour on ice followed by chromatin digestion by adding CaCl_2_ for 30 minutes at 4°C. Chromatin fragment retrieval was performed by adding Stop buffer (CST) and incubating for 30 minutes at 37°C. DNA fragments were purified with DNA Purification Buffers and Spin Columns (CST) according to manufacturer’s protocol. Libraries were prepared with the Kapa Hyperprep Kit (Roche) using half reaction volumes suggested by manufacturer’s protocol. DNA concentrations were determined using a Qubit (Invitrogen) and quality was confirmed with a TapeStation (Agilent). Pooled libraries were sequenced on a Nextseq 500 with a 75-cycle high output kit, paired end (Illumina). CUT&RUN was performed in triplicate in independently processed samples.

### CUT&RUN Data Analysis

Raw fastq files were trimmed to remove Ns using TrimGalore (0.6.7) (Krueger F-TrimGalore) using the following options -j --fastqc --paired --gzip --basename. Trimmed reads were aligned to mouse mm10 genome using bowtie2 (2.3.4.1) (64)--very-sensitive-local -X 800. Samtools (65) was used to generate aligned sorted BAM files and reads with mapping quality (mapq) scores < 10 were filtered out. TMM on high-abundance regions (peaks) was performed using the csaw package (66) by generating scale factors with effective genome size of 2407883318 and flagging --peaks --winSize=200 --maxFrag=500 --pe=both to normalize for binding efficiency and technical variation between samples. Scale factors were used to calculate effective library size, divided by a million, and the reciprocal was supplied to --scaleFactor in csaw generating bigwig files. Triplicate bigwig files were then averaged using wiggletools (67) mean for visualization in Integrative Genomics Viewer (IGV) (68). Peak calling was performed with MACS2 (2.2.7.1) (69) -q 0.05 -g 2407883318 using the control mouse IgG sample for background subtraction and blacklisted peaks were removed with bedtools. The R package DiffBind (Stark--Bioconductor) was used for differential binding analysis by counting reads with 250 bp summits, generating a consensus peak set using default parameters, normalizing with contrast, normalize=DBA_NORM_NATIVE, background=FALSE, and using the default DESeq2 parameters to identify differentially bound peaks. Heatmaps were generated using DeepTools computeMatrix and plotHeatmap with the consensus peaks flanked by 5000 bp on either side. Peaks were annotated to the nearest gene according to GRCm38v100.

### RNA-Seq analysis

Total RNA was extracted from mouse (NP^sh^-1-IDH2^WT^, NP^sh^-1-IDH2^R172K^) and human cells (RBE) with the RNAeasy extraction kit (Qiagen). Bulk RNA-SEQ of NP^sh^-1-IDH2^WT^, NP^sh^-1-IDH2^R172K^ cells was performed by Admera. Briefly, RNA sample quality was assessed by RNA Tapestation (Agilent Technologies Inc., California, USA) and quantified by Qubit 2.0 RNA HS assay (ThermoFisher, Massachusetts, USA). Paramagnetic beads coupled with oligo d(T)25 were combined with total RNA to isolate poly(A)+ transcripts based on NEBNext® Poly(A) mRNA Magnetic Isolation Module manual (New England BioLabs Inc., Massachusetts, USA). Prior to first strand synthesis, samples are randomly primed (5’ d(N6) 3’ [N=A,C,G,T]) and fragmented based on manufacturer’s recommendations. The first strand is synthesized with the Protoscript II Reverse Transcriptase with a longer extension period, approximately 30 minutes at 420C. All remaining steps for library construction were used according to the NEBNext® UltraTM II Directional RNA Library Prep Kit for Illumina® (New England BioLabs Inc., Massachusetts, USA). Final libraries quantity was assessed by Qubit 2.0 (ThermoFisher, Massachusetts, USA) and quality was assessed by TapeStation HSD1000 ScreenTape (Agilent Technologies Inc., California, USA). Average final library size was about 400bp with an insert size of about 250bp. Illumina® 8-nt dual-indices were used. Equimolar pooling of libraries was performed based on QC values and sequenced on an Illumina® HiSeq (Illumina, California, USA) with a read length configuration of 75 SE for 25M SE reads per sample.

RNA-sequencing analysis of NP^sh^ cells expressing an IDH allelic series (i.e. NP^sh^-1-IDH2^WT^, NP^sh^- 1-IDH2^R172K^) to determine the effect of mutant IDH on p53 dependent gene expression through 2- HG was performed as follows. Alignment of fastq files to the GRCm38v100 version of the mouse genome and transcriptome was performed with STAR 2.7.6a (70). Gene abundance for each sample was quantified with Salmon v1.4.0. (71). DESeq2 1.22.2 Bioconductor Package was used to normalize and perform differential gene expression analysis on all features with greater than 10 reads in at least one sample using R 4.0.3 (72,73). Multiple hypothesis testing was corrected by using the false discovery rate and genes were considered differentially expressed with an FDR corrected p-value ≤ 0.05 and a fold change ≥ 2 or ≤ -2. Rescued genes with AG-221 were non-differentially expressed with threshold of FDR corrected p-value > 0.05.

For gene set enrichment analysis of differential expression in NP^sh^-1-IDH2^WT^, NP^sh^-1-IDH2^R172K^ cells RNA-SEQ data was analyzed as follows. Approximately 20 million 150-bp, paired-end reads were retrieved per replicate condition. Resulting RNA-seq data were analyzed by removing adapter sequences using Trimmomatic v0.36 (74), aligning sequencing data to GRCm38-mm10 with STAR v2.7.11a (70), and quantifying genome-wide transcript count using featureCounts v1.6.3 (75) to generate raw count matrix. Differential gene expression analysis was performed using the DESeq2 v1.28.1 package (72) between experimental conditions, using 3 independent biological replicates per condition, implemented in R (http://cran.r-project.org/). DEGs were determined by > 2-fold change in gene expression with adjusted P < 0.05.

Bulk RNA-seq of RBE cells was performed by Novogene on the Illumina NovaSeq PE150 platform utilizing paired-end sequencing approach with 150-bp read length. RNA-sequencing analysis of RBE cells was performed via the Partek™ Flow™ analysis software version 10.0.23.0720. (Partek^TM^ Flow^TM^ REF) Raw reads were trimmed to exclude reads with quality score < 20. STAR – 2.7.8a (23104886) was used to align reads to the hg19 genome using default parameters. Aligned reads were counted using HTSeq 0.11.0 (76) followed by normalization and differential expression analysis using DESeq2 excluding features with an average coverage of less than 1.

### Gene Set Enrichment Analysis (GSEA)

GSEA analysis of NPsh-1-IDH2^WT^ versus NPsh-1-IDH2^R172K^ cells was performed as follows. Gene set enrichment analysis (GSEA v4.1.0) was performed using the GSEAPreranked tool for signatures in the Molecular Signatures Database (MSigDBv7.2; http://software.broadinstitute.org/gsea/msigdb). The metric scores were calculated using the sign of the fold change multiplied by the inverse of the p-value. GSEA between RBE cells treated with 5 uM Nutlin+Vehicle or 5 uM Nutlin+ 5 uM AG-120 was performed as follows. Genes from counts sheets containing transcripts per million were collapsed to remap to Human Gene Symbols (MSigDB.v2023.2.Hs.chip) and ranked with default Signal2Noise parameters in GSEA (V4.3.2). 1000 gene set permutation were used to query the human hallmarks gene sets collection (h.all.v2023.2.Hs.symbols.gmt) or a curated list of p53 target gene sets (Supplementary Table 2).

### ATAC-Seq

ATAC-Seq was performed according to the OMNI-ATAC protocol (28846090). Briefly, 75,000 viable NP^sh^-1-IDH2^WT^ or NP^sh^-1-IDH2^R172K^ cells under indicated conditions (ON/OFF dox, AG-221) were sorted (Sony MA900) and washed with ice cold PBS. Pelleted cells were then washed in 1 ml of cold ATAC-seq resuspension buffer (i.e. RSB: 10 mM Tris-HCl pH 7.4, 10 mM NaCl, and 3 mM MgCl_2_), and lysed in 50 μl of RSB supplemented with 0.1% NP-40, 0.1% Tween-20, and 0.01% digitonin on ice for 3 minutes. This raw nuclei suspension was then washed with RSB containing only 0.1% Tween-20 and pelleted nuclei were subject to tagmention in 50ul of transposition mix (25 ul 2X TD buffer, 100 nM TDE enzyme (Illumina), 16.5 ul PBS, 0.5ul 1% digitonin, 0.5ul 10% Tween-20, 5ul H2O) for 30 minutes at 37C with shaking at 1000 rpm in a thermomixer. Tagmented DNA was column purified (Qiagen MinElute). Tagmentated DNA was amplified with barcode primers. Library quality and quantity were assessed with Qubit 2.0 DNA HS Assay (ThermoFisher, Massachusetts, USA), Tapestation High Sensitivity D1000 Assay (Agilent Technologies, California, USA), and QuantStudio® 5 System (Applied Biosystems, California, USA). Equimolar pooling of libraries was performed based on QC values and sequenced on an Illumina® HiSeq (Illumina, California, USA) with a read length configuration of 150 PE for 50M PE reads (25M in each direction) per sample.

### ATAC-Seq analysis

The ENCODE consortium ATAC-Seq workflow was used to process ATAC-Seq data (77). Bowtie2 was used to align reads to GRCm38 (64), duplicate reads were removed, and MACS2 was used to call peaks (69). Overlapping peaks were merged to generate a consensus peak list and peaks with fewer than 10 reads in at least sample were removed. Peaks were annotated to the nearest gene in the mouse transcriptome version VRCm38v100. The number of reads under each peak was used as the raw chromatin accessibility value and DESeq2 (72) was used for normalization and differential openness analysis using R (73). To correct for multiple hypothesis testing, false discovery rate p-values were used.

### Statistical analysis and quantification

Statistical tests used for each experiment are described in accompanying figure legend and were performed with GraphPad Prism 10. Unless noted, threshold for significance is p<0.05. Sample number (n) for all experiments is noted in figure legends and represents biological replicates as described.

### Figure preparation

Illustrations were generated with Adobe Illustrator 2024 or BioRender.com where noted.

### Data availability

CUT and RUN, RNA-SEQ, and ATAC-SEQ data from this study will be made available in the Gene Expression Omnibus

## Author’s Contributions

**C. Martin:** Conceptualization, data curation, formal analysis, methodology, validation, investigation, visualization, writing—original draft, writing—review and editing. **W.B. Sullivan:** Investigation, visualization, validation, writing—review and editing. **J. Brinkman:** Investigation, formal analysis, validation. **D. Scoville:** Investigation, methodology. **J.J. Yashinskie:** Investigation, validation, writing—review and editing. **S. Tian:** Investigation, validation. **R.E. Mezzadra:** Investigation, validation, methodology. **Y.J. Ho:** Formal analysis, methodology, visualization. **R.P. Koche:** Formal analysis, methodology, validation. **T. Baslan:** Methodology, validation, writing—review and editing. **J. Raab:** Methodology, writing—review and editing. **D. Corcoran:** Formal analysis, data curation, methodology, visualization, writing—review and editing. **L.W.S. Finley:** Methodology, resources, writing—review and editing. **S.W. Lowe:** Methodology, resources, writing—review and editing. **J.P. Morris IV:** Conceptualization, investigation, resources, data curation, formal analysis, supervision, funding acquisition, visualization, methodology, project administration, writing—original draft, writing—review and editing.

## Acknowledgements

We thank J. Simon and W. Luan for technical support with animal experiments and members of the Morris, Finley, Raab, and Lowe laboratories for advice and discussion. RBE and SNU-1079 cells were a kind gift from S. Kugel. J.J.Y. was supported by T32HD060600. C. Martin was supported by NIGMS 5T32GM119999. J. Raab was supported by R35GM147286. J. Brinkman was supported by 5T32GM135128-03. J.P.M.IV was supported by a Pancreatic Cancer Action Network Career Development Award.

---

**Supplemental Figure 1.**
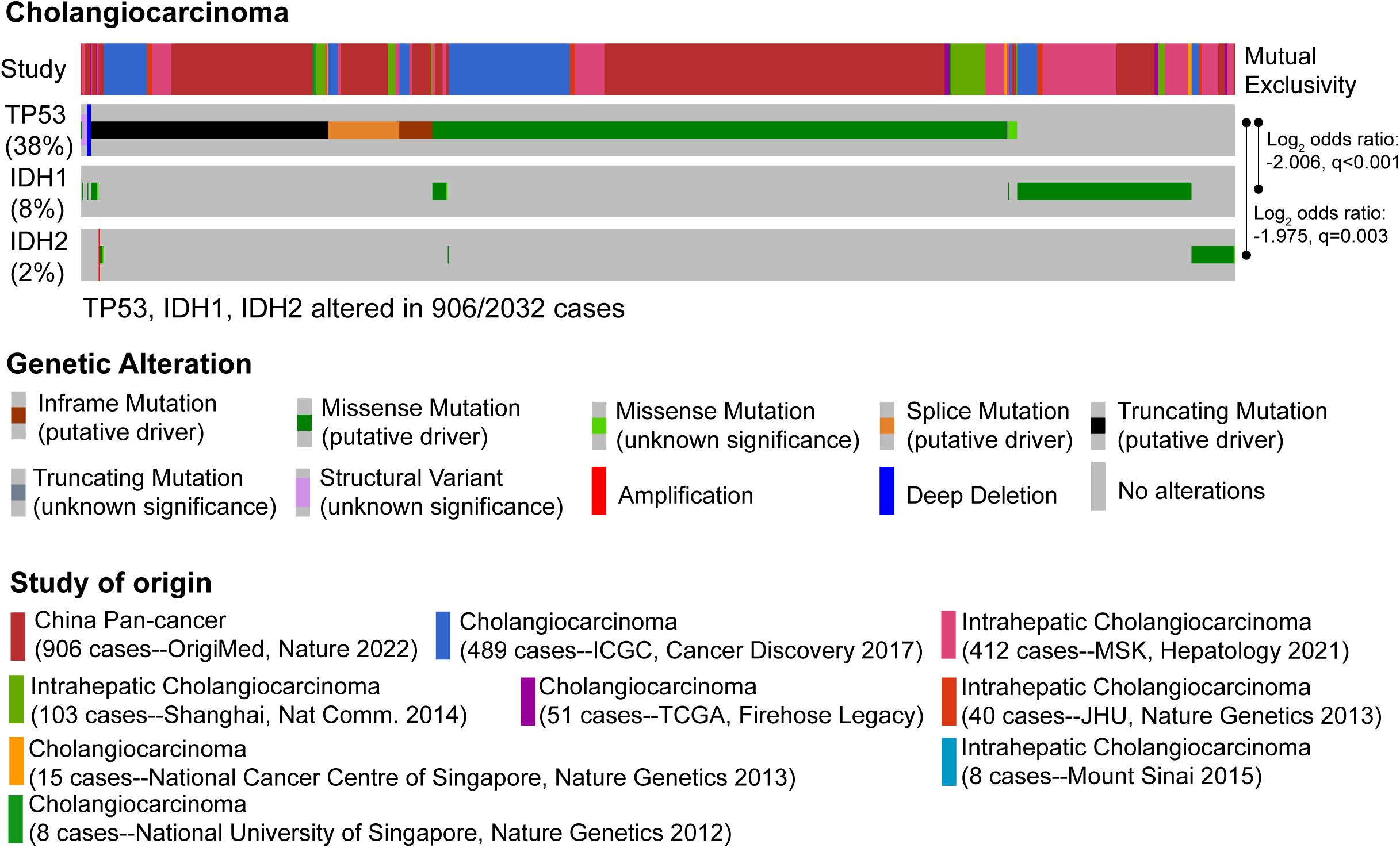
TP53 and IDH1/2 mutations are mutually exclusive in cholangiocarcinoma. Data from http://cbioportal.org

**Supplemental Figure 2.**
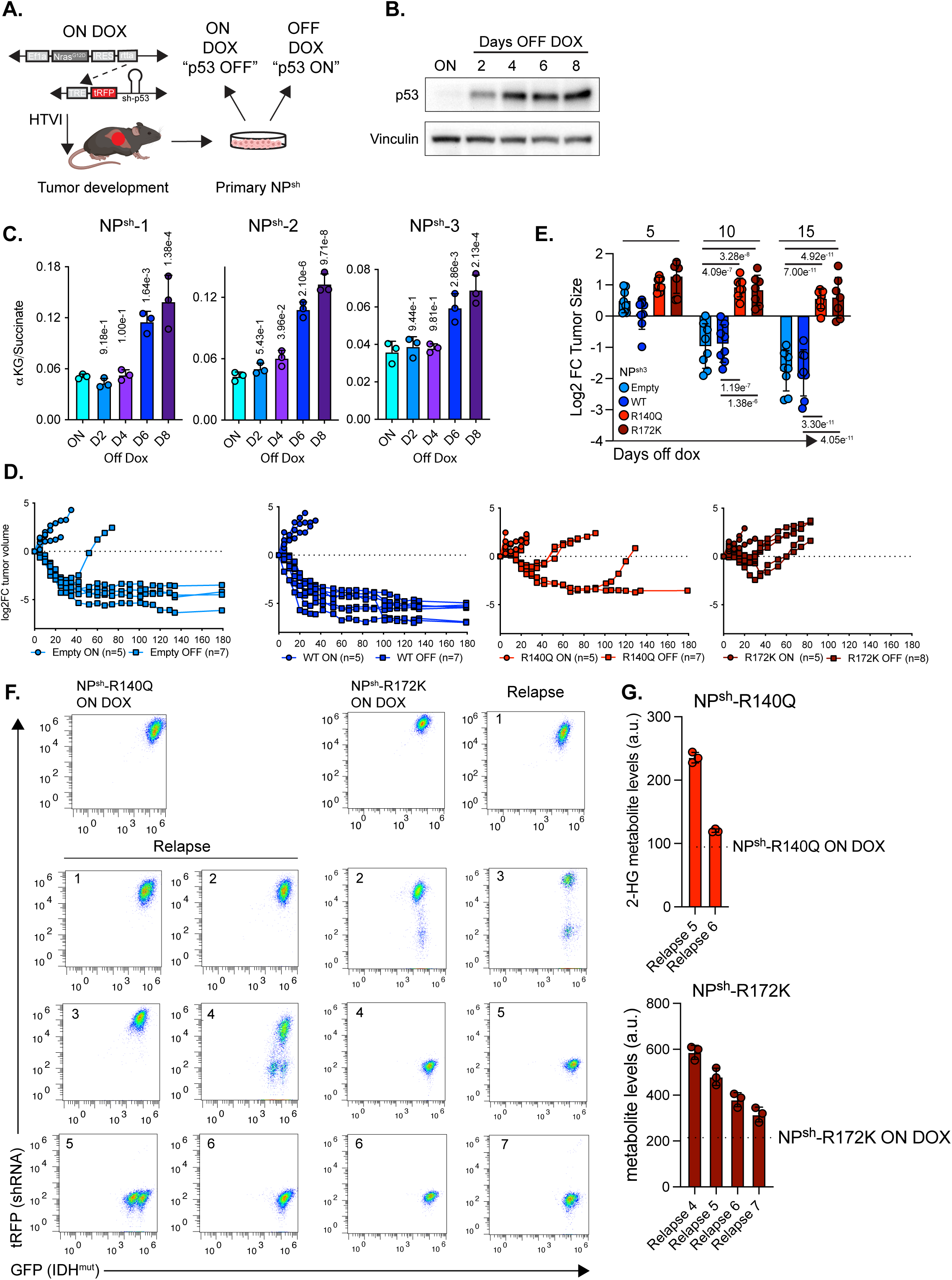
Cancer associated IDH mutants prevent p53 triggered tumor regressions and promote tumor relapse that tolerates active p53. **a.** Schematic of generation and p53 restoration of NP^sh^ liver cancer cell lines. Mouse image from Biorender.com. **b**. Western blot of p53 at indicated times following dox withdrawal in NP^sh^-1 cells. **c**. GC-MS measurement of cellular αKG:succinate ratio in 3 NP^sh^ cell lines at indicated times after dox withdrawal. **d**. Longitudinal, relative volume of NP^sh^-1 empty vector, IDH2^WT^, IDH2^R140Q^, or IDH2^R172K^ orthotopic tumors on dox and following withdrawal of dox chow. **e**. Relative volume of indicated NP^sh^-3 empty vector, IDH2^WT^, IDH2^R140Q^, or IDH2^R172K^ orthotopic tumors following withdrawal of dox chow. **f**. Flow cytometry analysis of primary cell lines generated from tumors arising through orthotopic transplant of NP^sh^-1-IDH2^R140Q^ cells (left) or NP^sh^-1-IDH2^R172K^ (right) into mice kept on dox and after relapse off dox. **g**. GC-MS measurement of 2-HG in NP^sh^-1-IDH2^R140Q^ and NP^sh^-1-IDH2^R172K^ relapse tumors progressing without reactivation of the p53 shRNA. Dotted line indicates average 2-HG level in parent lines from Figure 1B. c, e, significance calculated with one way ANOVA with Tukey correction for multiple comparisons.

**Supplemental Figure 3.**
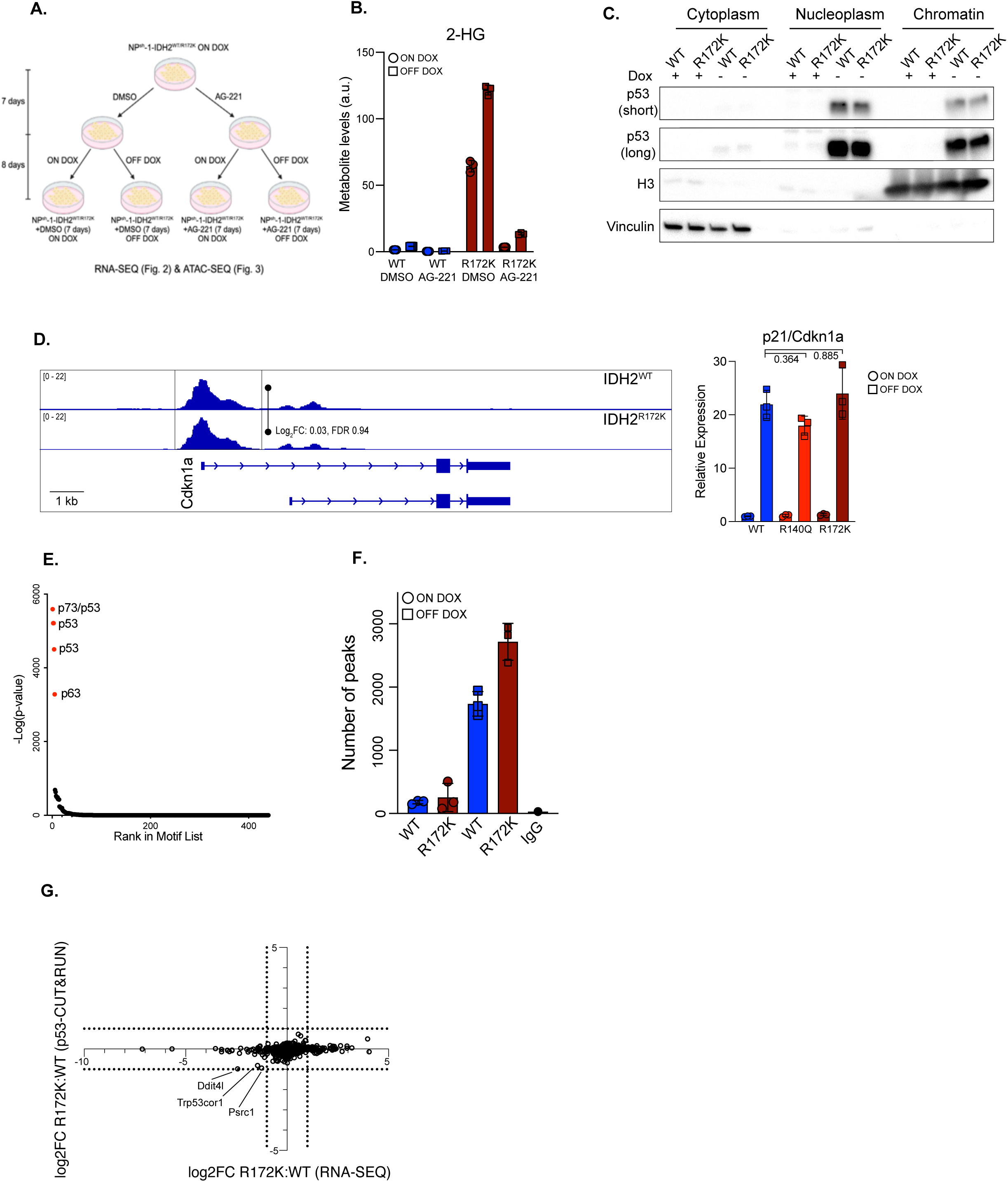
Mutant IDH uncouples p53 from gene regulation without preventing p53 accumulation, or DNA binding. **a.** Schematic of RNA-SEQ (Figure 2) and ATAC-SEQ (Figure 3) analysis of NP^sh^-1-IDH2^WT^ and NP^sh^-1-IDH2^R172K^ cells pre-treated with 10 uM AG-221 before maintenance on dox or p53 restoration (8 days). Illustration generated with BioRender.com. **b**. GC-MS measurement of cellular 2-HG levels of NP^sh^-1-IDH2^WT^ and NP^sh^-1-IDH2^R172K^ cells treated on and off dox (8 days) with AG-221 as described in **a** (n=3 wells per condition). **c**. p53 western blot in indicated fractions of NP^sh^-1-IDH2^WT^ and NP^sh^-1-IDH2^R172K^ cells following dox withdrawal. **d**. Left, average p53 CUT&RUN (n=3) profiles of the Cdkn1a locus in NP^sh^-1-IDH2^WT^ and NP^sh^-1-IDH2^R172K^ cells off dox for 8 days. Right, qRT-PCR of Cdkn1a in NP^sh^-1-IDH2^WT^, IDH2^R140Q^, and IDH2^R172K^ cells on and off dox for 8 days normalized to 36b4. **e**. Enrichment plot of known motifs identified by HOMER analysis in p53 peaks identified by CUT&RUN. **f.** Number of p53 peaks identified in NP^sh^-1-IDH2^WT^ and NP^sh^-1-IDH2^R172K^ cells on dox and 8 days off dox using MACS2 (FDR<0.05, n=3 per condition, n=1 for IgG). **g**. Correlation of average differential gene expression versus peak magnitude between direct p53 targets in off dox NP^sh^-1-IDH2^WT^ versus NP^sh^-1-IDH2^R172K^ cells. d, significance calculated with one way ANOVA with Tukey correction for multiple comparisons.

**Supplemental Figure 4.**
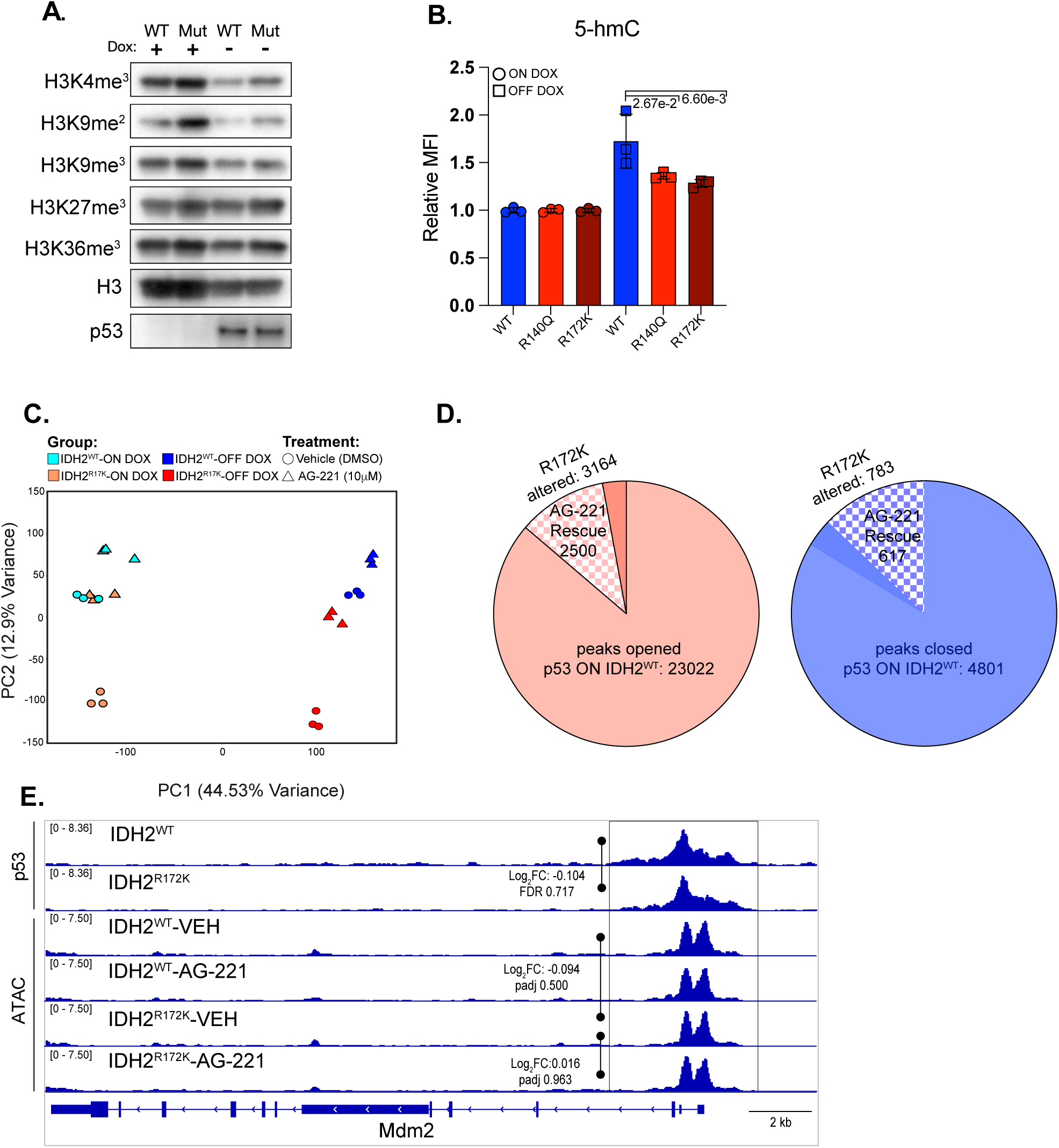
Mutant IDH perturbs αKG dependent chromatin remodeling that links p53 activity to gene expression. **a.** Western blot of H3K4me3, H3K9me2, H3K9me3, H3K27me3, H3K36me3, and p53 in NP^sh^-1-IDH2^WT^ and NP^sh^-1-IDH2^R172K^ cells on and 8 days off dox. **b**. Flow cytometry analysis of 5-hmC (median fluorescence intensity, MFI) in NP^sh^-1-IDH2^WT^, IDH2^R140Q^, and IDH2^R172K^ cells on and 8 days off dox. **c**. Principal component analysis of ATAC-SEQ of NP^sh^-1-IDH2^WT^ and NP^sh^-1- IDH2^R172K^ cells following p53 restoration (on dox versus 8 days off dox) treated with vehicle or AG-221. **d**. Quantification of p53 opened and closed peaks, differentially opened in the setting of mutant IDH and rescued by inhibition of 2-HG. **e**. Average p53 CUT&RUN and ATAC-SEQ (n=3) profiles of the Mdm2 locus in NP^sh^-1-IDH2^WT^ versus NP^sh^-1-IDH2^R172K^ cells 8 days after p53 restoration.

**Supplemental Figure 5.**
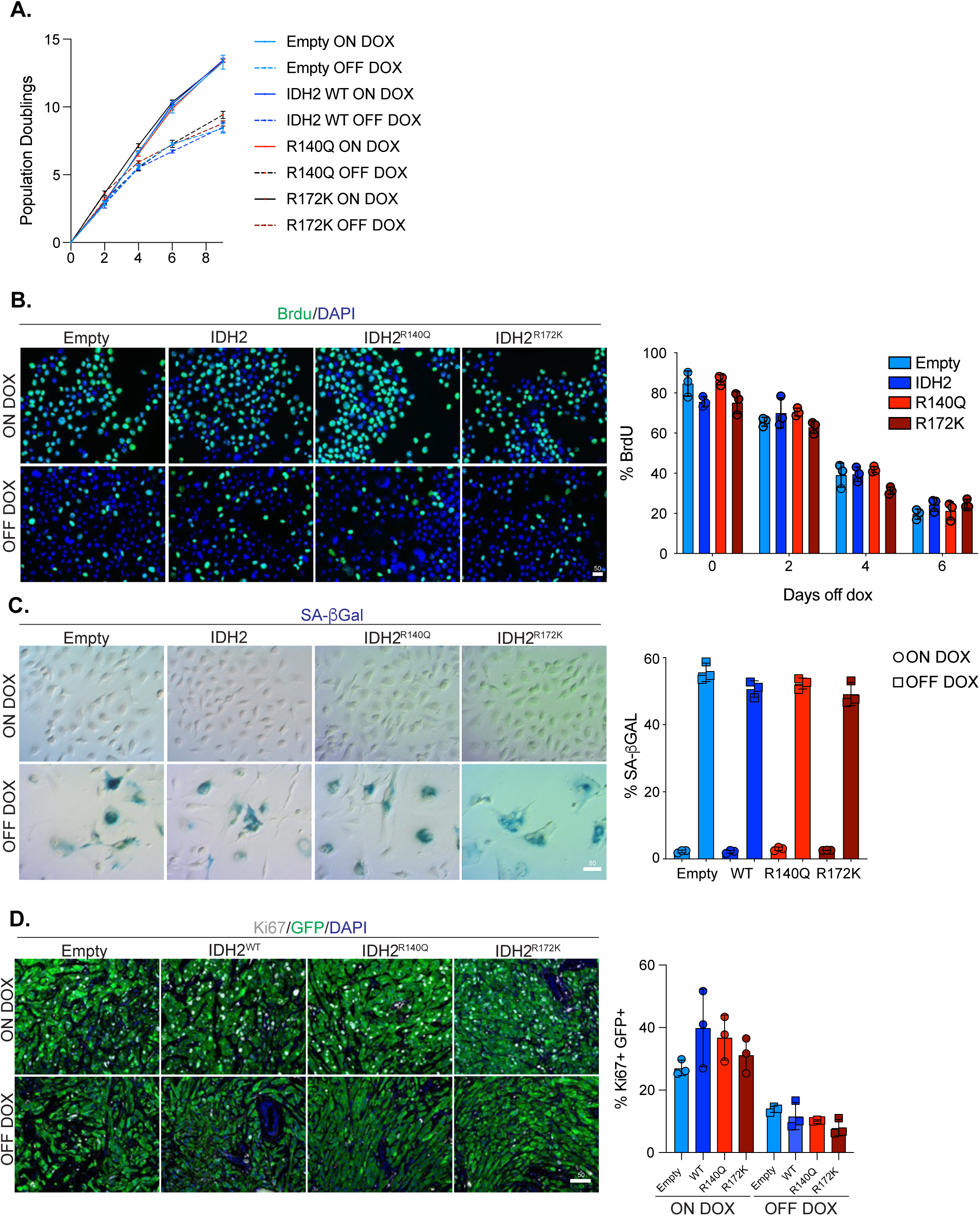
Mutant IDH does not prevent p53 induced arrest or senescence. **a.** Population doubling curve of NP^sh^-1 empty vector, IDH2^WT^, IDH2^R140Q^, or IDH2^R172K^ cells on and off dox. **b**. Left, representative Brdu staining in NP^sh^-1 empty vector, IDH2^WT^, IDH2^R140Q^, or IDH2^R172K^ cells on dox or 6 days after dox withdrawal. Right, quantification of Brdu staining in NP^sh^-1 empty vector, IDH2^WT^, IDH2^R140Q^, or IDH2^R172K^ cells on dox or 2, 4, and 6 days after dox withdrawal. **c**. Left, representative senescence associated beta galactosidase (SA-βgal) staining in NP^sh^-1 empty vector, IDH2^WT^, IDH2^R140Q^, or IDH2^R172K^ cells on dox or 8 days after dox withdrawal. Right, quantification of SA-βgal in NP^sh^-1 empty vector, IDH2^WT^, IDH2^R140Q^, or IDH2^R172K^ cells on dox and 8 days after withdrawal. **d**. Left, representative co-immunofluorescence of Ki67 and GFP in NP^sh^-1 empty vector, IDH2^WT^, IDH2^R140Q^, or IDH2^R172K^ orthotopic tumors in dox fed mice and in mice 10 days after dox withdrawal. Right, quantification of percent double Ki67/GFP positive cells. Scale bars b,c,d 50µM.

**Supplemental Figure 6.**
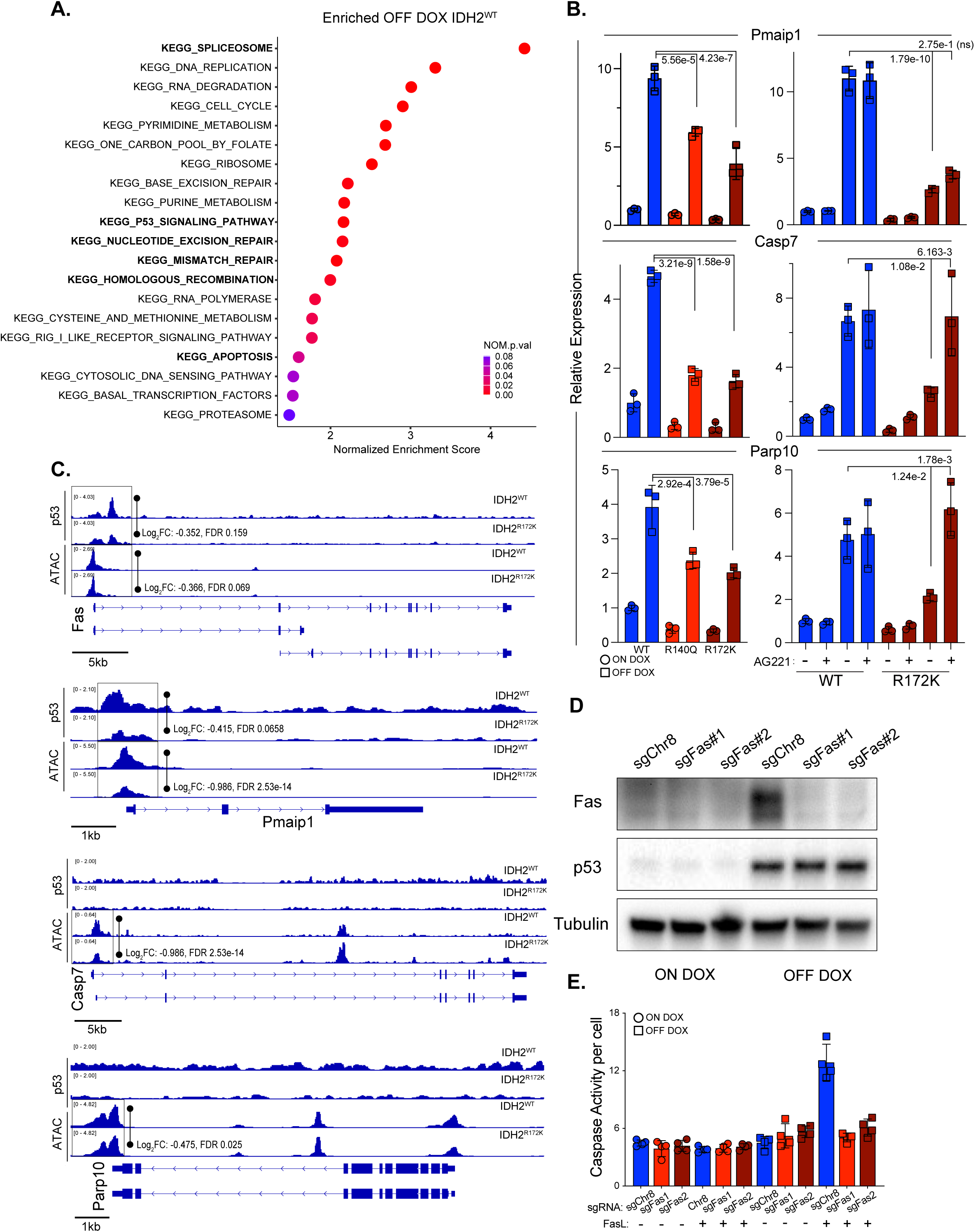
Mutant IDH interferes with expression of pro-apoptotic p53 targets. **a.** Hallmark GSEA terms of RNA-SEQ enriched in NP^sh^-1-IDH2^WT^ over NP^sh^-1-IDH2^R172K^ cells 8 days off dox. Bolded terms have been implicated in tumor suppression by p53. **b**. Left, qRT-PCR of Pmaip1/Noxa, Casp7, and Parp10 in NP^sh^-1- IDH2^WT^, IDH2^R140Q^, and IDH2^R172K^ cells on and 8 days off dox normalized to 36b4. Right, qRT-PCR of Pmaip1/Noxa, Casp7, and Parp10 in NP^sh^- 1-IDH2^WT^ and NP^sh^-1-IDH2^R172K^ cells on and 8 days off dox treated with AG-221 normalized to 36b4. **c**. Average p53 CUT&RUN and ATAC-SEQ (n=3 each condition) profiles of the Fas, Pmaip1, Casp7, and Parp10 loci in NP^sh^-1-IDH2^WT^ versus NP^sh^-1-IDH2^R172K^ cells 8 days after p53 restoration **d**. Western blot of Fas and p53 in NP^sh^-1 cells expressing sgChr8, sgFas-1, and sgFas-2 on dox and 8 days off dox. **e**. Cleaved caspase 3/7 activity measured by Caspase-Glo 3/7 luminescence assay in NP^sh^-1 cells expressing sgChr8, sgFas-1, and sgFas-2 on dox and 8 days off dox treated with FasL or vehicle.

**Supplemental Figure 7.**
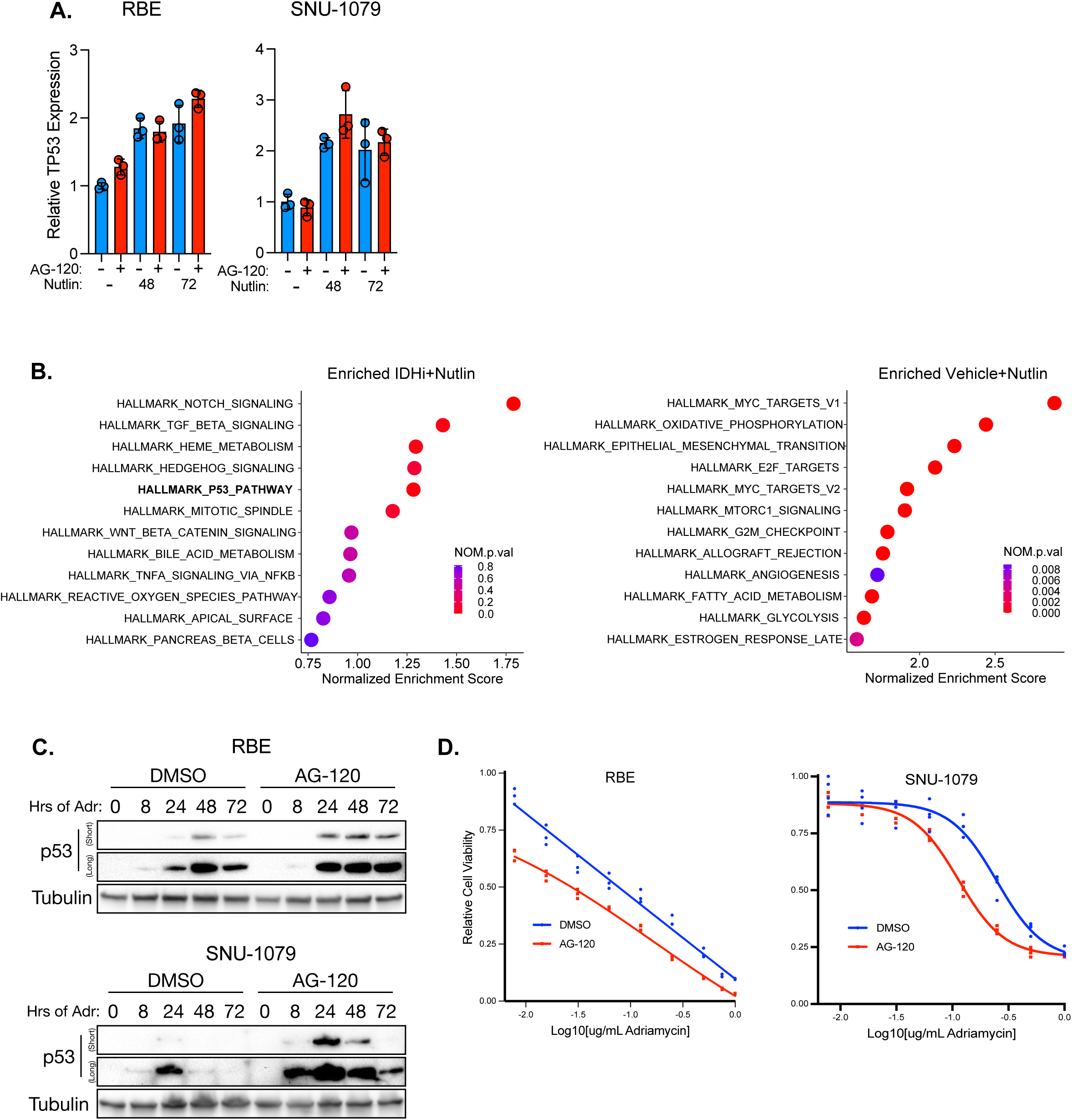
Inhibiting mutant IDH alters the response to p53 accumulation in cholangiocarcinoma cells. **a.** qRT-PCR of TP53 in RBE (left) and SNU-1079 (right) pre-treated with AG-120 or DMSO and 5 uM nutlin for 48 and 72 hours normalized to ACTB (n=3 wells per condition). **b**. Normalized enrichment score of GSEA human hallmark gene sets in 5 uM nutlin treated RBE cells enriched in cells pretreated with 5 uM AG-120 (left) or vehicle (right). **c**. Western blot of p53 in RBE (top) and SNU-1079 (bottom) pre-treated with AG-120 or DMSO and given 500ng/ml Adriamycin (Adr.) at indicated time points. **d**. RBE (left) and SNU-1079 (right) were treated with indicated concentrations of adriamycin and viability was measured with CellTiter-Glo. Luminescence was normalized to no adriamycin condition to account for differences in seeding between AG-120 and DMSO pretreated cells.

## Notes

### Competing Interest Statement

While not directly related to this manuscript, S.W.L. declares competing interests outside consultancy and equity for Oric Pharmaceuticals, Blueprint Medicines, Mirimus, Senecea Therapeutics, Faeth Therapeutics and PMV Pharmaceuticals; and outside consultancy (no equity) for Fate Therapeutics. While not related to this manuscript R.P.K. is a co-founder of and consultant for Econic Biosciences. All other authors declare no conflicts of interest.

