## Supplementary Table 1 for "Mutant IDH uncouples p53 from target gene regulation to disable tumor suppression"

| p53 Up and Mutant IDH Down | p53 Up and Mutant IDH Up | p53 Down and Mutant IDH Up | p53 Down and Mutant IDH Down |
|---|---|---|---|
| Ephx1 | Prl2c3 | Id1 | Gnai1 |
| Egfr | Wfs1 | Palm3 | Trim47 |
| Inhba | Aldh1a1 | Serpina1b | Kif21b |
| Lsr | Ivl | Apoc2 | Eprn |
| Arap3 | H2ac19 | Id2 | Ffar4 |
| Bcl2l15 | Il1rn | Itga7 | Slc7a8 |
| Psrc1 | Mcpt8 | Epb41l4b | Adamts1 |
| Itgb4 | Plxnd1 | Id3 | Fgfr4 |
| Acan | Gm20634 | Adora1 | Tgfb2 |
| Glp1r | Upk1b | Pcdhac2 | |
| Myo5c | Dhrs3 | Rtn4rl1 | |
| Slfn2 | Krt7 | Erbb3 | |
| Atf5 | Adssl1 | Krt6a | |
| Htra1 | Gm49774 | Sox11 | |
| Pros1 | Gm5662 | Maf | |
| Efemp2 | Vip | Nhsl2 | |
| Nes | Snx20 | Tgfbr3 | |
| Ildr2 | Sec16b | Msx1 | |
| Stard5 | Lilr4b | Slco5a1 | |
| Lad1 | 1110002E22F | Tubb3 | |
| Rasgrf2 | Slc35f1 | Gpc6 | |
| Ripor2 | Mturn | Aqp1 | |
| Piezo2 | Sorcs2 | Sprr1b | |
| Cfap69 | Nova2 | Vgf | |
| Hexa | Gm21939 | Fhl1 | |
| Zbtb4 | Sardh | Ceacam1 | |
| Tceal9 | Plin4 | Phospho1 | |
| Vcan | H3c15 | Sprr2a2 | |
| Anxa13 | Usp17ld | Gngt2 | |
| Fcho1 | Nrcam | Clca3b | |
| Irgm1 | Tdpoz8 | Gsta1 | |
| Nkx2-3 | Nbl1 | Naip5 | |
| Pmaip1 | 6330403L08F | Sult1d1 | |
| Tceal8 | Atp1a3 | Rpl7a-ps11 | |
| Acsl6 | Lamc3 | Atoh8 | |
| Sema5a | Ccl7 | Pax8 | |
| Sgce | Ces2f | Gm42793 | |

| | | |
|---|---|---|
| Synpr | Lrrc66 | Habp2 |
| Prrx1 | Serpinb11 | Fbn1 |
| Atp6v0e2 | Cp | Mal |
| Tcf24 | Mx1 | Gm3776 |
| Ddit4l | Plb1 | Glp2r |
| Map1lc3a | Ces2e | Cd209e |
| Scamp5 | Lce1g | Tmem229b |
| Kcnab1 | Ampd3 | Hsd17b14 |
| P2ry1 | Ly6f | Gm2694 |
| Klhl1 | Gm36660 | Abcc2 |
| Cnp | Olfr1318 | Glyatl3 |
| Npr2 | Trdc | Rab39b |
| 2210016F16F | Xlr | Tomt |
| Unc5c | Abca4 | Muc13 |
| Gm9926 | Nckap1l | Chrna2 |
| Apln | Gm5543 | D6Ertd527e |
| Dsg2 | R3hdml | Rimbp3 |
| Cdca7l | Gm9121 | Ppfia3 |
| Shroom1 | Tmem184a | Sptssb |
| Pcdhgb6 | Cercam | Cyp2d26 |
| 9930111J21R | Trdv4 | 5033430I15Rik |
| Palmd | Tnip3 | Foxa3 |
| Tmem181a | Espnl | Slc6a19 |
| Dzip1 | Noxa1 | Pou4f1 |
| Smarca2 | Gm12620 | Zan |
| Hoxb8 | Ssc5d | Tnxb |
| Rgs20 | Gm45304 | Sult1c2 |
| Hoxc9 | H2ac18 | Fam20c |
| Gm37233 | Gm9195 | ccdc198 |
| Hoxb7 | Alox5 | Rfx2 |
| Notch3 | Axin2 | Ppbp |
| Agtrap | Mab21l4 | Trib2 |
| Chic1 | Acox2 | Muc4 |
| Slc27a6 | Gstm2 | Ppp1r14c |
| Slc44a3 | Cst6 | Ehf |
| Tbx20 | Kcnh2 | Fbxo2 |
| Ttc12 | Cd177 | Ugt2b36 |
| Pced1b | Fam25c | Ckmt1 |
| Enpp5 | Lmntd2 | Muc5b |
| Itpkb | Arhgdib | Hs3st1 |
| Dlx3 | Hsd3b5 | Gm30214 |

| | | |
|---|---|---|
| Hoxb4 | Gm9113 | Cbln1 |
| 1700017M07 | Naalad2 | Serpina1a |
| Repin1 | Zfp978 | Adgrf5 |
| Hey1 | Tenm4 | Neurl1b |
| Jag2 | | Ppargc1b |
| Zfp518b | | Lcp1 |
| Mbp | | Pgk1-rs7 |
| Rtl8b | | Phf21b |
| Ifi47 | | Ugt2b35 |
| Oplah | | Fam131b |
| Hoxb6 | | Kctd15 |
| Tdrkh | | Thbs2 |
| Ass1 | | |
| Rab11fip4 | | |
| Rtl8a | | |
| Gca | | |
| Lhpp | | |
| Abca1 | | |
| Foxl2 | | |
| Slc39a8 | | |
| Casd1 | | |
| Lbx1 | | |
| Cdh10 | | |
| Tcea3 | | |
| H2-T23 | | |
| Tmem121 | | |
| Gulp1 | | |
| 4930471E19Rik | | |
| Prdm8 | | |
| Pla2r1 | | |
| Pou6f1 | | |
| Tap1 | | |
| Fst | | |
| Mta3 | | |
| Tmem150c | | |
| Kcnq1ot1 | | |
| Adcyap1 | | |
| Plxnb1 | | |
| Glt8d2 | | |
| Snrpn | | |
| B3galnt1 | | |

Nme4
Gm7575
Hoxd8
Il3ra
Grik4
Cyp27a1
Csf2ra
Hoxa5
Gem
Dab2
Papss2
Cers4
Rnf39
Nfatc2
Ctla2a
Apobec3
Gdf15
Has2
Lipo3
Cpeb1
Cmtm3
Cxxc4
Ltk
Rab3a
Plcl1
Mal2
Snca
Amigo2
Ndp
Crisp1
Gjb5
Pax6
Pde5a
Rasgrp3
Armcx2
Rtl5
Cmtm8
Acsf2
Cish
Hebp2
Ophn1

Slitrk5
Hsf4
Rbm12b1
Tns1
Map7
Chadl
Ociad2
Hoxc6
Nap1l2
Hecw1
Kctd12
Armcx3
Rab38
Nrk
Bcl7a
Rgs5
En2
Ramp1
Zfp536
Hoxb5
Gria3
Adam12
Tmem200a
Unc93b1
Hoxb3
Dnaja4
Alpk1
Fam78b
P3h3
Phactr2
Snhg18
Hoxa1
A930007A09Rik
Gap43
Hcn4
BC034090
Mapk12
Vangl2
Prickle3
Ccn4
Bnc1

Pdk4
Fuom
Ifih1
Mapk4
Trp53cor1
Cpt1c
Creb3l1
Adamts10
Hoxa7
Cadm2
Lgi2
Fstl5
Mageh1
Hoxd9
Mical1
Adamts7
Tram1l1
Tex15
Zfp677
Cntnap1
Col11a1
Ndn
Gm37035
Prokr2
Tmem159
Aplp1
Bhlhe22
Herc6
Htr2a
Ctsf
Kcp
Il15
Gstt3
Gabre
Cited4
Rbm43
Klrc1
Casp7
Tmem74
Gm20467
Gm37277

Ccdc149
Trim12a
Gm43162
Tmem47
Dpyd
Cntnap5a
Nrip3
Hotairm1
5730409E04Rik
Mark1
Car12
Tbkbp1
Tlr5
Hykk
Hoxc4
Plxdc2
Hoxc10
Hoxb3os
Lpar4
Itm2a
Hook1
Msx2
Gm6133
Cbx7
Hoxa9
Gm12608
Slc35g2
S1pr1
Dclk1
Ror2
Zfp629
