## Supplementary Table 2 for "Mutant IDH uncouples p53 from target gene regulation to disable tumor suppression"

| BIOCARTA_P53_PATHWAY | INGA_TP53_TARGETS | CEBALLOS_TARGETS_OF_TP53_AND_MYC_UP | KANNAN_TP53_TARGETS_DN | CEBALLOS_TARGETS_OF_TP53_AND_MYC_DN | TANG_SENESCENCE_TP53_TARGETS_UP | GHANDHI_DIRECT_IRRADIATION_DN |
|---|---|---|---|---|---|---|
| BioCarta Pathways | Inga et al. 2002 (Pubmed 12446780) | Ceballos et al. 2005 (Pubmed 15856024) | Kannan et al. 2001 (Pubmed 11402317) | Ceballos et al. 2005 (Pubmed 15856024) | Tang et al. 2007 (Pubmed 17533371) | Ghandhi et al. 2008 (Pubmed 19108712) |
| Mdm2 | BAX | STAT5B | NOP14 | TRIM31 | A2M | EVI5L |
| Bax | BBC3 | HSPA2 | PMP22 | CSK | ABCA8 | BRINP3 |
| Ccnd1 | CCNG1 | HSPH1 | CPM | DGKZ | ADAMTS5 | ELANE |
| Ccne1 | CDKN1A | DRG1 | TMEM97 | GPM6B | ALDH3A2 | SYBU |
| Cdkn1a | FASLG | NEK3 | COL18A1 | DDB2 | APLP1 | NFE2 |
| Gadd45a | FOS | MYC | BRCA1 | ATRX | BACE1 | PBX2 |
| E2f1 | GADD45A | SRSF7 | G0S2 | CHAF1B | C7 | HOXB3 |
| Rb1 | IER3 | INSIG1 | CCNE1 | NPRL3 | CCND2 | MIDN |
| Apaf1 | IGFBP3 | ISLR | PURA | PIM1 | CDKN1A | G6PD |
| Cdk4 | MDM2 | MTHFD2 | INA | SNCA | RAPGEF4 | OPRL1 |
| Mdm2 | PCNA | UAP1 | PPAT | PLAUR | COX7A1 | DEPP1 |
| Pcna | PMAIP1 | JUNB | SLC19A1 | REN | CCDC85B | RGMA |
| Timp3 | SESN1 | CTH | LAMC2 | F2R | GBX2 | NR3C2 |
| Trp53 | SFN | CD55 | BIRC3 | VARS1 | GMFG | S1PR1 |
| Cdk2 | THBS1 | PDIA3 | BIRC3 | KIF14 | GABBR2 | IRF1 |
| Bcl2 | TNFRSF10B | PFKFB1 | SORL1 | MLF2 | HBG1 | MDFI |
| | | ACSL1 | ARL4A | TCN2 | HBG2 | PLEKHA6 |
| | | ACSL3 | CDC6 | KRT8 | HSD17B2 | S100P |
| | | ZNF532 | APBA2 | VCAM1 | LCAT | MN1 |
| | | | EIF4A1 | CYFIP2 | MMP3 | FZD4 |
| | | | GTF2H2 | KIF21B | TSPAN12 | VSTM2L |
| | | | CXADR | COMT | NRG1 | FAM50B |
| | | | FOSL2 | SLC2A5 | IVNS1ABP | PER1 |
| | | | | POM121 | PSTPIP2 | ARHGEF25 |
| | | | | GLT8D1 | CLIP1 | TOX |
| | | | | CKAP2L | SIM2 | SEMA4D |
| | | | | HBEGF | SLC4A4 | OLFML2B |
| | | | | RNPEP | SNCA | CCDC85C |
| | | | | MDM2 | TFAP4 | PTK7 |
| | | | | ISCU | USF2 | DEPP1 |
| | | | | SEPTIN4 | VAMP8 | MRPS6 |
| | | | | SLC4A1 | WFDC1 | BCL11A |
| | | | | TPM1 | ZMAT3 | CNKSR3 |

KCNK3
PRDM6
ARHGEF37

| ONGUSAHA_TP53_TARGETS | KERLEY_RESPONSE_TO_CISPLATIN_UP | TANG_SENESCENCE_TP53_TARGETS_DN | BIOCARTA_P53HYPOXIA_PATHWAY | WARTERS_IR_RESPONSE_5GY | KANNAN_TP53_TARGETS_UP | KEGG_P53_SIGNALING_PATHWAY |
|---|---|---|---|---|---|---|
| Ongusaha et al. 2003 (Pubmed 12802282) | Kerley-Hamilton et al. 2005 (Pubmed 15940259) | Tang et al. 2007 (Pubmed 17533371) | BioCarta Pathways | Warters et al. 2009 (Pubmed 19580510) | Kannan et al. 2001 (Pubmed 11402317) | KEGG hsa04115 |
| TP53 | ACTA2 | ADARB1 | HIC1 | BEX3 | GDF15 | CDK2 |
| ZMAT3 | ALDH1A3 | ADH1B | HSPA1A | ACER2 | LSR | CDK4 |
| DGKA | ANK1 | NUSAP1 | TP53 | POLH | SMAD7 | CDK6 |
| DAXX | ANXA1 | DBF4 | ABCB1 | FAS | SCRIB | CDKN1A |
| SLC66A3 | ANXA4 | BARD1 | MDM2 | COQ9 | TMSB15A | CDKN2A |
| PHLDA3 | MS4A1 | BIRC5 | ATM | FBXW7 | TST | GADD45G |
| LPIN1 | BTG2 | BUB1 | CDKN1A | MDM2 | SUSD6 | CHEK1 |
| DDIT4 | TIGAR | TPX2 | TP53 | PHLDB3 | PMAIP1 | CHEK2 |
| PITPNC1 | CAV1 | CCNA2 | IGFBP3 | GDNF | HRAS | SESN3 |
| MDM2 | CDKN1A | CCNB2 | NQO1 | CHADL | MARCHF8 | DDB2 |
| ANXA8L1 | CXCL14 | CDK1 | ABCB1 | TYMSOS | DMPK | GADD45A |
| AK1 | DDB2 | CDC20 | IGFBP3 | AEN | ATF3 | RCHY1 |
| MATN4 | DDIT4 | CDC25A | AKT1 | B2M | GALE | BBC3 |
| CTSF | DGKA | CENPA | AKT1 | APP | FHL2 | SESN1 |
| CDKN1A | EPS8L2 | CENPE | HSP90AA1 | TIGAR | QSOX1 | SFN |
| ENPP2 | FADS3 | CTH | NQO1 | FAS | LCAT | APAF1 |
| COX6B2 | FDXR | EZH2 | NQO1 | DDB2 | PCNA | IGF1 |
| TOB1 | DRAM1 | FBXO5 | FHL2 | CDKN1A | APOC1 | IGFBP3 |
| CSRP2 | FOSL1 | FOSL2 | HIC1 | GADD45A | SLC3A2 | FAS |
| SAT1 | GADD45A | H2AC16 | TP53 | MDM2 | ZNF7 | CD82 |
| COL18A1 | GDF15 | IGFBP5 | TP53 | PCNA | GADD45A | MDM2 |
| IVL | GLS2 | IL12RB2 | TP53 | PPM1D | IGFBP6 | MDM4 |
| CCNG1 | HSD17B3 | KIF4A | TP53 | TNFRSF10C | NHLH2 | GADD45B |
| FAS | IER3 | KIF11 | TP53 | SH2D2A | SPAG1 | ATM |
| CTSH | INPP5D | KIF23 | TP53 | ATF3 | BCL6 | RRM2B |
| NQO1 | ISYNA1 | LMNB1 | TP53 | PLK3 | MAN2B1 | SERPINE1 |
| APOBEC1 | LMNA | MAF | GADD45A | FDXR | CDKN1A | SHISA5 |
| PLTP | PIDD1 | MAP1B | GADD45A | NINJ1 | ZP3 | GTSE1 |
| PTPRVP | NEFL | KIF20B | CDKN1A | RRAD | VCAN | SERPINB5 |
| ERCC5 | PDE4A | MRE11 | CDKN1A | CAVIN2 | BAK1 | PMAIP1 |
| EFS | PHLDA3 | NEK2 | HIF1A | GDF15 | DDB2 | CYCS |
| TAP1 | PLAT | NRGN | NFKBIB | HAS3 | CARTPT | ATR |
| TNFRSF18 | PLK2 | NUP62 | EP300 | POLH | LOXL1 | STEAP3 |

| | | | | | | |
|---|---|---|---|---|---|---|
| EPHX1 | PLK3 | OR1E1 | FHL2 | BTG2 | SCGB2A2 | PIDD1 |
| SERPINE2 | PPM1D | PDP1 | HIF1A | MDM2 | NDN | RPRM |
| FXYD3 | PROCR | EXOSC9 | GADD45A | OR11A1 | APLP1 | PTEN |
| ROBO1 | RPS27L | PPL | MDM2 | HSPA4L | PLK3 | ADGRB1 |
| SRXN1 | SERPINE1 | PRC1 | NFKBIB | BBC3 | FEZ1 | BAX |
| | SESN1 | PTTG1 | MAPK8 | SESN1 | EXT2 | CCND1 |
| | STK17A | PTTG3P | RPA1 | PLCL2 | BTG2 | RRM2 |
| | TNFRSF10C | QPCT | BAX | GASK1B | NINJ1 | BID |
| | FAS | KIF20A | TAF1 | NOTCH1 | ABCA3 | TP53AIP1 |
| | TP53TG1 | RACGAP1 | AKT1 | THSD1 | CTDSP2 | PERP |
| | TRIM22 | SMC4 | HSPA1A | TIGAR | BBC3 | COP1 |
| | ZMAT3 | SPAG5 | HSP90AA1 | DNAI2 | H2AC6 | ZMAT3 |
| | | AURKB | DNAJB1 | RHOXF2 | TAX1BP3 | SIAH1 |
| | | PLK4 | HIC1 | TP53INP1 | DCAF7 | THBS1 |
| | | TCF4 | CDKN1A | FBXW7 | NEFL | TP53 |
| | | TFAM | BAX | CDKN1A | ACTA2 | TP73 |
| | | TK1 | BAX | DNAI3 | DGKA | TSC2 |
| | | TOP2A | BAX | PIDD1 | FAS | CASP3 |
| | | PBK | TAF1 | ZNF597 | ENG | SESN2 |
| | | TREM1 | MAPK8 | VWCE | VIPR1 | CASP8 |
| | | TTK | MAPK8 | SNX20 | PDLIM4 | CASP9 |
| | | UBE2C | HIF1A | BLOC1S2 | PLIN2 | PPM1D |
| | | ZFY | FHL2 | LRATD2 | CES2 | CCNB3 |
| | | | FHL2 | E2F7 | PRKAB1 | TNFRSF10B |
| | | | | | SURF1 | CCNB1 |
| | | | | | SMTN | CCND2 |
| | | | | | | CCND3 |
| | | | | | | CCNE1 |
| | | | | | | CCNG1 |
| | | | | | | CCNG2 |
| | | | | | | CCNB2 |
| | | | | | | CCNE2 |
| | | | | | | EI24 |
| | | | | | | TP53I3 |
| | | | | | | CDK1 |

| WARTERS_RESPONSE_TO_IR_SKIN | WP_P53_TRANSCRIPTIONAL_GENE_NETWORK | GHANDHI_DIRECT_IRRADIATION_UP | PID_P53_DOWNSTREAM_PATHWAY | P53_DN.V2_UP | P53_DN.V2_DN | CONCANNON_APOPTOSIS_BY_EPOXOMICIN_DN |
|---|---|---|---|---|---|---|
| Warters et al. 2009 (Pubmed 19580510) | Ding et al. WikiPathways | Ghandhi et al. 2008 (Pubmed 19108712) | Schaefer et al. 2008 (Pubmed 18832364) | Elkon et al. 2005 (Pubmed 15892871) | Elkon et al. 2005 (Pubmed 15892871) | Concannon et al. 2007 (Pubmed 16983338) |
| ASGR1 | CDK2 | MYPN | TNFRSF10A | ADAM12 | ABCG5 | IL7 |
| NODAL | CDKN1A | MT2A | E2F3 | ALOX5AP | ACOX2 | DHFR |
| TRIAP1 | IRF9 | DUSP2 | APAF1 | AMELX | ACSM3 | TNFAIP6 |
| TIGAR | SIVA1 | CXCL5 | TNFRSF10B | ANGPTL3 | ADAM11 | POLD1 |
| NOPCHAP1 | SMR3B | PPP6R1 | TNFRSF10C | ANGPTL4 | AGPAT3 | PTX3 |
| MDM2 | CPT1C | TSLP | TP73 | ANXA10 | ALOX5 | TOP2A |
| THSD1 | ADORA2B | INHBA | SNAI2 | APOL3 | ANPEP | UBE2C |
| PRF1 | E2F7 | TRIAP1 | TRIAP1 | ATP12A | BMAL1 | MAD2L1 |
| RPS27L | DDB2 | TIGAR | BNIP3L | AVPR2 | ATP4B | PDGFRA |
| TGFBI | GADD45A | MMP10 | SH2D1A | BARX2 | ATP6V1B1 | PDGFRL |
| GDF15 | ERCC5 | SOD2 | PRDM1 | BAX | BCAT1 | STMN1 |
| TP53INP1 | FANCC | PTPRE | CCNK | BFSP2 | BCL2L11 | CDK1 |
| NINJ1 | SLC7A11 | MDM2 | DKK1 | XAF1 | BMP3 | MYC |
| RRM2B | MTOR | THSD1 | MAP4K4 | BPI | BMPR1B | CCNA2 |
| RAD51C | FUCA1 | BMP2 | EGFR | C4BPA | BMPR2 | SMAD6 |
| PAPPA | BBC3 | PLEK2 | TGFA | CA3 | C1orf21 | MYC |
| JAM3 | GLS2 | BCL2A1 | AFP | CACNG2 | C5AR1 | MAP2K6 |
| BLOC1S2 | SESN1 | SLC16A6 | TP53 | CCL20 | PPP1R17 | MYB |
| TNFRSF10C | SFN | SERPINB2 | SERPINE1 | CCL4 | CCNF | CD24 |
| CABYR | GPX1 | MMP3 | JUN | CD5L | CDADC1 | SLC18A1 |
| CFAP57 | APAF1 | MT1H | RB1 | CD68 | CENPA | HMGN2P9 |
| PALM2AKAP2 | ICAM1 | GDF15 | GPX1 | CD69 | CFD | TMSB10 |
| SNX20 | FAS | MMP1 | CTSD | CD83 | CHD1L | H4C12 |
| MTCL1 | FASLG | LAMC2 | SP1 | CDH4 | CLEC4M | TMSB4X |
| HSPA4L | IRF5 | RRM2B | MMP2 | CEACAM5 | CLTCL1 | PCOLCE |
| FDXR | LIF | AEN | MET | CH25H | CNTF | RLN2 |
| BBC3 | MIR145 | MT1G | BCL2 | CHRNA10 | CST4 | ABCA12 |
| DEPDC7 | MIR200C | MT1E | SPP1 | CHST8 | CYP27B1 | ESRRG |
| DNAI3 | MIR34A | CYP26B1 | VDR | CLCF1 | CYP39A1 | CADPS |
| ETV7 | MIR34B | NFKBIZ | PCNA | CLEC1A | DIO1 | H1-10 |
| ZNF219 | MIR34C | FGF2 | VCAN | CTLA4 | E2F2 | MFSD10 |
| PPFIBP1 | MGMT | FRMD4A | HGF | CXCL1 | ELAVL2 | AK4 |
| MDM2 | MLH1 | GADD45A | CCNB1 | CXCL12 | ELL2 | MARCKS |

| | | | | | | |
|---|---|---|---|---|---|---|
| DDB2 | MSH2 | DKK1 | LIF | CXCL2 | ENPEP | DOK4 |
| VWCE | NCF2 | TCIM | TYRP1 | CXCL5 | EPHA7 | PEBP1 |
| CCNG1 | NOTCH1 | TCIM | DDX5 | CXCL6 | EPHX2 | ICA1 |
| OR11A1 | RRM2B | TNFRSF10C | IGFBP3 | CYP17A1 | EVC | AGPAT1 |
| NOTCH1 | SERPINE1 | IFNE | ATF3 | CYP2A13 | EYA1 | AURKB |
| STING1 | PCNA | MT1X | EDN2 | CYP2C8 | FAAH | CDK1 |
| BTG2 | PRKAG2 | NAMPT | FDXR | CYP2D6 | FABP4 | RPS6KA2 |
| FAS | POLK | MDM2 | NFYA | DLX6 | FKBP5 | ACYP1 |
| FBXO22 | SERPINB5 | ASCC3 | GADD45A | DPF1 | FPR2 | ADCY1 |
| CAVIN2 | PRKAG3 | CXCL2 | APC | DPYS | FRS2 | SPTSSA |
| PLCL2 | PMAIP1 | IL33 | NFYB | EML3 | GAD2 | SPRY2 |
| COMP | PML | LRP8 | FAS | ETV5 | GJA10 | RTL8C |
| SESN1 | POLH | PRDM1 | CD82 | EVI2A | GMIP | IL7 |
| RTTN | DDIT4 | HSPA4L | DUSP1 | EVX1 | GNA13 | CHRNA3 |
| TMEM120B | DRAM1 | MT1B | S100A2 | FCRL2 | GPR75 | H4C9 |
| ZNF79 | PIDD1 | ZC3H12C | EPHA2 | FGF22 | GPSM2 | SIX3 |
| CES2 | PRKAA1 | MDM2 | CASP1 | FOSL2 | GRK4 | RGS4 |
| TRIM22 | PRKAA2 | VWCE | PML | FOXL2 | GUCY2C | RGS4 |
| FBXW7 | PRKAB1 | RELB | BDKRB2 | FPR3 | GZMM | VSNL1 |
| FGFBP3 | PRKAB2 | KYNU | SFN | FZD9 | HBD | NFE2L3 |
| MICALL2 | PRKAG1 | CLDN1 | SERPINB5 | GALK1 | HCG9 | APRT |
| TYMSOS | RPRM | C3orf52 | CDKN1A | LGSN | H1-4 | TSPOAP1 |
| ICAM5 | TIGAR | ANXA10 | COL18A1 | GP5 | H2AC13 | MIR124-1HG |
| TP53TG1 | PTEN | OR11A1 | MLH1 | GPA33 | HMMR | LINC00847 |
| SCN9A | ADGRB1 | POU5F1 | BCL6 | GSC2 | HNRNPA0 | MYOF |
| MAGT1 | RPTOR | CDKN1A | HTT | GUCY2D | HOXA11 | NFIB |
| ANKRA2 | BAX | HSD11B1 | MSH2 | GYPB | HOXA2 | ADM |
| CHADL | SAT1 | FAS | TSC2 | H6PD | HOXB3 | AURKA |
| DRAM1 | CCL2 | RGS9 | SMARCA4 | HAO1 | HOXD13 | AURKA |
| DNAJB4 | CX3CL1 | IL11 | CCNG1 | H1-6 | HP1BP3 | KIFC1 |
| SNORA24 | TP53AIP1 | IL6 | PMS2 | H2AC4 | HPGD | KIFC1 |
| ACBD6 | PERP | CXCL1 | CSE1L | HS3ST2 | HS3ST3B1 | PHOX2B |
| TIGAR | MLST8 | IL1A | CASP6 | HTR1E | HSF2BP | LMO1 |
| NDUFAF8 | ZMAT3 | G0S2 | FOXA1 | IER3 | HTN1 | LMO1 |
| CDKN1A | DEPTOR | EYA3 | BID | IKZF1 | IGFBP4 | MYCN |
| TMTC3 | SLC2A1 | TESC | PCBP4 | IL36A | IGFBP6 | CETN3 |
| RAP2B | AURKA | IL1B | PTEN | IL1RAPL1 | IL17A | CETN3 |
| GM2A | THBS1 | BTG3 | CX3CL1 | IL1RN | ITGA4 | INSIG1 |
| MDM2 | TNF | PLCL2 | BTG2 | CXCL8 | KCNIP1 | RGS13 |
| FRMD4A | TSC2 | LAMB3 | MDM2 | INPP5D | KCNJ8 | ELMO1 |
| PIDD1 | XPC | IL12A | E2F1 | INSL3 | KCNJ9 | CMC4 |

| | | | | | | |
|---|---|---|---|---|---|---|
| ANKRD33B | XRCC5 | SESN1 | POU4F1 | INSL4 | NDC80 | TMEM158 |
| PPM1D | BTG2 | BIRC3 | CAV1 | ITM2A | LILRA2 | LRRN3 |
| POU2F2 | NANOG | KYNU | TAP1 | KCNK3 | LIPE | ANAPC15 |
| E2F7 | ULBP2 | LIF | CEBPZ | KCNMB1 | LLGL1 | NUDT1 |
| FOXP1 | ULBP1 | MT1X | BAX | KRT14 | PRICKLE3 | EMILIN1 |
| LASP1NB | SESN2 | GPATCH2L | BCL2L1 | KRT31 | MMP16 | GM2A |
| BTG3-AS1 | ULK1 | TNFAIP3 | MCL1 | LALBA | MTHFR | CADM1 |
| UNC5B-AS1 | ACAD11 | TNFAIP3 | EP300 | PIDD1 | MYBPC3 | NELL1 |
| POLH | AKT1S1 | GCH1 | POU4F2 | LRRC32 | NFKBIB | MLLT3 |
| ALCAM | ALDH4A1 | CXCL3 | IRF5 | MAGEC2 | NOL3 | TENM4 |
| LINC00926 | TNFRSF10D | SLC7A11 | TP53BP2 | ME3 | NPL | ALDH7A1 |
| POLH | TNFRSF10B | TNFRSF10B | PMAIP1 | MMP13 | NRXN3 | MARCKSL1 |
| | CCNE1 | SERPINB2 | NFYC | MMP17 | NTF3 | PDK1 |
| | CCNG1 | PTGS2 | TFDP1 | MMP3 | NUDT6 | TMSB15A |
| | TP53INP1 | CXCL3 | E2F2 | MOG | PBX2 | TRAPPC6A |
| | TP53I3 | CXCL2 | HIC1 | MPZ | PCDHB13 | SCHIP1 |
| | TRAF4 | NEFM | BCL2A1 | MT1G | PDE6H | DHCR24 |
| | ISG15 | BLOC1S2 | TAF9 | NCKAP1L | PER3 | LSM7 |
| | ULK2 | PTGFR | BAK1 | NF1 | PGF | HMMR |
| | CDC25A | BBC3 | DUSP5 | NFIX | PIP | EID1 |
| | CDC25C | MT2A | TP53I3 | NFKBIA | PLEK2 | CDH11 |
| | SCO2 | DUSP2 | PPM1J | NTRK3 | PNMA3 | ETV1 |
| | | FRMD4A | STEAP3 | NTSR1 | PNOC | KIF23 |
| | | OSBPL3 | RPS27L | OR10H3 | POU3F2 | DNASE1L1 |
| | | NAMPT | RRM2B | OSGIN1 | PPIL2 | PLK1 |
| | | INHBA | RNF144B | OSGIN2 | NPY4R | DLGAP5 |
| | | TIGAR | TADA2B | P2RY11 | PRMT8 | MAD2L1 |
| | | SLC16A6P1 | CARM1 | P2RY6 | PTGDS | TFRC |
| | | MIR23AHG | JMY | PCDH1 | PTPN11 | SNRPG |
| | | PTGS2 | COP1 | PCDHA6 | QTRT1 | HMGN3 |
| | | C2CD4B | DGCR8 | PDC | RAB17 | HMGN3 |
| | | ANKRD20A1 | DDB2 | PF4V1 | RAB38 | C7 |
| | | CDKN1A | NDRG1 | PKD2L1 | RAB4B | HDAC9 |
| | | MYBL1 | HDAC2 | POU2AF1 | RAB8B | TBC1D30 |
| | | THBS1-IT1 | CREBBP | POU6F1 | RAD52 | ENSG00000291 |
| | | MDM2 | KAT2A | PPEF1 | RASAL1 | CHRDL1 |
| | | GPR68 | BCL2L2 | PRDM11 | RECK | DBI |
| | | CREB5 | CASP10 | PRKCE | RNF17 | AKAP1 |
| | | HIP1 | TP53INP1 | PRRG3 | RNF2 | MYC |
| | | SLC30A1 | PERP | PRTN3 | SCN9A | GNG11 |
| | | FAS | PPP1R13B | PTK2B | SELL | CADM1 |

| | | | | |
|---|---|---|---|---|
| FAM20A | RCHY1 | PTX3 | SERGEF | MTUS1 |
| PIDD1 | ZNF385A | PYY | SERPINF1 | LMO3 |
| SMC5 | ARID3A | RELB | SERPING1 | LUM |
| SLC7A11 | PRMT1 | RFPL2 | SIL1 | HMGB2 |
| E2F7 | GDF15 | RFX2 | SLC24A1 | ATP5MC1 |
| CREB5 | AIFM2 | RGR | SLC27A6 | IGSF3 |
| GDNF | BBC3 | RHD | SLC6A3 | S100A4 |
| SLC7A11 | BCL2L14 | RIPK4 | SLCO2A1 | VCAN |
| POLH | TP63 | RND2 | SMPX | VCAN |
| NPIPB13 | PLK3 | RNF39 | SNAI1 | PCLAF |
| NAMPTP1 | RGCC | RORC | SNPH | ESPL1 |
| CXCL8 | PIDD1 | CNOT9 | SNTB1 | PPFIA4 |
| | NLRC4 | SARDH | SREBF1 | SLC1A1 |
| | TIGAR | SDC4 | SSTR1 | SLC1A1 |
| | DROSHA | SLA | SSTR2 | SORL1 |
| | DDIT4 | SLC15A2 | SYT17 | CDC20 |
| | SCN3B | SLC16A4 | TAF12 | CYB5A |
| | TNFRSF10D | SLC38A3 | TDRD1 | CYB5A |
| | PYCARD | SLC4A10 | TINAGL1 | NDUFB1 |
| | PRKAB1 | SPINK1 | NKX2-1 | LIMK2 |
| | TRRAP | SPRR1B | TLR1 | METTL7A |
| | SESN1 | SPRR2C | ANO2 | TMX4 |
| | | SYN1 | TNNC2 | EXOSC9 |
| | | TDRKH | TP53TG1 | KIFC1 |
| | | TFAP4 | TP73 | SPC25 |
| | | TNFAIP3 | TRPV6 | PPEF1 |
| | | TNFRSF9 | TTR | NDUFB6 |
| | | TNNI3K | TULP2 | DPY19L1 |
| | | TNR | BEST1 | MFAP4 |
| | | TP53 | ZNHIT2 | MXI1 |
| | | TPTE | | NME4 |
| | | TUBB1 | | H2AZ2 |
| | | WNT3 | | GTF2H5 |
| | | | | PFKP |
| | | | | RTN1 |
| | | | | MAB21L1 |
| | | | | C10orf95-AS1 |
| | | | | TFAP2B |
| | | | | NREP |
| | | | | CBX5 |
| | | | | FABP5 |

DNPH1
DUSP7
LDOC1
H4C3
ALDOC
TOP2A
GDF11
DACH1
TWIST1
CLCN4
SLC6A2
PTTG1
COX17
TMX4
BIRC5
ASCL1
ASCL1
NDUFA2
NFIB
CCNA2
KCNMA1
CHGA
LAGE3
HK2
RPH3A
RAPGEF4
RAPGEF4
ABHD14A
BUB1
NFIB
NHP2
BCL11A
LSM2
LDHA
BCKDHB
PLPPR4
MAOA
MAOA
MAOA
MKI67
MKI67

NFIB
POLR2L
CENPA
FZD2
CXCR4
RPA3
MXI1
MXI1
TOP2A

| CONCANNON_APOPTOSIS_BY_EPOXOMICIN_UP | FISCHER_DIRECT_P53_TARGETS_META_ANALYSIS | MARTINEZ_TP53_TARGETS_DN | MARTINEZ_TP53_TARGETS_UP | GRAESSMANN_APOPTOSIS_BY_DOXORUBICIN_UP | PEREZ_TP53_TARGETS | GRAESSMANN_APOPTOSIS_BY_DOXORUBICIN_DN |
|---|---|---|---|---|---|---|
| Concannon et al. 2007 (Pubmed 16983338) | Fischer et al. 2016 (Pubmed 27280975) | Martinez-Cruz et al. 2008 (Pubmed 18245467) | Martinez-Cruz et al. 2008 (Pubmed 18245467) | GRAESSMANN et al. 2007 (Pubmed 17160024) | Perez et al. 2007 (Pubmed 17563751) | GRAESSMANN et al. 2007 (Pubmed 17160024) |
| DUSP1 | ABCA12 | HUWE1 | SNRNP27 | NUDT22 | EPHB3 | TLCD1 |
| IL15 | ABCB6 | PABPN1 | RRN3 | ATP6V0E2 | PRR22 | HIKESHI |
| THBS4 | ACER2 | MRPL52 | TXNDC12 | SDHAF1 | ZDHHC11 | ERG28 |
| CD44 | ACTA2 | UBLCP1 | GJA1 | AAR2 | BEST4 | SNRNP40 |
| SAT1 | COQ8A | UBLCP1 | SCD | RBM43 | TTC39C | C17orf49 |
| HSPA6 | ADIRF | DDR1 | SERP1 | RETSAT | MEGF11 | TMEM176A |
| PSMD11 | AEN | CSNK1D | AGTR2 | YKT6 | GAMT | CCDC127 |
| PSMD12 | AGBL5 | JUNB | LDHC | COPS9 | DUSP19 | NEURL4 |
| SSX2 | AK3 | IMPACT | TGFBI | JOSD2 | FGF1 | CZIB |
| CYP4F3 | AKAP9 | MARCKS | SGPL1 | TMEM140 | DOCK8-AS1 | CEP20 |
| PSMD1 | ALDH1A3 | ABCE1 | LPL | HHATL | KCNG3 | TRMT112 |
| CYP4F2 | AMZ2 | CCT2 | ACTC1 | EOLA1 | MMEL1 | CUTA |
| FAS | ANKRA2 | SMAD1 | SMOC2 | ENTREP3 | KBTBD6 | TMEM167A |
| HSPA4 | ANXA4 | MAP4 | SLC2A4 | UBALD2 | LACC1 | RMND1 |
| ELL2 | APAF1 | FKBP5 | HDLBP | CLDND1 | LACC1 | LMBRD1 |
| DNAJB4 | APOBEC3C | ERRFI1 | CASP6 | NAT9 | ZNF777 | LAMTOR4 |
| IL6R | APOBEC3H | RPS6 | TXNIP | ANGEL1 | UMODL1 | STEAP3 |
| SMOX | APOD | APBA3 | PRDX1 | BRIX1 | ARID2 | PPIH |
| EIF5 | ARAP1 | HMGA1 | CLDN11 | RGCC | ABCC12 | CEBPZOS |
| BAK1 | ARFGEF2 | PHB2 | TSPAN3 | DRAM1 | FBXL21P | NUDT16L1 |
| BBC3 | ARHGAP42 | RPL37A | CCN1 | FRMD8 | ABCA13 | TEFM |
| INPP5D | ASCC3 | CTNS | H1-2 | IKBIP | FOXC1 | C11orf58 |
| MXD1 | ASTN2 | PSMC4 | CCND2 | LRWD1 | KCNJ5-AS1 | SARNP |
| RRAD | ATF3 | PSMC4 | RAD21 | TSPAN33 | CEL | SPHK1 |
| DUSP4 | BAX | PRELP | CYP4B1 | YIPF2 | RASEF | TMEM109 |
| TGFBR2 | BBC3 | PDCD7 | IL33 | ERO1B | MYO10 | MCMBP |
| MBOAT7 | BBS2 | PFKFB3 | AQP1 | CHP1 | XG | CARD19 |
| MAP2 | BCL10 | IGDCC4 | GPD1 | RFLNB | GALNT18 | NDUFAF4 |
| DDIT3 | BCL2L1 | ATP2A2 | ADH1C | FAM216A | PHF21A | SNHG6 |
| LGALS8 | BCL6 | EFTUD2 | NADK | RNASEH2C | SNX32 | RITA1 |
| MDM2 | BCL9L | CLCN3 | MCM6 | METTL6 | TRIM73 | NSMCE2 |
| SKIL | BHLHE40 | SPINT1 | RGS4 | C19orf12 | KCNS2 | PRXL2C |
| MDM2 | BLCAP | SLC1A5 | RGS4 | LRRC51 | CERS3 | TOMM5 |

| | | | | | | |
|---|---|---|---|---|---|---|
| GDF15 | BLOC1S2 | UBE3A | IL2RG | OSGIN1 | CERS3 | PRPF38B |
| GADD45A | BRMS1L | PLOD2 | LITAF | NAT8B | NOCT | UFC1 |
| FOS | BTBD10 | S100A10 | PRELP | SGF29 | FKBP6P2 | SLC44A2 |
| FOS | BTG1 | PAM16 | PC | CEP112 | DRAXIN | LYSMD3 |
| MDM2 | BTG2 | CRYM | PEA15 | C22orf23 | GINS4 | PPP4R3A |
| CDKN1A | BORCS7 | CRYL1 | ACOX1 | LARP1B | KIAA1217 | C1QTNF12 |
| CD44 | NOPCHAP1 | UBL5 | NSMAF | SLX9 | HILPDA | RREB1 |
| STIP1 | TIGAR | NCDN | CAT | CHAC1 | MOXD1 | FOXK2 |
| SPP1 | LINC02875 | C1D | DDX3X | TCIM | MINDY3 | LGALSL |
| DUSP3 | NDUFAF8 | SLC25A10 | PDPK1 | AAMDC | FAM13C | TRMT1L |
| ATF3 | CCDC90B | SOX2 | LXN | IPPK | ARHGAP27 | C16orf74 |
| CDK11B | CCNG1 | FOXO1 | NR4A1 | RAB43 | CEP120 | SMIM11 |
| GTF2A1 | CD82 | CYP17A1 | FSCN1 | TTC39B | DUOXA1 | TANC1 |
| ANXA2 | CDC42SE1 | FABP4 | BAD | SMIM3 | KCNS2 | ERGIC1 |
| TNKS | CDH8 | TRIM2 | BAD | ALKBH4 | CEMIP | TRIL |
| BCL2L11 | CDIP1 | KRTCAP2 | GLRX | TECPR1 | TRIM7 | RAVER1 |
| HSPA1A | CDKN1A | MEIS2 | ME1 | SPNS1 | IPP | THA1P |
| ZNF10 | CEL | ETHE1 | EIF3A | PTRH1 | MTUS2 | RRP8 |
| TAF13 | CFLAR | WDR12 | PRKAR1B | CES2 | FCHO2 | RSRC2 |
| ME1 | CHST14 | PDK4 | DNAJB1 | CYSRT1 | MCF2 | HSPBP1 |
| IDE | CLCA2 | NDUFA5 | WEE1 | RNF225 | TFAP2E | DIMT1 |
| PHLDA2 | CLP1 | MTA1 | IDH3G | PSAPL1 | CAMTA1 | IFT46 |
| FILIP1L | CMBL | ATF1 | TIA1 | C19orf47 | FBXL21P | NFIC |
| RFPL1 | CNOT6 | ITPR3 | AMD1 | TMEM158 | FBLIM1 | YAE1 |
| CCNT2 | COBLL1 | FZD6 | LY6D | SFI1 | ZMAT3 | RBM26 |
| CREM | COL7A1 | DMD | SLC25A10 | TMBIM1 | ZAP70 | POLR1HASP |
| CREM | CPEB2 | SULT2B1 | SIRPA | PSME3IP1 | TREML1 | DDI2 |
| CAMK2G | CPSF4 | NDUFA2 | SIRPA | RIPOR1 | BAP1 | DNAJC24 |
| ZNF185 | CROT | TUBA4A | C1QB | ENDOD1 | IRGQ | ADISSP |
| LMO2 | CSF1 | ELOVL6 | SLC6A4 | ARHGAP27 | MICALL2 | ARMT1 |
| SLC7A5 | CSNK1G1 | ELOVL6 | RBM6 | CAVIN1 | MTSS2 | CCSAP |
| PSMD13 | CYFIP2 | CCND1 | FAH | TEPSIN | HUNK | EFCAB2 |
| EPB41 | CYSRT1 | GNMT | CAPZB | LRR1 | B3GNT7 | KANSL1 |
| TOP3B | DCP1B | CD248 | PTGS2 | HAUS2 | MAGEE1 | CFAP141 |
| SUSD5 | DDB2 | CAP1 | LRG1 | TM2D2 | ZNF451 | STAMBPL1 |
| ENSG00000275 | DDR1 | CAP1 | CAV2 | CCDC166 | HOMER2 | C11orf98 |
| CLTB | DGKA | CTSB | PEG3 | ZBTB8A | MROH2A | TMEM176B |
| JUN | DHRS3 | LYAR | COX6A1 | MTFR1L | MROH2A | CCNQ |
| SMG1 | DNAJB5 | NFE2L3 | SRGN | RNF181 | ANKRD65 | LRRC58 |
| SMG1 | DRAM1 | EIF4EBP1 | FNTA | WIPI2 | PRRT3 | FUOM |
| ACTA2 | DUSP14 | CTNNBIP1 | IFT46 | UBE2F | TLNRD1 | RETREG1 |

| | | | | | | |
|---|---|---|---|---|---|---|
| AKR1C1 | DYRK3 | SUPT6H | PLAC1 | NT5DC2 | ZNF654 | TMEM126A |
| CHM | E2F7 | MYBL2 | FABP1 | KIAA2013 | ZNF654 | COX16 |
| IFRD1 | EDA2R | NECTIN2 | CKM | FAM32A | DRAXIN | CXorf38 |
| CTH | EDN2 | MYH6 | SOD3 | ARL16 | MIR570 | SNHG16 |
| STX6 | EFCAB10 | CERS4 | CDKN1C | MTFR2 | MOSPD1 | MDP1 |
| TUBA4A | EFNB1 | SFXN3 | SLC4A10 | NOP9 | BLCAP | BOLA1 |
| SLC16A3 | EI24 | TNMD | APOA4 | TRMT5 | BLCAP | MMACHC |
| ZFR | EID2B | DCT | SNCG | MACROD2 | KCP | CYB5B |
| GCLM | EML2 | NEUROD4 | ESD | EMC4 | C7orf57 | SIL1 |
| CDK11B | TEPSIN | LTBP2 | CLDN5 | DOP1B | PHLDB3 | TMEM179B |
| RAB21 | EPHA2 | KERA | SPON2 | TTC27 | ZKSCAN1 | CCDC91 |
| HEG1 | EPM2AIP1 | ST14 | CFD | SKIC8 | MBNL1 | MFSD4B |
| BSCL2 | EPS8L2 | NR2F1 | SLC39A3 | UFSP1 | BCL2L11 | IARS2 |
| CAP2 | ETV7 | KRT25 | NDST2 | RTRAF | LUZP1 | ALKBH4 |
| CAP2 | F5 | VDR | CCL15 | DNAJC18 | MUC20-OT1 | MAGOHB |
| PLAT | GASK1B | USP18 | CCDC71 | MOB2 | AHNAK2 | BORCS7 |
| SSX2 | FAM210B | FXYD4 | RGCC | AEN | ARIH1 | SNHG8 |
| PHKG2 | INKA2 | PFN2 | CPD | UBE2T | SLC46A1 | COA7 |
| HMOX1 | LRATD2 | PFN2 | CFL2 | LRRC57 | SORBS2 | USE1 |
| CCPG1 | FAM98C | KRT23 | APOC2 | CALHM2 | PLK5 | SLC44A1 |
| NSFL1C | FAS | NME7 | NDUFS4 | OLA1 | ZNF765 | ASNSD1 |
| NSFL1C | FBN2 | COL18A1 | UCP1 | DPH7 | ZNF765 | ABHD17C |
| UBR4 | FBXO22 | RBFOX2 | LIMS1 | MTARC2 | CEP164 | TRAK1 |
| WWTR1 | FBXW7 | EFNB1 | LIMS1 | PHYKPL | CYP4F35P | DMAC2 |
| PRDM2 | FCHO2 | LPIN1 | GBP2 | URM1 | CGAS | TEN1 |
| MAGEB1 | FCHSD2 | SEPHS2 | GET3 | SLC25A42 | ELFN2 | COX20 |
| NTS | FDXR | SLC7A5 | CPT1B | CCDC137 | MVP-DT | CMBL |
| SSX1 | FLRT2 | SDHAF4 | RECQL | ZNF593 | LINC02754 | MTFMT |
| SLC7A11 | GABARAPL2 | RETNLB | TNNC1 | OSER1 | MDFIC | C11orf52 |
| PXDC1 | GADD45A | LAD1 | CAV3 | SMPD4 | SLC25A34 | TMEM106B |
| PRSS12 | GBE1 | RIPK4 | TNFAIP6 | AHCYL2 | RMDN2 | KANSL2 |
| PRSS12 | GCH1 | MTF2 | MFAP5 | RIN2 | WDFY2 | TMEM79 |
| SAT1 | GDF15 | PTPRE | CXCL14 | ACOT6 | KLHL6 | NEAT1 |
| SPP1 | GNAI1 | PLIN4 | MTX1 | FBXL20 | TMEM165 | AARSD1 |
| P4HA2 | GNAS | COL11A1 | TFPI2 | SPRYD4 | ELFN2 | PSMG4 |
| HSPH1 | GPC1 | NOTCH1 | ALDH1A1 | PM20D1 | B3GALNT2 | EDEM3 |
| DOK1 | GPR87 | EGLN3 | AVPR1A | SIMC1 | MEIOC | C6orf136 |
| N4BP2L2 | GPX1 | ACTN3 | IL36A | TMEM184B | KIAA1217 | NR2C2AP |
| MXD1 | GREB1 | KRT34 | UBE2H | CCDC96 | AATBC | MRRF |
| GLS | GRHL3 | PLXNB3 | RBMS1 | CNBD2 | ZMIZ1-AS1 | LMF1 |
| LRIG1 | GSS | ALDH3A1 | TNNT2 | SPSB1 | CAST | NSMCE4A |

| | | | | | | |
|---|---|---|---|---|---|---|
| GLRX3 | HES1 | TRH | STAR | DRC3 | MEIOC | TMEM121 |
| TNFRSF10B | HEXIM1 | FADS3 | NIT2 | SLC35F6 | GLS2 | DCTPP1 |
| KIF1B | HHAT | LMTK2 | OPA1 | TUBB4B | AFF3 | CEP57L1 |
| TNFSF9 | H2BC21 | RPN2 | MS4A6A | MPHOSPH8 | DIP2C | PTRHD1 |
| KCTD20 | HRAS | CEBPB | NUBP1 | SPEF1 | ZNF160 | TP53I13 |
| SRP19 | HSPA4L | OVOL1 | CLDN15 | SUSD6 | ITPRIPL2 | NDUFAF7 |
| GSR | ICOSLG | FGFBP1 | ETFRF1 | HERPUD2 | CEP43 | WDR75 |
| COL16A1 | IER3 | NPY | GZMB | IDNK | CYP27C1 | TUBB2B |
| LDAF1 | IER5 | RHBG | SERPINB2 | ELAPOR1 | ENSG0000026C | PCF11 |
| MIR22HG | IGDCC4 | ACVR2B | STOM | DPCD | CYSRT1 | FRYL |
| CYB5R1 | IGFBP7 | DNAJC3 | NLK | PSRC1 | FAM43B | EEF1AKMT1 |
| FLNC | IKBIP | FHL2 | SLC22A2 | FAM110A | KCNIP3 | CHAC2 |
| MAP2 | IRF2BPL | KRT12 | PDGFC | OGFOD2 | ZFYVE28 | NSMCE1 |
| IL11 | ISCU | PAX6 | AKR1C3 | RHBDD3 | ANKRD20A1 | RRP1B |
| PCYT1B | ISYNA1 | HOOK2 | CD163 | LRRC56 | EME2 | RBM25 |
| PLAAT3 | ITGA3 | RARG | SOX4 | SLC25A33 | EME2 | SMIM30 |
| GABARAPL1 | ITPKC | NKD2 | HIPK3 | ACAD11 | ANKRD20A1 | CHMP1B2P |
| JMJD6 | KCNN4 | FOXC1 | ZFP37 | GFOD2 | GRN | JMJD8 |
| MAP1A | KCTD1 | KRTAP15-1 | TCEA1 | TRABD | CKB | PTMS |
| SYCP2 | MARF1 | MSH4 | ACAD8 | RPE | MLF2 | WDSUB1 |
| MICB | KIAA1217 | IRX4 | PON3 | INAVA | BLCAP | PBRM1 |
| SSX1 | KLHDC7A | CLEC10A | ATP2A1 | CRELD2 | CPE | STRADA |
| HSPA6 | KLHL12 | TNNT1 | GOSR2 | NABP1 | LRPAP1 | QRICH1 |
| LIN37 | KRT15 | PURB | S100A8 | RTP4 | BTG2 | MND1 |
| SPATA2 | KRT8 | TLE3 | EREG | RAP2A | ADRM1 | LYRM2 |
| LPXN | LCE1E | GLIPR1L2 | IMMP2L | SP110 | TRAK1 | WDR77 |
| PLD3 | LIF | PADI3 | IL1R2 | NUB1 | ENO2 | DRAM2 |
| SYT2 | LIMA1 | PLEC | MS4A6A | NIP7 | BAP1 | SCAF11 |
| TAC1 | LMNA | KRT72 | BBOX1 | IFITM10 | BUB3 | PRELID1 |
| SLC7A1 | LNPEP | CSF1R | IER3 | PPCS | BUB3 | CCDC50 |
| C18orf25 | LPXN | ARHGEF25 | LTC4S | ZNF729 | JUNB | PPDPF |
| CALCA | LRP1 | POM121 | HCAR2 | SLC10A6 | ID2 | UBE2G1 |
| PRKCZ | LRPAP1 | DNAJC17 | TIMMDC1 | SMIM7 | DLG5 | SMIM10L1 |
| SERPINB8 | MAP3K12 | SPESP1 | CLEC4E | C8orf76 | EGR1 | TMEM209 |
| DTNA | MARCHF8 | SNTB2 | RPTN | ATG101 | EGR1 | SNHG5 |
| SYNJ2 | MAST4 | PRSS12 | LILRB3 | RABEPK | ECE1 | DTL |
| TUBA4A | MBD5 | KRT35 | SREK1IP1 | ESRP2 | DVL3 | NSUN4 |
| MAFF | MCC | KRTAP5-2 | PRKD3 | SLAIN1 | AFF1 | NCAPD2 |
| SLC38A6 | MDM2 | MST1R | HAS3 | FLVCR1 | EXT2 | OLA1 |
| CLU | MFAP3L | PSORS1C2 | PRKACB | GBP7 | LTF | DPY30 |
| HSPB1 | MFGE8 | UBC | APP | GGCT | EFNA1 | FASTKD2 |

| | | | | | | |
|---|---|---|---|---|---|---|
| GLA | MGRN1 | DKK2 | UCP3 | EOGT | MX1 | RRP15 |
| NQO2 | MICALL1 | GPRC5D | DTX3 | UNC119B | TRAK2 | SKA1 |
| ITGA7 | MLF2 | SLURP1 | CHRND | ZFAND2A | CLK3 | TMEM135 |
| ORC3 | MON2 | KRTAP4-9 | SPRR2D | FAM83H | NEDD9 | ARHGEF10L |
| TMF1 | MRPL39 | TPO | QSOX1 | RIC8A | CELF2 | TMEM216 |
| ZYX | MRPL49 | HSPA8 | PTPRF | INO80E | CELF2 | FAM220A |
| UBE2H | MUC2 | VAMP8 | FGFR2 | GGNBP2 | CELF2 | RILPL1 |
| BLVRB | NADSYN1 | CRYBA4 | LANCL2 | CLP1 | BLMH | PHYKPL |
| KLF6 | NCEH1 | TENM2 | DCTN4 | C6orf132 | SUSD6 | PABIR1 |
| PPP1R15A | NDUFV3 | ELOVL3 | ADD1 | SLC35D1 | LIMK2 | UQCC2 |
| PSMA5 | NEFL | KRT32 | ASPH | C11orf68 | GAK | FAR1 |
| LAMP3 | NHLH2 | PMFBP1 | MAN1A2 | INCA1 | CDKN1A | PAGR1 |
| KLHL21 | NINJ1 | KRTAP6-3 | CNOT4 | GPAT4 | TRIM2 | NORAD |
| GEM | NKAIN4 | PTPRF | ANGPTL2 | DEPDC7 | MAML1 | TMEM175 |
| DYNC1H1 | NTPCR | ELN | PITPNB | LPGAT1 | MXI1 | DMAC1 |
| TTC1 | NUPR1 | CTNNAL1 | ACYP1 | PKDCC | IGF2 | RPL22L1 |
| UPP1 | NYNRIN | SRRM1 | GNA12 | ABCB10 | HDAC5 | ZNF706 |
| DNAJB2 | OBSL1 | SON | GSTA2 | TAP1 | DHRS3 | C2orf76 |
| LMNA | ORAI3 | ENC1 | MRC2 | ABHD4 | MLXIP | C15orf40 |
| AKR1C3 | PADI4 | CA6 | RAMP2 | ABHD5 | IRF1 | FAM118A |
| SLCO3A1 | PANK1 | AGPAT1 | SNRPN | ABTB1 | AK1 | EARS2 |
| DNAJC2 | PARD6B | ADGRG1 | SERPINA12 | ACAD8 | AK1 | FOXN2 |
| FAS | PARD6G | EDN3 | RTN4 | ACAD9 | TMEM63A | LIMCH1 |
| FAS | PBLD | FOXP1 | RPGRIP1 | ACADSB | TRIM26 | RNF145 |
| IFRD1 | PCNA | NFIA | ITGAV | ACADVL | H2BC21 | C3orf14 |
| MEST | PDE4C | ATM | LGALS9 | ACOX2 | ITM2A | TMEM186 |
| PHTF1 | PHLDA3 | EDARADD | MEF2A | ACOX3 | ITM2A | E2F8 |
| TSPAN9 | PHPT1 | PIGB | SGCG | ACSL1 | LPCAT3 | UBR5 |
| SLC3A2 | PIDD1 | SH3RF1 | PECAM1 | ACTR1B | GAA | ZNF329 |
| NQO1 | PLK2 | MAP3K5 | HSPB7 | ADA | ADM | ATRX |
| WARS1 | PLK3 | HHIP | IKZF1 | ADAM15 | ADAM12 | GID8 |
| SERPINE1 | PLXNB2 | KRT31 | CA13 | ADAM1A | ARNT2 | C5orf34 |
| NEAT1 | PMAIP1 | KRTAP19-8 | PLA2G2F | ADAM8 | ZBTB5 | SEPTIN10 |
| F2RL1 | PML | VN1R17P | MAP2 | ADAMTS15 | CHST15 | RHBDD1 |
| CASP7 | POLH | S100A3 | IL36RN | ADAMTS4 | KLHL21 | KLHL24 |
| ALDH1A2 | POLR2H | TCF20 | PDLIM5 | ADAMTS7 | FECH | CKAP5 |
| RIT1 | POU2F2 | IGFBP4 | PTPN14 | PLIN2 | FECH | POMK |
| CEBPB | POU3F1 | DLX3 | RACGAP1 | ADRB2 | IFIT1 | FRMD6 |
| ACADVL | PPFIBP1 | AFF2 | KLK1 | AGPAT5 | GATM | PPP2R3C |
| ICAM2 | PPM1D | KRTAP9-9 | KLF3 | AGTPBP1 | B4GAT1 | TCTN3 |
| IL15 | PPP1R3C | POU1F1 | CDKN1A | AHNAK | ABCB6 | TAF1D |

| | | | | | | |
|---|---|---|---|---|---|---|
| MDM2 | PRDM1 | FGF7 | PSAP | AHR | FAM53B | BCDIN3D |
| ZRSR2 | PRKAB1 | PKDREJ | TWSG1 | AK1 | MYO6 | TOE1 |
| GDF7 | PRKAB2 | LGALS7 | MAP1B | AK4 | MYO6 | VWA1 |
| CEP170B | PRKD2 | EPHX1 | FGL2 | AKR1B1 | RFK | WDR87 |
| GOSR2 | PRRG2 | SMAD4 | NR3C1 | ALAD | LGALS9 | AP5M1 |
| GOSR2 | PSTPIP2 | PMEL | ST3GAL2 | ALAS1 | UNC119 | CFAP97 |
| ADRM1 | PTEN | GNA13 | TPP2 | ALDH1A1 | DFFA | ERI2 |
| STK17A | PTP4A1 | MT1A | MR1 | ALDH2 | PHF21A | OXSM |
| STK17A | RAB1A | RRAD | TCF12 | ALDH4A1 | EDEM1 | SCARA5 |
| TNPO2 | RABGGTA | RHOG | CAPN6 | ALOX12 | SH2B3 | SLC9B1 |
| TOM1 | RAD51C | LEP | NEDD4 | ALOXE3 | INPP5D | SAAL1 |
| GADD45G | RALGAPB | ADIPOQ | TSHR | FAM117B | LDB1 | SLAIN2 |
| SERPINH1 | RAP2B | PMEPA1 | ABCB8 | AMPD2 | COL15A1 | CNEP1R1 |
| SERPINH1 | RCBTB1 | CHEK2 | CCL20 | ANGPT4 | ABCA1 | RASSF8 |
| CEBPG | RETSAT | FKBPL | SERPINB9 | ANGPTL2 | CCS | PCMTD2 |
| AQP3 | REV3L | FBN2 | TPSAB1 | ANKRD10 | LOXL1 | GPATCH8 |
| PCYT1A | RFK | DIAPH3 | NPY4R | ANKRD17 | IGF1R | ALG14 |
| MAP1LC3B | RGL1 | INCENP | ART1 | AOX1 | IGF1R | TMEM80 |
| CAMK2G | RGMA | PPP4R3B | RNF14 | AP4S1 | DGAT1 | DPH6 |
| DDIT3 | RGS16 | KRT17 | NSF | APAF1 | CX3CL1 | TSEN15 |
| KIAA0930 | RIN1 | TFDP1 | TBK1 | ATG3 | FRZB | ETAA1 |
| SLC12A4 | RNF144B | ECRG4 | BNIP3 | APOBEC1 | FRZB | TMEM144 |
| STX16 | RNF19B | GJB2 | PEX13 | APOF | ALDH4A1 | ATL3 |
| RRAD | RPS19 | POLG | ACSL1 | APOH | ITPKB | NCAPG2 |
| KIAA0319 | RPS27L | TFAP2B | ILF3 | AQP1 | AQP3 | INTS7 |
| ABLIM3 | RRAD | RIOK2 | MYL4 | AREG | ARHGEF17 | SLF2 |
| PDLIM3 | RRM2B | BICC1 | TMEM45A | ARMC7 | MANBA | DENND1A |
| RAB36 | S100A2 | SOX5 | KRT6C | BMAL1 | VSNL1 | UBE2W |
| UBFD1 | SAC3D1 | MAX | CASQ1 | ARRDC1 | VSNL1 | TSPAN2 |
| ARHGEF2 | SARS1 | NEO1 | ARPP19 | ARVCF | HBEGF | TMEM268 |
| ARHGEF2 | SCN2A | CCNL1 | AHR | ASAH2 | WDR47 | THOC2 |
| TSPYL2 | SCRIB | GSDMA | CTR9 | ASS1 | MMP11 | SFR1 |
| ACHE | CAVIN2 | COL1A1 | CTR9 | ATF3 | IQSEC1 | COA5 |
| DYNC1I1 | SEMA3B | MAN2C1 | BICD2 | ATOX1 | IQSEC1 | ZNF790 |
| DYNC1I1 | SERPINB5 | APLP2 | LIPE | ATP13A2 | HPGD | HJURP |
| HSPA4L | SERPINE1 | BAMBI | XRN2 | ATP6V0B | HPGD | AMZ1 |
| DAP3 | SERTAD1 | IGFBP4 | HMGA2 | ATP6V0C | CHST2 | SNRNP48 |
| ZFP36 | SESN1 | IGFBP4 | GZMH | ATP6V0D1 | HLA-DMB | MAGI3 |
| FAT1 | SESN2 | SHISA2 | SERPINB4 | ATP6V1B2 | ACVR1 | PIMREG |
| C1S | SFN | SHISA2 | SERPINB4 | ATP6V1C1 | KIF3B | PHF20 |
| ELL2 | SFXN5 | RASL11B | RNF114 | ATP6V1D | ARG2 | TMEM168 |

| | | | | | | |
|---|---|---|---|---|---|---|
| PCSK1 | SLC12A4 | PTGDS | IP6K1 | ATP6V1E1 | KDM6A | PDS5A |
| MSX2 | SLC25A45 | WRAP73 | NT5E | UBE2O | CELSR2 | VPS35L |
| MSX2 | SLC30A1 | PKIG | NTN1 | ATOSB | CEBPA | ZMYM6 |
| AKAP6 | SLC46A1 | NUMA1 | FOS | N4BP2L1 | AGAP1 | ARGLU1 |
| DHRS3 | SLC4A11 | GRHL1 | PUM1 | SLC25A44 | FRY | C1orf159 |
| NUCB1 | SLC9A1 | HRAS | IDE | DCAF1 | MYRF | KLF13 |
| SQSTM1 | SMAD3 | RETREG3 | RILPL2 | ZNF708 | TPST2 | WDR82 |
| ATP2B4 | SMAD5 | RMND5B | TCAP | TMEM51 | EEF2 | APOOL |
| CDR1 | PPP4R3A | GCLC | VAPB | C9orf78 | GNG11 | OCEL1 |
| PALLD | SNX2 | PPCDC | CD36 | HELZ2 | HSD11B2 | CNTROB |
| DUSP3 | SOCS4 | HOMER2 | RPA1 | DDX60 | ENPP4 | PCMTD1 |
| DNAJB6 | SPATA18 | SLC27A4 | SORT1 | TMEM150A | KLHL20 | WASHC4 |
| ABCG1 | SRA1 | LPCAT1 | LAPTM4A | NMRK1 | KLHL20 | PTRH2 |
| MAP1B | STAT3 | CPSF4 | GIT2 | CCDC117 | TMEM243 | CCDC88A |
| RHBDD3 | STX6 | CAMKK2 | FBLN2 | MAP7D1 | GCH1 | MRM1 |
| HSPA9 | SULF2 | CCDC137 | ZIC3 | WRAP53 | CHKA | PCID2 |
| YKT6 | SUPT7L | ZCCHC9 | PCK1 | TKFC | ZNF195 | KRBA1 |
| YKT6 | SUSD6 | BMERB1 | NNAT | FAM43A | FLI1 | UTP25 |
| PLK2 | SYNC | CDKN1A | CCT6A | CLBA1 | GNA11 | DCAF8 |
| SEL1L3 | SYTL1 | GABRP | LMNB1 | PPP1R37 | CEP164 | DBF4 |
| TTC39A | TANC1 | USP19 | NPM3 | CRACR2B | VDR | KRCC1 |
| C11orf80 | TCAIM | TAOK1 | LYZ | VPS37B | VDR | BRAT1 |
| NACC2 | TEAD3 | SLC39A6 | LUM | PLEKHG6 | CHKA | DIP2B |
| ABHD3 | TET2 | CALHM5 | LRRC8C | KLHL25 | EDNRB | MFSD4B |
| DGKE | TGS1 | MIS12 | ACO1 | NLRX1 | EDNRB | D2HGDH |
| SNRK | TLCD1 | MFAP3 | CTDSP2 | CEP104 | SORBS2 | TTC28 |
| EPCAM | TLR3 | KIF1C | CTDSP2 | BAG1 | TECPR2 | MYORG |
| KITLG | TM7SF3 | CALD1 | PRR13 | BAIAP2 | TECPR2 | UCK2 |
| CREB5 | TMEM131 | NDEL1 | GPSM1 | GPANK1 | TRAF5 | ARF6 |
| CAP2 | TMEM243 | SLC39A11 | SAR1A | TM2D1 | REEP1 | CDKN1B |
| DNAJB1 | TMEM63B | EGFR | ARCN1 | BCAP31 | REEP1 | CEP152 |
| ANXA2 | TMEM64 | SOX9 | RNF41 | BCAR1 | CLP1 | MRTFA |
| ANXA2 | TMEM68 | PIP5K1C | NCAPD2 | BCL2L11 | FGFR3 | SND1 |
| GPX3 | TNFAIP8 | RHOV | HMGCS2 | BCL2L2 | FGFR3 | KIAA0319L |
| POLR3C | TNFRSF10A | WRN | CELF2 | BECN1 | CAMK1 | PPP4R3B |
| PGF | TNFRSF10B | PDS5A | CTTN | BET1L | GRK5 | TAFA5 |
| UFD1 | TNFRSF10C | RBBP6 | ROPN1L | WDR46 | FZD1 | RPAP2 |
| DOK1 | TNFRSF10D | C6orf132 | GINM1 | BLCAP | FZD1 | RIMOC1 |
| DOK1 | TP53 | SHMT1 | RXYLT1 | BMP1 | EDNRA | AAK1 |
| ME1 | TP53I3 | KRTAP6-3 | NCOA5 | CELF4 | BRPF1 | AATF |
| BAK1 | TP53INP1 | CEBPG | PTBP3 | BST2 | KCNQ1 | ABCB7 |

| | | | | | | |
|---|---|---|---|---|---|---|
| PTPRN | TRAF4 | AGFG2 | SDCBP2 | NACC1 | BID | ABCC1 |
| MAP3K14 | TRIAP1 | KRTAP19-3 | GLO1 | BTG1 | EVPL | ABCD3 |
| | TRIB1 | KRTAP19-8 | GLO1 | BTG2 | DMTN | ABCE1 |
| | TRIM22 | QKI | ALAS1 | BTG3 | BRD1 | ACAA2 |
| | TRIM32 | CUX1 | CAPZB | CAPRIN2 | TOX | ACACB |
| | TRIM35 | KRTAP19-4 | BPHL | C1QTNF1 | GABRE | ACBD6 |
| | TRIP6 | PLCB1 | AIF1L | C1QTNF5 | INPP4A | ACHE |
| | TRUB1 | KRTAP6-3 | ARHGAP1 | C1S | DZIP1 | ACIN1 |
| | TSKU | SLC25A30 | RAB40C | ZC3H12C | IRF4 | ACSL4 |
| | TSPAN11 | MASP1 | C4orf3 | EEIG1 | ABCG1 | ACTB |
| | TUBB6 | TRO | PSMF1 | ENTR1 | MMP12 | ACTR8 |
| | TXNIP | CLK1 | UCK1 | C4B | NUAK1 | ACVR1 |
| | TYMSOS | KRTAP19-1 | BCL2L13 | TBC1D24 | VCAN | ACVR1B |
| | UQCC1 | SCMH1 | MOB1A | NT5DC3 | VCAN | ADAM10 |
| | USP15 | POLR2A | DRAM1 | FAM8A1 | ACP5 | ADAM17 |
| | UTP3 | CWH43 | SLC35A3 | CABYR | MUC2 | ADAMTS5 |
| | VCAN | ZDHHC5 | CUX1 | CAMK2B | RAPGEF5 | ADAT1 |
| | VWCE | SIVA1 | THRSP | CAMP | ATP2B2 | ADCK1 |
| | DNAI3 | BAALC | GALNT11 | CAPG | INPP5E | ADD3 |
| | WDR81 | CELF1 | PITX2 | CAPN1 | DNAJC6 | ADK |
| | XPC | STRBP | TRIB1 | CAPN10 | COL9A3 | ADNP |
| | YAP1 | DDX3Y | MARCHF2 | CARHSP1 | CNKSR1 | AFG3L1P |
| | ZC3HAV1 | ZMYND11 | TMEM234 | CASQ2 | IFIT3 | AK3 |
| | ZMAT3 | TP53 | PRMT6 | CBX1 | SMAD7 | AKAP12 |
| | ZNF219 | IFFO2 | TM6SF2 | CCNG1 | EML1 | PALM2AKAP2 |
| | ZNF337 | UTY | ZNF106 | CCNL2 | EML1 | AKAP8 |
| | ZNF37A | CDH3 | WARS1 | CCPG1 | MYB | AKR1E2 |
| | ZNF385A | FLNA | IL18R1 | NOCT | CACNA2D2 | ALDH18A1 |
| | ZNF423 | EPRS1 | TNFRSF19 | CCS | CCNF | ALDH1B1 |
| | ZNF561 | MTREX | EMP2 | CD14 | GLDC | ALDOA |
| | ZNF564 | MGLL | HIF3A | CD207 | MAD1L1 | ALG12 |
| | ZNF79 | SMARCA4 | TGFBR2 | CD24 | APAF1 | ALG3 |
| | | MKI67 | GRB10 | CD274 | IL10RA | ANAPC1 |
| | | INTS6L | GRB10 | CD68 | VAMP5 | ANAPC11 |
| | | TMEM131L | COL4A5 | CDC25A | HPN | ANAPC4 |
| | | EDC4 | CD74 | CDC42EP3 | SHROOM2 | ANGPT1 |
| | | RCC2 | HLA-B | CDIPT | GPC4 | ANGPTL6 |
| | | CELF4 | TF | CDK5R1 | GPC4 | ANKH |
| | | NHSL1 | LMBR1 | CDKN1A | AHDC1 | ANKRD17 |
| | | COL6A2 | TGFBR3 | CDR2 | DOCK4 | KANK2 |
| | | TPR | CEACAM1 | CELSR2 | HAGH | ANLN |

| | | | | |
|---|---|---|---|---|
| CRIM1 | PLD1 | CENPE | VIPR1 | ANP32B |
| FA2H | RXRA | ARAP2 | ZBTB48 | ANXA6 |
| OS9 | SORBS1 | ARAP1 | IDUA | AOC3 |
| LAMA5 | SLC26A7 | CES2 | CRIP1 | AP1G1 |
| SCARA3 | IL36G | CETN2 | EFNA4 | AP2B1 |
| SLC39A14 | CHPT1 | CHMP1B | ARHGEF4 | AP3D1 |
| ENOX2 | CLDN10 | CIRBP | FGF13 | AP3M2 |
| SMO | OSR2 | CITED2 | FGF1 | APBB1 |
| RBBP8 | CAMK4 | CITED4 | USH1C | APBB2 |
| MYH3 | IGHG1 | CKAP2 | ABCD1 | SPEG |
| KRT83 | TPSD1 | CLDN23 | SMAD5 | APEX1 |
| SREK1 | TWSG1 | CLDN9 | SMAD5 | ATG10 |
| KDM5B | KRAS | CLEC4G | MAP3K14 | ATG5 |
| KRT33A | IFI27L2 | TPP1 | ALDH1L1 | AQP5 |
| FRYL | LAMA2 | CLSTN3 | EHHADH | NAA10 |
| KRTAP8-1 | BABAM2 | CLTB | IL1RAP | ARF6 |
| WWC1 | HLA-B | COL11A2 | GAD1 | ARFGEF1 |
| XIST | COA5 | COL17A1 | BMP2 | ARHGAP21 |
| CLIP4 | ZC3H11A | COL18A1 | BMP2 | ARHGAP5 |
| KRT81 | PPP3CA | COL4A4 | IL2RB | ARHGAP6 |
| RYR1 | DDX3Y | COMP | BAIAP2 | ARHGEF10 |
| KRT79 | SLC11A2 | CORO1A | BAIAP2 | ARHGEF12 |
| KRT86 | PJA1 | CORO2A | SMPDL3B | ARID4B |
| KRTAP3-1 | RIMOC1 | COTL1 | APC2 | ARIH1 |
| KRT75 | ATPAF2 | CPEB4 | MN1 | ARL6IP5 |
| HOXA3 | ADD3 | CPOX | GAMT | ARMCX1 |
| PRMT5 | QPCT | CPT2 | MDM2 | ARPC1B |
| KRTAP19-8 | SLC45A3 | CRABP2 | ANK1 | ARRB2 |
| ATF7IP2 | KPNA6 | CRAT | SMAD3 | ASB7 |
| EGR2 | SEPTIN8 | CRB3 | SMAD3 | ASH1L |
| GALR1 | ARRDC4 | CREM | SMAD3 | ASPH |
| KRT82 | PSME4 | CRIP2 | DCLK1 | ATAD1 |
| TP53 | CCN3 | CRKL | ARHGAP44 | ZFHX3 |
| HLA-A | SET | CRLF1 | GPR183 | ATE1 |
| KRT36 | FBXW8 | CROT | GSTT2 | ATF2 |
| TRBV28 | RDH10 | CSF1 | SAC3D1 | ATF4 |
| H2BC17 | OS9 | CSF3 | CASP10 | ATF6 |
| KRTAP19-3 | PCSK6 | CSTB | ISG15 | ATM |
| HERC6 | LANCL1 | CTSB | DPYSL4 | ATP10A |
| KRTAP20-1 | FTH1 | CTSE | DPYSL4 | ATP11A |
| MYH1 | EIF4G1 | CTSV | GHR | ATP1A2 |

| | | | | |
|---|---|---|---|---|
| RPL34 | C5orf15 | CUL4A | PDE10A | ATP2A2 |
| VIRMA | SREK1 | CX3CL1 | CYP17A1 | ATP5F1C |
| RAB12 | HNRNPLL | CXCL10 | GLS2 | ATP5MC1 |
| LSM12 | ZNF445 | CXCL3 | MSX2 | ATP5MC2 |
| KRTAP13-1 | MIR214 | CXCR4 | CHST1 | ATP6V0A1 |
| ACSL5 | PILRA | CXXC5 | CEP43 | ATP6V0A2 |
| DYNLT1 | CARS1 | CYCS | ANGPT1 | ATP9A |
| ATXN7L3B | SUCO | ZFTRAF1 | ANGPT1 | ATRX |
| ARL8B | SIX2 | CYP2F1 | MST1 | ATXN7L3 |
| POLR2L | BMP1 | CDR2L | KIN | CEP131 |
| DOP1B | EML5 | FAM50A | BCL2A1 | HEATR1 |
| TARDBP | SPAG5 | NOPCHAP1 | F8 | CNTLN |
| RPS24 | CLNS1A | ZMIZ2 | AKAP7 | CACHD1 |
| BASP1 | GSTZ1 | WIPI1 | BIK | DCAF1 |
| MYH14 | ATF2 | DHX58 | ITGAM | ANKRD13B |
| CANX | ABCC5 | NOP16 | TUFT1 | CPTP |
| MT2A | ACTA1 | LMBR1L | BRINP1 | RILPL2 |
| RPL21 | GNAS | ARHGAP39 | HSD17B1 | ODR4 |
| RNF149 | ZNF260 | DESI1 | GREB1 | UQCC4 |
| TECR | NDUFAF7 | TMEM191B | LIFR | ERCC6L |
| PATZ1 | C1orf54 | C21orf91 | CCNA1 | LPCAT1 |
| PINLYP | TMCC2 | TANGO2 | MSX1 | PRMT7 |
| KRTAP9-9 | SLC14A1 | VWA7 | FOXF1 | CDCA7L |
| SNCA | ATXN7L3B | CCDC86 | CLTCL1 | SLC44A3 |
| PANK1 | LMAN1 | PPP6R3 | KCNK3 | EVA1A |
| EFNA3 | PNPLA2 | CDHR4 | ITGA9 | RUSF1 |
| MGA | ACAA2 | MAVS | MERTK | OXNAD1 |
| TMOD2 | DHX40 | FICD | ASPA | TTI2 |
| CXXC5 | CENPV | GPN3 | FUT1 | MIOS |
| PPIH | NSUN4 | C12orf4 | MCC | TMEM161A |
| YPEL1 | ITIH5 | HGSNAT | CYP4F11 | KRI1 |
| AATF | STARD4 | OTUD5 | MAB21L1 | DHRS11 |
| APOE | ZMYM4 | DAB2 | ARHGAP6 | PLEKHG5 |
| SLC35A2 | TRIP13 | DAPK1 | ADRB2 | METTL22 |
| UBC | CALCOCO2 | DAXX | CILP | KLHL36 |
| HMGCS1 | IMMT | DBR1 | CROCC | ZNF768 |
| ADGRG1 | CD59 | DBT | LY6D | KIAA1217 |
| AQR | GNA13 | DCBLD1 | MAF | ZNF395 |
| RPS21 | RPS25 | DCTN3 | FOXF2 | CCDC102A |
| SLC44A1 | SCOC | DCXR | BDNF | TMEM60 |
| NAA11 | MRPL3 | DDIT3 | ANK3 | B9D2 |

| | | | | |
|---|---|---|---|---|
| CALR | RNF14 | DDIT4L | LGALS7 | MYL6B |
| CDH5 | HSPA9 | DDR1 | USP6 | CRYBG3 |
| PDAP1 | NUDT7 | DDX39A | LRRTM2 | DNPH1 |
| ZNF574 | SH3KBP1 | DEGS1 | DHRS2 | KICS2 |
| POLDIP3 | AGK | DENR | ACSBG1 | BAZ2B |
| TOMM70 | RAB31 | DEXI | CYP4F3 | BACE1 |
| TCHH | NEU2 | DFFB | ZNF702P | BAG5 |
| POLR3A | DNM2 | DGAT1 | AKR1B10 | MAGI1 |
| WFDC21P | HBP1 | DGAT2 | CEACAM1 | DDX39B |
| LDHB | SH3KBP1 | DGKA | ADGRG1 | BBS2 |
| NPEPL1 | ACKR1 | DGKQ | APOBEC3B | BBX |
| TJP2 | AK3 | DMPK | IL7 | BCCIP |
| PLEC | UBAP2L | DNAJA1 | GPR143 | BCKDHB |
| HSD17B11 | WASHC2A | DNAJB1 | EDNRB | BCKDK |
| SAP18 | CHCHD10 | DNAJB2 | EPB41L3 | BCL2 |
| EXTL3 | JAK1 | DNAJB9 | WNT11 | BCL2L2 |
| MLLT1 | KRT77 | DNAJC9 | HOXC5 | BCL3 |
| PKIG | KRTDAP | DNM2 | EDN2 | BCL7B |
| COPS9 | AGPAT5 | DOK4 | HOXC6 | BCL9 |
| RPL37 | S100B | CBARP | BGLAP | BCLAF1 |
| RBBP4 | USP34 | DPP7 | HSD17B3 | BCR |
| MRTFA | CPD | DRAP1 | FGF18 | BCS1L |
| NEO1 | SPPL2B | DRG2 | TSC22D3 | BET1 |
| CACFD1 | TMEM106C | RCAN1 | EN2 | BGN |
| UNC5B | PLA2G2F | DSTN | SMAD6 | BICC1 |
| ATP5PD | DGCR2 | USP17L2 | GAB1 | BICD1 |
| CELSR2 | FCGR2B | USP17L2 | GGT1 | BIN1 |
| ATP6V0C | BTC | DUSP6 | CITED1 | XIAP |
| DUS1L | UBB | DUSP9 | KRT32 | BIRC6 |
| SLC5A6 | CDC42BPA | SLC66A3 | FRK | BMPR1A |
| PCBP2 | CENPB | LRRC14 | DNASE1L2 | BNIP1 |
| RHBDL3 | RSAD2 | E2F1 | CNGA3 | BNIP3 |
| HR | APOA4 | ZNF764 | GALNT10 | BNIP3L |
| NAV2 | SOX11 | PDE12 | DGKZ | BOC |
| MBP | RAB2B | EAF2 | MR1 | BPNT1 |
| DSTYK | TMEM109 | ECE1 | THEMIS2 | BRAF |
| PGD | PIGO | ECM1 | BLNK | BRCA2 |
| MAN2B1 | DVL1 | EDEM1 | ZNF304 | BRD4 |
| ILF3 | RARRES2 | EED | FDXR | BRD8 |
| SCYL1 | EPHA5 | EFNA1 | FLNC | BABAM2 |
| INTS9 | CCN1 | EFNB1 | MCF2 | BRI3 |

| | | | | |
|---|---|---|---|---|
| RPL31 | TPP2 | EFNB2 | CBFA2T3 | MPC1 |
| CLNS1A | PTTG1 | EFNB3 | CYRIA | NCAPH |
| XIST | DNPEP | EGR1 | COL21A1 | CLCF1 |
| RPL26 | EIF4G1 | EGR2 | MAN1A1 | BTBD3 |
| SNX30 | BRAT1 | EHD1 | EPPK1 | ZBTB46 |
| PCOLCE | DPYSL3 | EHD4 | FGF1 | BTC |
| LIG3 | ENPP3 | EI24 | GGT1 | BYSL |
| BRD2 | DNPEP | EIF1AX | ANXA13 | C1QBP |
| MARCKSL1 | ARL5A | EIF2B2 | MEF2A | PEAK1 |
| IAPP | MSR1 | EIF3B | CSH1 | IGF2BP2 |
| PTTG1IP | SLC25A48 | EIF4E3 | ANK1 | CIP2A |
| PGD | RRM1 | EIF4EBP1 | ANK1 | NT5C3B |
| IGFBP4 | TNFRSF19 | ELF3 | C4A | FANCM |
| RTRAF | IGF2 | ELOVL7 | GABRA4 | URI1 |
| CLU | LCP1 | EMP3 | CFLAR | EFR3A |
| AMMECR1L | PRPS1 | ENDOG | MC5R | CAB39L |
| FUBP1 | LASP1 | ENO2 | WNT4 | COQ8A |
| RPS8 | GPD1 | ENTPD7 | TSC22D3 | CACNA1A |
| PPIB | TMED9 | EPHA2 | ALDH7A1 | CALM3 |
| UBC | KLHL13 | EPHX1 | CRIP2 | CAMK2D |
| BACH2 | SERPINB1 | EPM2AIP1 | KDM2A | CAMK2G |
| CLU | AP3M1 | ERBB3 | KRT8 | CAP1 |
| CPT1A | CAPRIN1 | ERCC5 | CADM1 | CAPN6 |
| NFIB | ASB6 | ESRRA | CADM1 | CAPZA1 |
| NFIB | FAM107B | ETFDH | CADM1 | CA9 |
| FASTKD2 | LBP | EVI2A | COL18A1 | CARM1 |
| CLCN3 | BID | EXOC8 | COL18A1 | CASC3 |
| RPL14 | NCF2 | EXOSC9 | DDAH1 | CASK |
| TOR2A | PBK | EXTL1 | CDKN1B | CASP2 |
| ZDHHC14 | CPA3 | F11R | PLIN2 | CAT |
| PGD | CP | FA2H | MAN2B1 | CTNNB1 |
| PIGQ | IFI16 | FAH | ADCY6 | CTNNBIP1 |
| ZDHHC14 | TBRG4 | FAS | KIF1B | CAV1 |
| RPS17 | CYP11A1 | FBXO2 | CRYAB | CAV2 |
| FHDC1 | USF1 | FBXO33 | ID4 | CBL |
| TC2N | CXCL12 | FBXW4 | ULK1 | CBX1 |
| EIF3E | DUSP1 | FBXW9 | DEAF1 | CBX3 |
| PIGU | MSRA | FCGR3A | LRP5 | CBX5 |
| NHERF2 | LTBP1 | FDXR | USP15 | CBX6 |
| HMGB1 | AKR1B10 | FETUB | RBM4B | CBX8 |
| KRTAP5-4 | CCL15 | FEZ1 | CEACAM1 | HAUS1 |

| | | | | |
|---|---|---|---|---|
| KAT7 | SELENOF | FGF18 | CACNB3 | CCDC9 |
| UCP2 | KRT16 | FGFR4 | ARHGEF6 | CCL5 |
| MRPL33 | CA4 | FHL2 | HIP1R | CCNH |
| PLK1 | DUSP22 | FKBP9 | DLK1 | CCNI |
| SFRP2 | MFAP5 | FLCN | NSG1 | CCNT2 |
| RPL27 | FBP2 | FOLR1 | NSG1 | CD200 |
| CCND2 | CTF1 | FOS | LDLRAD4 | CD276 |
| PRNP | CCDC43 | FOSB | PLAAT3 | CD2AP |
| HMGB1 | FECH | FOSL1 | BMP7 | CD44 |
| RAB6A | RETN | FOXJ1 | BMP7 | CDC27 |
| G6PD | BAG4 | FOXQ1 | MXRA5 | CDC42BPA |
| SNRPG | PDGFA | ALDH1L1 | GATA3 | CDC6 |
| LGALS3BP | KIF20A | FTSJ3 | GATA3 | CDC7 |
| GJB6 | SLC22A5 | FUCA1 | GATA3 | PIGU |
| SMARCD2 | SKP2 | FYTTD1 | CES1 | NUF2 |
| KRT71 | FADS2 | FZD5 | GPR37 | CDCA7 |
| FBP1 | STOM | FZD7 | CYB561D2 | CDK2 |
| GGT1 | ATF3 | ISG15 | CES2 | CDK4 |
| GAS1 | TREX2 | TRA2A | CES2 | CDK5RAP1 |
| EPHX2 | HOXB2 | CENPT | DOK4 | CDK5RAP2 |
| RASSF3 | PIP4K2A | GADD45A | MOXD1 | CDKAL1 |
| UPP1 | PLEK2 | GAL3ST1 | GATA2 | SLC44A1 |
| HIPK2 | CA3 | GALM | ABCG2 | CEBPB |
| CAMK2B | CMA1 | GALNT3 | JAG2 | CEBPG |
| UBQLN4 | RBP7 | GATA3 | CASP6 | AGAP1 |
| SLC35B1 | SLC25A13 | GBP2 | CHST3 | CETN3 |
| ATP5IF1 | ZFR | GBP6 | FUT4 | CGNL1 |
| ALDH1A3 | KLB | GCC1 | FUT4 | CHCHD3 |
| GOLGA4 | HDLBP | GCDH | ITSN2 | CHD1 |
| WNT5A | PTGFR | GCH1 | ITSN2 | CHD1L |
| MRPL39 | SCUBE1 | GCNT2 | GGT1 | CHD4 |
| RHOU | LCE1F | GDF15 | CAMK2B | CHD7 |
| RAB4A | DLK1 | GEM | CSDC2 | CHEK1 |
| ZNF148 | SEC61A2 | GFER | ASCL1 | CHEK2 |
| SPRR1A | CAB39L | GGA2 | GABBR2 | CHM |
| RTP4 | CACNA2D1 | GGTA1 | ABCB1 | CHML |
| CHST1 | TSLP | GKAP1 | TAF5 | CHST10 |
| S100A14 | NCOA4 | GLCCI1 | MAPK13 | VSX2 |
| PIWIL2 | ITGA9 | GLUD1 | MAPK13 | CLASP2 |
| RDH11 | CFAP97 | GLUL | MARCHF2 | CLK4 |
| PIP | ANGPTL2 | GNA13 | ANXA9 | CLOCK |

| | | | | |
|---|---|---|---|---|
| ACTRT2 | GNA12 | GNA15 | HR | CLPP |
| ABCA2 | PTPRG | GNL2 | ARC | CLSTN1 |
| BACH1 | ITGA4 | GNS | IGFBP3 | CLYBL |
| ERO1A | FUT2 | GORASP1 | KCNJ15 | CNN3 |
| TIMP3 | CLCF1 | GPAA1 | CHRNA7 | CNOT1 |
| HACD2 | TNFSF14 | GPD1 | LTB4R | CNOT2 |
| AURKC | ASB7 | GPHA2 | HIPK3 | CNOT4 |
| KRT27 | THBS1 | GPLD1 | SEC14L5 | CNOT6 |
| KRT33A | SLC48A1 | GPR146 | MR1 | CNOT6L |
| RPL3L | CAPN6 | TPRA1 | IRX5 | CNOT7 |
| DLX1 | PCYT1A | GPSM1 | WNT7A | CNTF |
| GDPD3 | HCFC1 | TAMALIN | MSX2 | CNTN1 |
| MSX2 | TCF7 | GRINA | GNMT | COASY |
| KRTAP1-4 | EPS8L1 | GSN | MYT1 | COG1 |
| CYP2G1P | CEACAM1 | GSS | DEFB1 | COIL |
| GSTP1 | POU2F3 | GSTA2 | CAMK2B | COL12A1 |
| SIAH1 | CYSRT1 | GSTO1 | GOLGA8A | COL1A2 |
| PKP2 | VN1R17P | GSTZ1 | GOLGA8A | COL3A1 |
| ACAN | ACSL1 | GTF2A1 | CYP4F2 | CERT1 |
| SPESP1 | TYROBP | GTF2B | MYO6 | COL4A6 |
| EPAS1 | TNNI1 | GTF2F2 | LIMK2 | COL5A2 |
| GIP | SAA3P | GTF2IRD2B | TP53I3 | COMMD1 |
| KRTAP1-3 | BZW1 | NATD1 | THEMIS2 | COMMD2 |
| CX3CR1 | LIPA | GTPBP1 | AHSG | COPG2 |
| FZD7 | CD36 | GTSE1 | CAPN3 | COPS7A |
| CER1 | TGFB2 | GUSB | AKAP9 | COQ3 |
| UBE2F | MAGOHB | HLA-DRB5 | KCNB1 | ENOX2 |
| PDE3A | TCEAL3 | SELENOS | FGF18 | COX17 |
| MGLL | NDRG1 | HAGH | HECTD4 | CPNE1 |
| TGS1 | ITM2A | HAP1 | CROCCP2 | NUDT21 |
| YIPF4 | BCAP31 | HAS2 | USH1C | CPT1B |
| HNF4G | CMTR1 | HBA2 | GALR2 | CRADD |
| SF3A2 | ETFDH | HBEGF | GRN | CREB1 |
| MT4 | FBLN1 | HERC1 | CFLAR | CRELD1 |
| MYO10 | PINK1 | HES6 | COL13A1 | EID1 |
| GNA13 | ZNF410 | HEXB | CASP1 | CRK |
| RIMS2 | CEP85 | H1-2 | GGT1 | CRYZL1 |
| LY6G6D | ZNF330 | H2BC10 | CASP6 | YBX3 |
| TG | GLO1 | H3C14 | CAMK2B | CSNK1D |
| CENPA | ZNF521 | H3C14 | GH2 | CSNK2A1 |
| PPP4R3B | SPON1 | H2AC25 | DSTYK | CSTF2 |

| | | | | |
|---|---|---|---|---|
| TEX261 | TM6SF1 | HMG20B | HPGD | CSTF3 |
| HACD4 | MRAP | HMGA1 | APAF1 | CTBS |
| TJP2 | KDM8 | INO80B | APAF1 | CTCF |
| LGR5 | FAM162A | HMOX1 | VCAN | CTDSP1 |
| XDH | PCCA | JPT1 | MDFIC | CTDSP2 |
| GSK3B | TCIM | HOXA5 | ANXA9 | CTDSPL |
| AARS1 | USHBP1 | HPN | BID | CTTN |
| DAP | EIF4H | HR | GINS4 | CUL2 |
| IDI1 | IFI16 | HRAS | EPB41L3 | CUL3 |
| GTF2B | HAVCR2 | PLAAT3 | MDM2 | CUL4B |
| LSR | HOXB6 | RIDA | CEACAM1 | CXADR |
| ERC1 | WDR45 | HS6ST1 | CEACAM1 | CXCL6 |
| CHD4 | NCF1 | HSBP1 | CAPN3 | CXXC1 |
| CHAC1 | HDC | HSD11B2 | ARHGEF4 | CLIP2 |
| GABRP | SPTBN1 | HSD17B4 | TRAF4 | CYP1B1 |
| CUX1 | GHR | HSPA1B | MERTK | CYP24A1 |
| HJURP | SMAD5 | TMEM132A | EXOC7 | CYP2C18 |
| XPO5 | TP63 | HYAL1 | GSE1 | CYP4F3 |
| MIF4GD | RFC1 | HYAL2 | GSE1 | CYP51A1 |
| EGFR | SIGMAR1 | RNF19B | PRRC2B | AHDC1 |
| SOX9 | MAPRE2 | ICAM5 | PRRC2B | AMBRA1 |
| SSBP2 | ALOX5AP | ICMT | ADGRG1 | NREP |
| CIBAR1 | MGAT2 | IDH1 | RHOB | MRPL42 |
| EFNB1 | DCUN1D1 | IDUA | IGFBP3 | GATD3 |
| CES4A | MISP | IER5 | GALNT10 | CCDC43 |
| ANK3 | ZBED3 | IRGM | TXLNA | GHDC |
| WDR75 | ACBD3 | IFI16 | CREB3L2 | FAM104A |
| AP4S1 | WDR48 | IFI16 | EEIG1 | SOCS2 |
| PLXNA2 | SWSAP1 | IFI30 | PCMTD2 | RNF214 |
| KRTAP4-1 | TRBC2 | IFI35 | ZCCHC24 | CCDC50 |
| HOPX | MORC3 | IFI44 | ZCCHC24 | MYDGF |
| CRTC3 | IFI16 | IFIH1 | KIAA0232 | PUM3 |
| HLA-B | TMEM191B | IFIT1B | FYN | TUBGCP4 |
| TMCC3 | CIDEC | IFIT3 | KDM4B | KNSTRN |
| KRTAP6-3 | PTPRZ1 | IFNGR2 | KDM4B | SUPT20H |
| TENM4 | TMEM106B | IGFBP4 | KDM4B | CSDE1 |
| PARP3 | AP2B1 | IGSF8 | ADO | RUFY3 |
| FRMD4B | PMEPA1 | IRGM | ADO | SEPSECS |
| THRAP3 | MYO5B | IRGM | DIP2C | COQ5 |
| PTPN13 | KIF11 | IK | DIP2C | MEPCE |
| MAML1 | INTS12 | IKBKE | MOAP1 | CENATAC |

| | | | | |
|---|---|---|---|---|
| RCC2 | HACD3 | IL15 | ARAP1 | OPA3 |
| SPTBN2 | B2M | IL17D | WEE1 | FAM3C |
| IFI16 | HLA-DQA1 | IL18 | MEF2A | TCF25 |
| BCR | ART3 | IL24 | SPRY1 | C19orf53 |
| ZFP36 | CD44 | IL4R | AUTS2 | DAP |
| H2BC5 | CTBS | IL6R | ZCCHC14 | DARS1 |
| SAFB2 | ELOC | IMPA1 | CAMK2G | DAZAP1 |
| MBD3L2B | TP53INP2 | IMPACT | EPB41L3 | DBN1 |
| RNASET2 | MYL1 | INCENP | RETREG3 | DCLK1 |
| C9orf64 | TMEM222 | INHBB | MICAL3 | ECI1 |
| NDUFA3 | NAP1L1 | INPP5B | SYNM | DCPS |
| SCAF11 | ITGAV | INPP5D | ANKRD46 | DCTN4 |
| RPS24 | PRKCE | IRAK1BP1 | ARHGAP19 | ASAP1 |
| STX18 | GNA13 | IRF6 | MON2 | DDR2 |
| MGLL | DYNLT1 | IRF7 | CAMK2G | DDX10 |
| GRK4 | HBS1L | ISG20 | GRAMD4 | DDX17 |
| IGFBP2 | SERPINF1 | ISLR | ABR | DDX19B |
| DPYSL3 | DRAM2 | ITGA2B | GRAMD1B | INTS6 |
| MBP | CCNL2 | ITGA6 | MCF2L | DDX3X |
| THRA | CAPZB | ITGB4 | GNPTAB | DDX6 |
| ESRP1 | IDE | ITM2C | HIC2 | ADAT2 |
| TMED10 | SRPK1 | ITPA | HIC2 | DEK |
| CLU | ERI2 | ITPKA | ACKR3 | DEPTOR |
| RPL23 | RBBP4 | IVD | AHSA2P | DGCR2 |
| SRSF3 | PIP5K1A | JAK3 | HRAS | DGCR8 |
| TNRC6A | LGALS1 | KDM5B | AHNAK2 | DHCR24 |
| LDHB | UBE2D3 | JUNB | URB1 | DHCR7 |
| RPS7 | HOXA9 | JUP | ATPAF2 | DHDDS |
| LASP1 | PAPOLA | KCNK2 | HEG1 | DHODH |
| RPS18 | TUBGCP2 | KCTD10 | TMCC2 | DHPS |
| RPS17 | CHD9 | KCTD11 | TSPYL5 | DHRS4 |
| ECE1 | SNX17 | KCTD12 | ZBTB20 | DHX36 |
| RPL21 | HADH | KLHL22 | ZBTB20 | DHX9 |
| CCND2 | UQCRB | KIF17 | CDKN1C | DIABLO |
| GCLC | FERMT3 | KIF22 | GOLGA6L9 | DIAPH3 |
| ADAMTS4 | SERINC3 | KIFC3 | CEP170B | DLAT |
| SF3B2 | FUCA1 | KLC2 | VPS13B | DLGAP5 |
| MBP | RANBP9 | KLF5 | ADCY1 | DLG1 |
| MCL1 | TGFBI | KLHL42 | MARCHF3 | DLX5 |
| TSPO | ESYT3 | KLHL10 | CAMTA1 | DMP1 |
| FAAP20 | ZNF574 | KRT14 | ZNF248 | DNA2 |

| | | | | |
|---|---|---|---|---|
| CDK2AP1 | PROKR1 | KRT5 | DOP1A | DNAJB12 |
| ATP11A | PFKFB3 | TRIM5 | CLIC5 | DNAJC13 |
| BBIP1 | ZSCAN26 | MSANTD5 | CDKN1C | DNAJC14 |
| C1R | SPIN1 | CPOX | MTCL1 | DNM1L |
| SP110 | HADH | LACTB | CDHR1 | TRDMT1 |
| UAP1 | YKT6 | LANCL1 | MST1P2 | DOCK1 |
| ZNF280D | CA3 | LASP1 | MFSD9 | DOCK5 |
| C22orf39 | CLCA3P | LBX2 | MLC1 | DOCK9 |
| KRT85 | IL6ST | LCMT1 | MFSD9 | REEP6 |
| PKD1 | CSRP3 | LDB3 | COL6A1 | DPAGT1 |
| GOLGA4 | PLGRKT | LDLRAP1 | MFHAS1 | DRG1 |
| TIMP1 | INPPL1 | LENG1 | PGAP1 | RCAN3 |
| KLK10 | PDE4DIP | LGALS3BP | KAZN | PSMG1 |
| BRD3 | RABGAP1 | LGALS9 | COL2A1 | PIGP |
| RIOK3 | NABP1 | LHX2 | ABTB2 | DSG2 |
| KMT5A | TRIAP1 | LIF | SPTSSA | DST |
| COL9A3 | AP2A1 | LIG3 | CES2 | DTNBP1 |
| MRPL52 | PC | LITAF | FAM106A | DTX3 |
| DNPH1 | PC | LMAN2 | NSG1 | DTYMK |
| | PC | LMO4 | ATP6V0E2 | DUSP11 |
| | TFRC | ADGRL1 | ICAM2 | DUSP12 |
| | ACTB | LPIN1 | INPP5B | DUSP16 |
| | | LRFN1 | GLB1L2 | DUSP19 |
| | | LRRC8A | CACNB2 | DUSP22 |
| | | PTGR1 | GOLGA8N | DYM |
| | | LTBP2 | GNA11 | DYRK1A |
| | | LY6D | INSR | DYSF |
| | | LYNX1 | INPP5B | BBS9 |
| | | LZTR1 | ADAM23 | DENND11 |
| | | ACAA1 | HOXA11 | DAGLB |
| | | ARPP21 | HOXA5 | CBX1 |
| | | G3BP1 | TRIM3 | RCCD1 |
| | | SIVA1 | TRIM3 | LRRC8C |
| | | GIPC2 | ZNF20 | EBPL |
| | | GDE1 | GNA11 | EEA1 |
| | | ROGDI | TBC1D31 | EEF1E1 |
| | | ZBTB8OS | DHRS2 | EEF2K |
| | | SHISA5 | CEP43 | EFHD1 |
| | | ISYNA1 | MAN1C1 | EFHD2 |
| | | TRAFD1 | CSPG4 | EFNA4 |
| | | THYN1 | ATPAF2 | EFNA5 |

| | | |
|---|---|---|
| SAP30BP | MUC5AC | EFL1 |
| CEP295NL | C4A | DLK2 |
| HIGD1A | HSPA12A | EGLN1 |
| CHCHD10 | CAPN3 | EGLN2 |
| HEXIM1 | GPR37 | EHD2 |
| ZNF703 | TP53I11 | EIF2S2 |
| CNTRL | ABR | EIF4A1 |
| MAFB | GNA11 | EIF4A2 |
| MAFG | MEF2A | EIF4B |
| MAFK | ZNF468 | EIF4E2 |
| NAA35 | CLCN4 | EIF5B |
| MAL2 | RIMBP2 | CELA1 |
| MAN2B1 | BRWD1 | ELAC1 |
| MAP1LC3A | TMEM63A | ELAC2 |
| MAP2K3 | LRP5L | ELF5 |
| MAPK9 | ZNF37A | ELK1 |
| MAPKAPK2 | FAM149A | ELK3 |
| MAPKBP1 | FAM149A | ELK4 |
| MAPRE1 | SUSD5 | ELMO1 |
| MAPRE3 | MTUS2 | ELOVL5 |
| MARK4 | SPATA2L | ELOVL6 |
| MARVELD1 | LINC00667 | ELP3 |
| MASP1 | BRD1 | ELP4 |
| MCEE | ABCA12 | EMP1 |
| MCFD2 | AKAP9 | EMP2 |
| MCL1 | LAMB4 | ENC1 |
| CLCC1 | GGT1 | ENSA |
| MDH1 | VCAN | EPB41L2 |
| MDM2 | FLG | EPB41L4A |
| MED28 | WEE1 | EPC1 |
| MFGE8 | TSC2 | EPHA4 |
| MFRP | CYFIP2 | EPHB4 |
| MGAT2 | DUOX1 | EPN2 |
| MID2 | NPIPB3 | ERBB2 |
| MIDN | ARFGEF2 | ERBIN |
| MITF | ENSG00000237 | ERCC4 |
| MLF2 | GRN | ETF1 |
| MLH3 | FADS3 | ETS1 |
| MARCKSL1 | ATP2B2 | EWSR1 |
| MMD | GRM8 | EXOSC1 |
| MMP15 | MST1 | EXOSC10 |

| | | |
|---|---|---|
| IFI16 | BAIAP3 | EXOSC2 |
| MOAP1 | LTB4R | EXOSC5 |
| GBP6 | CDKN1C | EXOSC7 |
| MPP4 | MCF2 | EXT1 |
| MR1 | GNAS | EZH1 |
| MRPL28 | MDM2 | F2RL1 |
| MRPS25 | COL2A1 | HMBOX1 |
| MSH6 | LIMK2 | FAAH |
| MTHFR | C18orf25 | FADS1 |
| MUSTN1 | MDM2 | FADS2 |
| MX1 | MDFIC | FAIM |
| MXD3 | WNT7B | FANCA |
| MXD4 | CYBRD1 | FANCC |
| MYCBP | BACE1 | FANCL |
| PPP1R15A | CXCL14 | FARP1 |
| MYD88 | ZNF672 | FARS2 |
| MYO1H | COQ8A | FARSB |
| MYO5A | ISOC1 | FASTK |
| NANOS1 | EPS8L2 | FBF1 |
| NAPA | CDIP1 | FBXL8 |
| NAA80 | FYCO1 | FBXO21 |
| NBEAL2 | SMAD3 | FBXO4 |
| NCOA1 | MINDY3 | FBXO6 |
| NCOA4 | AKTIP | FBXW11 |
| NDRG4 | NLRP1 | FBXW2 |
| NDUFA11 | GMCL1 | FKTN |
| NFATC2 | BBS1 | FDFT1 |
| NFKBIE | HILPDA | FDPS |
| NFYB | RETREG1 | FECH |
| NID2 | RETREG1 | MYOF |
| NINJ1 | UCKL1 | FER |
| NINJ2 | FBXO34 | FEZ2 |
| NMI | MAFB | FGFR1 |
| NMNAT1 | MAGEH1 | FIGNL1 |
| ATG9B | ABHD4 | GSDME |
| NPC2 | POGLUT1 | FJX1 |
| SRXN1 | ZXDC | FKBP11 |
| NPTX1 | CHCHD7 | FLOT1 |
| NR4A1 | VWA1 | FMO1 |
| NUDCD2 | MSL2 | FMR1 |
| NXN | ANKRA2 | FNDC4 |

| | | |
|---|---|---|
| OAS1 | MED23 | FOSL2 |
| OAS1 | LIMD1 | FOXC1 |
| OAS1 | MOSPD1 | FOXF1 |
| OAS1 | GALNT12 | FOXF2 |
| OAS2 | CNNM4 | FOXJ3 |
| OAS3 | ASPSCR1 | FOXM1 |
| OASL | MAN1C1 | FOXP1 |
| OASL2P | CERS4 | FRMD4B |
| OGFRL1 | PLCXD1 | FRS3 |
| OMP | KRT23 | FSTL1 |
| OPLAH | INAVA | FUBP1 |
| OPN1SW | GALNT11 | SRSF10 |
| OTUB2 | PIDD1 | FUT8 |
| OVOL1 | PARP16 | FXN |
| P2RY6 | HPS6 | FYN |
| PA2G4 | TINAGL1 | FZD2 |
| PADI1 | ITIH5 | FZR1 |
| PADI2 | TMEM184C | ZNF771 |
| PADI3 | DACT1 | GAB1 |
| PAK1IP1 | MACROD1 | GABBR1 |
| PAQR4 | IFIH1 | GABPA |
| PARG | USP18 | GABRB2 |
| SLC45A3 | ZNF654 | GALK1 |
| CDK18 | SH3TC1 | GALNT2 |
| PCTP | INTS15 | GALNT4 |
| PCYOX1 | CHAC1 | GALNT7 |
| PDCD10 | TRPV2 | RALGAPA1 |
| PDCD6IP | ECHDC3 | GART |
| PDE1B | TMEM204 | GAS5 |
| PDGFA | WRAP73 | GATAD2B |
| PDGFB | GPRC5C | KAT2A |
| PDK4 | MICALL2 | GCNT1 |
| PDLIM1 | CLN8 | GCSH |
| PDLIM5 | ADAP2 | GDI1 |
| PDLIM7 | PRR5L | GDPD1 |
| PDPK1 | GRHL2 | GEMIN6 |
| PDRG1 | ESRP2 | GHR |
| PDXP | FA2H | GJC1 |
| PERP | ARHGAP10 | GLG1 |
| PEX26 | BCOR | GLI2 |
| PEX6 | TMEM40 | GMFB |

| | | |
|---|---|---|
| PFKFB2 | ELL3 | GMNN |
| PFN2 | SLC47A1 | GMPR |
| CBY1 | CDKN1C | GNL1 |
| PGF | SYNE3 | GNA12 |
| PGRMC1 | SLC35G2 | GNAO1 |
| JADE1 | ABCA7 | GNB1 |
| PHGDH | DUOX1 | GNG12 |
| PHLDA2 | ZNF767P | GNG5 |
| PHLDA3 | RTP4 | GNPNAT1 |
| PIAS3 | TMEM35A | GOLPH3 |
| PIGF | OTULINL | GOLIM4 |
| PITPNC1 | APOL6 | GORASP2 |
| PITPNM1 | GPATCH2L | GOT1 |
| PITPNM2 | AGPAT3 | GPAM |
| PIWIL2 | TREM2 | GPHN |
| PKP1 | DUOX2 | CAPRIN1 |
| PKP3 | ENTPD1-AS1 | GRK5 |
| PLA2G12A | PLPPR1 | GPT2 |
| PLA2R1 | SIDT1 | GRB14 |
| PLA2G6 | TRIM36 | EMG1 |
| PLAT | VASH2 | GRTP1 |
| PLCB3 | RBM41 | GSK3B |
| PLEC | HOXC13 | GSR |
| PLEK2 | GASK1B | GTF2I |
| PLEKHA3 | LHX6 | GTF2IRD1 |
| PLEKHA4 | MMP28 | GTPBP2 |
| PLK2 | LTBP3 | GYS1 |
| PLK3 | CABYR | GYS1 |
| PLS3 | ADAMTS5 | H1-0 |
| PLXNB2 | DUSP13 | HLA-G |
| PML | ZNF750 | HACE1 |
| PMM1 | ERO1B | HADH |
| PMPCA | LMBR1L | HARS2 |
| PNP | EDAR | HBP1 |
| PNPLA2 | FOXL2 | HCFC1 |
| PNPT1 | ZC3HAV1 | HDAC3 |
| POLE3 | GNA14 | HDAC7 |
| POLK | HAND1 | HDHD3 |
| POLR2A | FAM184A | HECTD1 |
| POLR3D | EPB41L4B | HELLS |
| POLRMT | LRRC8E | HERC4 |

| | | |
|---|---|---|
| POM121 | ZNF215 | HIBCH |
| PPIF | LRRC37A | HIF1A |
| PPM1A | LRRC37A4P | HIP1 |
| PPM1B | SMAD5-AS1 | UBE2K |
| PPM1M | CRYBG2 | HIPK2 |
| PPP2R5D | TENT5C | H1-4 |
| PPP5C | MYLIP | HIVEP2 |
| SLC66A2 | CES3 | ZSCAN22 |
| PRCP | ELMO2 | HLF |
| PRKCD | PODNL1 | HMBS |
| PRKD2 | KCNK7 | HMGA2 |
| PRKRA | SLC30A10 | HMGB1 |
| PRKRIP1 | PGAP1 | HMGB2 |
| PROCR | TSGA10 | HMGXB4 |
| PLPBP | DNMT3B | HMGCS1 |
| PRSS8 | KCNK10 | HNF4A |
| PSMB4 | CNNM3 | HNRNPA1L2 |
| PSMB9 | FAXDC2 | HNRNPD |
| PSMC3IP | CA10 | HNRNPDL |
| PSMF1 | C1QTNF1 | HNRNPK |
| PSTPIP1 | TSPAN14 | HNRNPL |
| PSTPIP2 | LBH | HNRNPR |
| PTGER1 | TRIM8 | HNRNPU |
| PTGER4 | C17orf80 | HOMER2 |
| PTHLH | PAXBP1 | HP1BP3 |
| TWF2 | CIDEB | AGFG2 |
| PTP4A1 | MSANTD2 | PRMT2 |
| HACD2 | RIPK4 | CDC73 |
| PTPN1 | KLHL3 | HSD11B1 |
| PTPN11 | TGFBRAP1 | HSD3B7 |
| PTPRM | GDF9 | HSPA4 |
| PTPRVP | S1PR5 | HSPA8 |
| PURG | NPIPA1 | HSPA9 |
| PVALB | CRISPLD2 | HSPB8 |
| PYGO2 | ABHD6 | HSPE1 |
| PPA1 | LAT2 | HTATSF1 |
| QSOX1 | PIDD1 | HUWE1 |
| RAB11FIP5 | ZDHHC11 | IARS1 |
| RAB40C | APOL2 | IBTK |
| RABGAP1 | LHX3 | CILK1 |
| RABGEF1 | ABHD6 | MRPL58 |

| | | |
|---|---|---|
| IFT27 | SIKE1 | ID3 |
| RDM1 | MAP6D1 | IDE |
| RAD9A | COL5A2 | IER3 |
| RALGPS1 | VCAN | IFRD2 |
| RAP1GAP | TNS1 | IGF1 |
| RAP2B | TNS1 | IGF1R |
| RASA4 | MAN1A1 | IGFBP2 |
| RASSF5 | MARCHF8 | IGFBP5 |
| RB1 | LONP2 | CADM1 |
| RBL2 | MINDY1 | IP6K1 |
| MAK16 | ARRB1 | IKBKB |
| RBM15 | WHRN | IKBKG |
| RBM4 | TRIM52 | IL13RA1 |
| RCN2 | SHISAL1 | IL15RA |
| RDH10 | FBXO41 | IL17RC |
| RDH5 | EMX2 | IL1R1 |
| RECQL4 | LOC100505915 | IL6ST |
| REN | MYMX | ILF3 |
| REN | FAM149A | ILKAP |
| RFNG | SCAF4 | IMMT |
| RGL1 | TGS1 | INSIG2 |
| RGN | ZDHHC9 | INVS |
| RGS16 | CYBRD1 | IPO11 |
| RHBG | BACE1 | IPO8 |
| RHOD | TOLLIP | IPP |
| RHPN2 | UBE2Q1 | IQCC |
| RIPK4 | CXCL14 | IRF3 |
| RNF103 | ZNF672 | ISOC1 |
| RNF128 | ARFGEF2 | ITCH |
| RNF135 | EPS8L2 | ITGA5 |
| RNF14 | ASH1L | ITGB3BP |
| RNF4 | MAFB | ITPR1 |
| RBM38 | CHCHD7 | IVNS1ABP |
| RNPC3 | ARRB1 | JARID2 |
| RNPEP | LIMD1 | KDM4B |
| RRP9 | GALNT12 | JTB |
| ROM1 | TNRC6C | KCNK1 |
| RPE | ZNF654 | KCTD1 |
| RPS27 | CLN8 | KDELR1 |
| RPUSD1 | ADAP2 | KEAP1 |
| RRAGC | TMEM127 | KHDRBS1 |

| | | |
|---|---|---|
| RSAD2 | EXO5 | KIF11 |
| RUSC1 | ARRB1 | KIF1B |
| RXRB | CXXC5 | KIF2A |
| RXRG | MYLIP | KIF3C |
| S100A1 | MYLIP | KIF4A |
| S100A13 | TRIM8 | KIF5B |
| S100A3 | TRIM8 | KITLG |
| SAC3D1 | YPEL3 | KLF13 |
| SAP18 | AGPAT3 | KLF3 |
| SAP30 | AGPAT3 | KLF6 |
| SBK1 | AGPAT3 | KLF7 |
| SCARB1 | SLC22A23 | KLHL20 |
| SCIN | TLNRD1 | KLHL5 |
| SCMH1 | POLR3GL | KLHL7 |
| SCN1B | ZNF655 | KIF2A |
| SCNN1A | TSPAN14 | KPNA1 |
| ERGIC3 | CCDC3 | KPNA3 |
| SDC1 | C17orf80 | KPNB1 |
| SORD | C17orf80 | KRAS |
| SEC61A1 | GLIS2 | KRTCAP2 |
| EXOC4 | CYP2S1 | LACTB2 |
| SEPHS2 | ZFYVE1 | LAMC1 |
| SEPTIN3 | ZFYVE1 | LANCL2 |
| MSRB1 | EPB41L4B | LBR |
| SERPINB5 | IRF2BPL | LDLR |
| SERPINE2 | OGA | LETMD1 |
| SESN2 | SPRTN | LGALS7 |
| SF3A1 | FBXO44 | LIG3 |
| SF3B5 | NECTIN4 | LIMK1 |
| CLASRP | ESPN | LIMS1 |
| SGCA | CA10 | LMBR1 |
| SGK2 | BCOR | LMNB1 |
| SGPL1 | HHATL | LMNB2 |
| SGPP1 | KIAA1549 | BORCS5 |
| CTR9 | ADGRV1 | LOX |
| SH2D3C | ZNF649 | LOXL4 |
| SH3BGRL2 | MEX3B | LPP |
| SH3BGRL3 | POLR1H | LRBA |
| SH3GLB2 | ZIC2 | LRCH4 |
| SH3YL1 | TRIM7 | LRP6 |
| SHMT2 | WNT10A | CARMIL1 |

| | | |
|---|---|---|
| SIAH1 | TTYH2 | LSS |
| SIAH1 | C2orf88 | LTA4H |
| ST3GAL2 | SUN1 | VANGL2 |
| ST6GALNAC4 | BVES | LTBP1 |
| SIDT2 | AKTIP | LUC7L2 |
| SIRT1 | CLN8 | LCLAT1 |
| SIRT7 | CDIP1 | PLA2G15 |
| SKP1 | KCNK7 | LYPLAL1 |
| SLC12A2 | KCNK7 | LYST |
| SLC12A4 | IL20 | LZTS2 |
| SLC12A7 | ABCC11 | PLIN3 |
| SLC12A9 | ETV7 | HEXD |
| SLC18A1 | CABYR | INPP5K |
| SLC19A2 | AGPAT3 | SWAP70 |
| SLC20A1 | EMILIN2 | FTO |
| SLC25A22 | TRPM6 | SERGEF |
| SLC25A25 | AIFM2 | TM9SF3 |
| SLC25A29 | MFSD9 | AHCTF1 |
| SLC27A1 | CDIN1 | PBRM1 |
| SLC30A1 | TMEM107 | C1D |
| SLC31A1 | CXXC5 | ACOT9 |
| SLC33A1 | DNAJC5 | ACOT9 |
| SLC35A5 | DNAJC5 | SINHCAF |
| SLC35E4 | DNAJC5 | BEAN1 |
| SLC37A2 | TBC1D14 | MYG1 |
| SLC39A4 | DYNC2I2 | JDP2 |
| SLC39A7 | SUN1 | SEC16B |
| SLC6A8 | GPCPD1 | ZNF704 |
| SLC7A7 | GPT2 | TIFA |
| SLC9A1 | STING1 | CASD1 |
| NHERF2 | STING1 | BRCC3 |
| VASN | GNG2 | MAD2L2 |
| SMARCAD1 | DUS3L | MAN1A2 |
| SMOX | H19 | MAP2K7 |
| SNCG | CMTM4 | MAP3K1 |
| SNX16 | CMTM4 | MAP3K4 |
| SNX2 | APCDD1 | MAP3K5 |
| SOX4 | DAB2IP | MAP4K3 |
| SP100 | EPB41 | MAP4K5 |
| SP6 | SIPA1L2 | MAPK3 |
| SPA17 | ZNFX1 | MAPK8 |

| | | |
|---|---|---|
| SPC25 | MDM2 | MAPKAP1 |
| SPHK2 | CRAMP1 | MAPRE2 |
| SPINT1 | KIF3B | MAPT |
| SPON2 | SLC25A25 | MARCKS |
| SPR | ZKSCAN1 | MBD2 |
| SRC | SMAD5 | MBD3 |
| SRF | NOL4L | MBTPS1 |
| SRMS | HS6ST1 | MCCC2 |
| SRP14 | SMAP2 | MCM2 |
| SRR | COL27A1 | ANAPC1 |
| SSX2IP | ETNK1 | MCRS1 |
| ST14 | COL27A1 | MDFI |
| ST3GAL2 | COL27A1 | MDFIC |
| ST6GALNAC4 | ATOSA | MDN1 |
| STAC2 | ARL8A | ME2 |
| STARD10 | ZNF275 | MECP2 |
| STARD5 | ZNF275 | MED25 |
| STAT1 | DEDD2 | MEF2A |
| STAT2 | ABHD17C | METRN |
| STAU1 | AGPAT3 | METTL1 |
| STIM1 | AMOTL1 | MGA |
| STK17B | AMOTL1 | MGST3 |
| STRAP | TMEM128 | MID1IP1 |
| STX3 | SOGA1 | MINPP1 |
| SURF4 | MFHAS1 | DCP1A |
| SYVN1 | DOCK8 | ZBTB17 |
| TACC3 | KIAA1671 | MKI67 |
| TAF13 | ACAP3 | MKKS |
| TAPBP | INAFM2 | KMT2A |
| TBC1D1 | LIFR | KMT2C |
| TBC1D8 | FBXO25 | KMT2E |
| TBX2 | HMGB3 | MLLT10 |
| TCF7 | GLIPR2 | MLLT3 |
| TFCP2L1 | GLIPR2 | MARCKSL1 |
| TCIRG1 | BCL2L11 | FAR1 |
| SERINC3 | ARHGAP27 | MMAB |
| TEAD3 | CIPC | MNAT1 |
| TESK1 | CCDC117 | MNT |
| TEX9 | SPNS2 | MORF4L1 |
| TGFB3 | CTHRC1 | MORF4L2 |
| TGM2 | FAM83D | MOSPD2 |

| | | |
|---|---|---|
| TGOLN2 | FBRSL1 | MPDZ |
| TGOLN2 | FBRSL1 | MPHOSPH10 |
| THBS1 | CEP95 | PALS2 |
| TIMM22 | SORBS2 | MRGPRF |
| TIPARP | TUBGCP6 | MRPL12 |
| TK2 | TSPAN33 | MRPL15 |
| TLR2 | CGNL1 | MRPL17 |
| TLR3 | CCNK | MRPL19 |
| TMC6 | LUZP1 | MRPL23 |
| TMCC2 | ZNF746 | MRPL3 |
| TMED5 | VASN | MRPL32 |
| DOLK | TYSND1 | MRPL34 |
| TMEM19 | KIF1B | MRPL40 |
| TMEM38A | MPRIP | MRPL50 |
| TMEM40 | LINC01002 | MRPS16 |
| TMEM41A | NPNT | MRPS28 |
| SHISA2 | TP53INP1 | MRPS5 |
| TMOD1 | FNIP2 | MS4A8 |
| TNFRSF18 | FNIP2 | MSH3 |
| TNFRSF21 | RNF213 | MSN |
| TNNI1 | RNF213 | MSRA |
| TNPO2 | ZNF655 | MSRB2 |
| TOB2 | NRBP2 | MTAP |
| TOP3A | METRNL | MAP2 |
| TOR2A | TMEM64 | MAP6 |
| TOR3A | MOB1B | MTBP |
| TPCN1 | GAB1 | MTCH2 |
| TRPT1 | GAB1 | MTDH |
| TRAF4 | CABLES2 | MTF2 |
| TREX1 | ZNF12 | MTHFD2 |
| TRIM11 | GPSM1 | MTHFS |
| TRIM5 | SCAF4 | SBF2 |
| TRIM17 | ZNF385A | MTMR2 |
| TRIM21 | ZNF436 | MTMR3 |
| TRIM5 | ZNF436 | MTMR4 |
| TRIM32 | RTL10 | MTSS1 |
| TRIM34 | ZSWIM6 | MUC4 |
| TRIM41 | MEG3 | BLOC1S5 |
| TRIM7 | INSR | MUTYH |
| ZNHIT3 | GDE1 | MVK |
| TP53INP1 | KDM2B | MYADM |

| | | |
|---|---|---|
| TP53INP2 | INSR | MYB |
| TSPAN13 | RIC1 | MYBBP1A |
| TSPAN7 | MCC | MYC |
| TTC13 | ELAPOR1 | MYNN |
| TTC16 | ATP8B1 | MYO10 |
| TTLL1 | LYNX1 | MYO1B |
| TUBA4A | ADAMTS2 | MYO1E |
| TUFT1 | ADSS1 | MYO9A |
| GLRX3 | MIGA2 | MYOC |
| CMPK2 | ABCC5 | MYOD1 |
| UBA7 | CXADR | KAT7 |
| UBE2E1 | C19orf25 | NAGA |
| UBE2I | ARL14EP | NAGK |
| UBE2V2 | C18orf25 | NAP1L3 |
| UBN1 | DUSP22 | NAA15 |
| UFD1 | HES6 | NASP |
| UGGT1 | INSR | NBEA |
| UGT1A1 | FNIP2 | NCK2 |
| UGT1A8 | DIPK2A | NCOR1 |
| UGT1A3 | KBTBD6 | NDFIP1 |
| UGT1A4 | NRARP | NDN |
| UGT1A6 | ZNF71 | NDST1 |
| UGT1A8 | ORAI1 | NDUFB4 |
| UHMK1 | EEIG2 | NDUFS7 |
| ULK1 | C12orf76 | NDUFS8 |
| UNC5C | SNHG14 | NEDD4 |
| UPF3B | C16orf87 | NEDD4L |
| UPP1 | UBE2QL1 | NEIL3 |
| PDE4DIP | FAM167A | NEK3 |
| USP18 | ARHGAP23 | NEK7 |
| USP2 | NATD1 | NET1 |
| USP20 | RORA | NF1 |
| USP27X | ADCYAP1R1 | NF2 |
| USP38 | CMPK2 | NFATC1 |
| UTRN | RNF169 | NFATC3 |
| VAX2 | ADAM12 | NFIB |
| VEGFA | SLC25A30 | NFIC |
| VPS18 | ABHD15 | NFIX |
| DCAF4 | FITM2 | NFKB1 |
| WDR45 | TENT5C | NFYA |
| WFS1 | STOX2 | NFYC |

| | | |
|---|---|---|
| ZMAT3 | GXYLT1 | BEX3 |
| CCN4 | MEGF6 | NICN1 |
| WNT7B | ATG4D | NIPBL |
| WSB1 | HVCN1 | NIPSNAP3A |
| WSB2 | LRRN1 | NISCH |
| XAB2 | UNC5B | NKD2 |
| XRCC1 | EGFLAM | NMT1 |
| YPEL5 | GUSBP1 | NOP58 |
| ZAP70 | EAF1 | NOP56 |
| ZBP1 | KIF1B | NOP10 |
| PARP12 | ZNF777 | NOLC1 |
| TUT7 | FGD5 | NONO |
| MORC4 | FGD5 | NOTCH2 |
| ZNF185 | WIPI2 | NOTCH4 |
| ZNF219 | PLD6 | NPM1 |
| ZPR1 | C11orf96 | CYB5R1 |
| PATZ1 | ZBTB34 | NR1D2 |
| ZNF281 | FOXP4 | NR1I3 |
| ZNF296 | ZBTB20 | NR2C2 |
| ZNF346 | LINC01002 | NR3C1 |
| ZFP36 | NAAA | NRAS |
| ZNF385A | BID | NRF1 |
| KLF17 | TNRC6C | NRIP1 |
| ZNF426 | DNAJC18 | NRM |
| ZNF503 | DNAJC18 | NRP2 |
| ZNF622 | PTAFR | NSFL1C |
| ZNF253 | AFF3 | NSMCE1 |
| ZNF394 | DIP2A | NT5C |
| ZFYVE21 | ZMAT3 | NT5E |
| ZNRF1 | CRACD | NTNG1 |
| ZNRF2 | CRACD | NUBP1 |
| ZRANB1 | LRP10 | NUDCD1 |
| | FGF11 | NUDCD3 |
| | ITGA9 | NUDT2 |
| | DIDO1 | NUDT4 |
| | BEND7 | NUMB |
| | HS3ST3B1 | NUP107 |
| | TUSC1 | NUP133 |
| | TBCEL | NUP160 |
| | PTPRJ | NUP58 |
| | FAM43A | NUSAP1 |

| | |
|---|---|
| TNFAIP8L1 | NUTF2 |
| CNKSR3 | NXT1 |
| SOX6 | MBOAT1 |
| SOX6 | TENM4 |
| ITPRIPL2 | OGG1 |
| CMBL | OGT |
| GLCCI1 | OLFML2B |
| WDR20 | OPHN1 |
| CBX4 | ORC3 |
| DIS3L2 | ORMDL1 |
| TSPAN11 | OSBPL6 |
| RIPOR3 | OSBPL9 |
| ATP23 | OSGEP |
| SMAD9 | OSMR |
| RNF166 | OSR1 |
| LYPD6 | OXA1L |
| ITPRIPL2 | P2RX4 |
| ZFP62 | P2RY2 |
| TLCD1 | PABPN1 |
| FGD3 | PAFAH1B1 |
| NTN1 | PAFAH1B2 |
| SMIM10L2A | PAFAH1B3 |
| ASB2 | PAICS |
| ZYG11B | PAK1 |
| LIN7A | PAK3 |
| ITPRIPL2 | PANK1 |
| FRMD8 | PANK3 |
| GPR157 | TENT2 |
| NRK | PAPOLA |
| RILPL2 | PAPSS1 |
| ZNF343 | PARD3 |
| ZBTB43 | PARD6A |
| CEP85L | PARD6B |
| POLR1H | PARN |
| CRACDL | PARP16 |
| GOLGA7B | PARP2 |
| MYLIP | PASK |
| KCNK3 | PAXIP1 |
| ZNF398 | PBK |
| RNF144B | PBX1 |
| PRXL2A | PBX2 |

| | |
|---|---|
| S1PR3 | KAT2B |
| DISP1 | PCBD2 |
| C2orf88 | PCBP4 |
| SOX6 | PCCA |
| FCHO2 | PCDHB18P |
| MIB2 | PCK2 |
| MAP7D2 | PCM1 |
| DEPDC7 | PCNX3 |
| CNIH4 | CDK16 |
| EMILIN3 | PDAP1 |
| ZMAT3 | PDE6D |
| BICDL1 | PDE7A |
| KLHL28 | PDE8A |
| PLIN4 | PDGFRA |
| KIF9 | PDGFRB |
| AIFM2 | PDHA1 |
| MAP6 | PDK1 |
| PLEKHA7 | PDK2 |
| GID4 | PDZRN3 |
| ZNF319 | PER2 |
| CDRT4 | PER3 |
| POLR1A | GATB |
| NANOS1 | PEX11A |
| MIR29B2 | PEX14 |
| ABTB3 | PEX7 |
| FAM83G | PFKL |
| UBXN2A | PFKM |
| XAF1 | PFKP |
| PEAR1 | PGLS |
| TMEM144 | PHB1 |
| ZNF37A | PHF12 |
| TOX2 | PHF13 |
| GRK3 | PHF21A |
| BVES | PHF3 |
| ALG9 | PHIP |
| GNG7 | PHKA1 |
| AMOTL1 | PHKA2 |
| CDT1 | PHLDB2 |
| COL22A1 | PIGC |
| FAM162B | PIGO |
| TUB | PIK3C2A |

| | |
|---|---|
| SEMA4D | PIK3C3 |
| ADAMTS7 | PIK3R1 |
| DANT2 | PIPOX |
| DNASE1 | PITPNB |
| LACC1 | PKP4 |
| INTU | PLA2R1 |
| CARNS1 | PLCE1 |
| DSTYK | PLD1 |
| GAB1 | PLD2 |
| JPH1 | PLEKHA5 |
| KAZN | FERMT2 |
| USP35 | PLEKHF1 |
| LRRC37B | PLK4 |
| LRRC57 | PLP2 |
| JPH3 | PLSCR1 |
| GPATCH2L | PMF1 |
| SPATA18 | PMPCB |
| ADAMTS5 | PNN |
| IHH | POFUT1 |
| GRTP1 | POLA1 |
| ILDR2 | POLB |
| HOXD10 | POLD1 |
| TPST2 | POLD2 |
| MICAL3 | POLD3 |
| ZNF367 | POLE |
| DISP2 | POLE2 |
| GRAMD2A | POLI |
| ERFE | POLM |
| EML6 | POLR2I |
| GPATCH2L | POLR3A |
| ZBTB42 | POLR3F |
| MDM2 | POLR3G |
| SMTNL2 | PORCN |
| FOXS1 | POT1 |
| ANO4 | POU2F1 |
| COL4A4 | PLPP3 |
| DCLK1 | PLPP2 |
| CCN4 | PPARGC1B |
| LINC00526 | PPAT |
| WIPF3 | PPIL4 |
| UNKL | PPP1CC |

| | |
|---|---|
| SHE | PPP1R3C |
| CLN8 | PPP2R1B |
| EVC2 | PPP2R3A |
| SHISA9 | PPP3CA |
| RNF213 | PPP3CC |
| DNAAF3 | PQBP1 |
| CNTN2 | PRELP |
| PROSER2 | PRIM2 |
| RIMBP3 | PRKACB |
| GDNF | CAVIN3 |
| ZXDC | PRKD3 |
| MCOLN2 | PRKCZ |
| SMIM43 | PRKDC |
| F2RL2 | THAP12 |
| CDH24 | PRNP |
| NIPAL4 | ERI3 |
| HIC1 | PRPF31 |
| SNORC | PRPF39 |
| LPAR5 | PRPSAP1 |
| NEURL2 | PRUNE1 |
| CST3 | PSENEN |
| NKX1-2 | PSIP1 |
| KNDC1 | PSTK |
| GLDN | PTBP2 |
| ENTPD2 | PTCD2 |
| S1PR5 | PTCH1 |
| TMEM52 | PTDSS2 |
| INSYN1 | TWF1 |
| DBH-AS1 | PTOV1 |
| GPCPD1 | PTPN11 |
| MCHR1 | PTPN12 |
| FAM182A | PTPN21 |
| RASGEF1A | PTPRA |
| GRIK3 | PTPRF |
| GJD3-AS1 | PTPRK |
| FRMD3 | PTPRS |
| P2RX7 | PTS |
| DENND2C | PUM1 |
| ZCCHC8 | PUM2 |
| GCNT2 | PURA |
| FRMD5 | PUS3 |

| | |
|---|---|
| GALNT10 | NECTIN2 |
| NOL4L | PYGB |
| POLH | QKI |
| GPR155 | RAB14 |
| SNX32 | RAB18 |
| ZNF570 | RAB2A |
| LINC02688 | RAB27B |
| AGAP3 | RAB28 |
| TRIM71 | RAB3D |
| KIF9 | RAB4A |
| TDRD10 | RAB4B |
| FGF18 | ERC1 |
| INAFM2 | RAB9A |
| CARD18 | RABGAP1L |
| S1PR3 | RABGGTA |
| GJB6 | RABGGTB |
| EOMES | RABIF |
| DLX3 | RACGAP1 |
| KIAA1217 | RAD18 |
| TIMM29 | RAD21 |
| HES2 | RAD23A |
| ELMOD1 | RAD23B |
| MARCHF8 | RAD51C |
| ZNF658 | RAD51B |
| RNF213 | RAD54L |
| STOX2 | RAF1 |
| TBCEL | RAI14 |
| ZNF30 | RANBP1 |
| GXYLT1 | RAP1GDS1 |
| KIF26A | RASL12 |
| KIF16B | RASSF1 |
| BCLAF3 | RBBP9 |
| RNF213 | RBL1 |
| EPPK1 | RBM10 |
| EPPK1 | RBM14 |
| MYORG | SCAF8 |
| SLC25A34 | RBM18 |
| KCNJ12 | RBM5 |
| KCNS2 | RBFOX2 |
| CDIN1 | RBMS1 |
| MEGF10 | RCE1 |

| | |
|---|---|
| TP73 | RCL1 |
| ZNF684 | RDH14 |
| GOLGA2P5 | RDX |
| LOXL1-AS1 | RFC1 |
| DOCK8 | RECK |
| OBSCN | REPS1 |
| FITM2 | REV3L |
| GOLGA2P5 | RFK |
| CNTNAP3 | COP1 |
| ONECUT2 | RGS11 |
| SIPA1L2 | RGS19 |
| CST11 | RHOBTB2 |
| TOR3A | RHOT2 |
| ASB2 | RHOU |
| CXXC5 | RIF1 |
| ESPN | RIPK1 |
| FAM95A | RNASEH1 |
| ESPNP | PCGF2 |
| KIF26A | RNF111 |
| STOX2 | RLIM |
| GMCL1 | RNF13 |
| ELOVL3 | RNF130 |
| ZNF470 | RNF138 |
| PHRF1 | RNF141 |
| SPRING1 | RNGTT |
| ABHD13 | RBM39 |
| SPICE1 | SNORD22 |
| VPS13C | ROCK1 |
| NXPE3 | ROCK2 |
| ADCY1 | PTBP3 |
| FBXL20 | ROR1 |
| TPM4 | RP2 |
| RASEF | RPA3 |
| ZNF561 | RPL22 |
| KSR1 | RPL3 |
| FAM83F | RPL30 |
| GPATCH2L | POLR1B |
| VASH2 | POLR1D |
| RMDN2 | POLR1A |
| MR1 | RPP14 |
| PLEKHM3 | RPP21 |

| | |
|---|---|
| ADAMTS5 | RPP40 |
| CFLAR | RPS19 |
| CCDC134 | RRBP1 |
| DDX31 | RRM1 |
| SMAD5 | CLIP1 |
| KCNC4 | RSRC1 |
| NEU3 | RTTN |
| LRPAP1 | RUNX1 |
| ASPRV1 | RUNX2 |
| RNF144B | RUSC2 |
| VPS37D | RYK |
| ILDR1 | S100B |
| MAP6 | SASH1 |
| EXT2 | MSMO1 |
| ZFP90 | SC5D |
| GANC | ATXN2 |
| KLHL28 | ATXN7 |
| TPST2 | SCAMP4 |
| LINC00472 | SCD |
| FRK | SCD |
| DOC2B | SPATS2 |
| DQX1 | SDC3 |
| WDR1 | FRRS1 |
| LTB4R | SEC14L1 |
| GGT6 | EXOC6 |
| ODF3L1 | SEC24D |
| ITGAV | EXOC4 |
| RORA | SEH1L |
| KRBA1 | SEMA6D |
| ADCYAP1R1 | SENP1 |
| EPB41 | SENP6 |
| WRAP73 | SERPINB1 |
| WRAP73 | SERTAD3 |
| CHRNA7 | SESN1 |
| ANO4 | SESN3 |
| ENSG0000026ç | SF3B3 |
| MEGF10 | SRSF1 |
| PRDM6 | SREK1 |
| LLGL1 | SRSF2 |
| KALRN | SCAF11 |
| LINC01719 | SRSF3 |

| | |
|---|---|
| C4orf47 | SRSF6 |
| COL27A1 | SRSF7 |
| C4orf47 | SFXN1 |
| ACO1 | SFXN4 |
| CST3 | SGTB |
| SLC7A1 | SH3BGRL |
| MDM2 | SH3BP5 |
| EDDM13 | ITSN2 |
| HEPACAM | SH3KBP1 |
| C6orf132 | SH3RF1 |
| SLC16A14 | SHKBP1 |
| ARID1B | SIN3A |
| TRIM8 | GEMIN2 |
| RBP7 | SIRT4 |
| RTCA | SKI |
| DNAJC18 | SKP2 |
| USP6NL | SLC12A6 |
| GATA5 | SLC19A1 |
| UQCC5 | SLC20A2 |
| FGFBP3 | SLC23A2 |
| TTC39C | SLC25A10 |
| TRAF3IP1 | SLC25A13 |
| ULBP2 | SLC25A24 |
| BRD7 | SLC29A2 |
| ZNF296 | SLC30A6 |
| KIF26B | SLC35A3 |
| FYB2 | SLC39A11 |
| ABTB3 | SLC39A3 |
| IRF1 | SLC4A3 |
| ENSG00000288 | SLC4A4 |
| MDM2 | SLC6A2 |
| RASSF10 | SLC7A2 |
| KCNJ5 | SLC9A8 |
| LRRTM1 | SLK |
| ARX | SMAD4 |
| BRWD1 | SMARCA2 |
| ZNF420 | SMARCAD1 |
| ACER3 | SMARCB1 |
| EFNA2 | SMARCE1 |
| SLC66A1L | SMC5 |
| FOXC2 | SMC6 |

| | |
|---|---|
| ZNF786 | SMR3B |
| HOGA1 | SMYD2 |
| MARVELD3 | SMYD5 |
| CXADR | SNX18 |
| CEP43 | SNAI2 |
| GMCL1 | SNAPIN |
| IRF5 | SNRPA |
| LRRC8E | SNX14 |
| CEP43 | SNX15 |
| ZNF284 | SNX5 |
| GPATCH2L | SNX9 |
| ZNF708 | SOCS2 |
| GPR155 | SOCS6 |
| HS3ST6 | SOD2 |
| GJA3 | SON |
| TLR3 | SOX12 |
| CFLAR | SOX6 |
| FOXO6 | SP3 |
| TRIM7 | SPAG7 |
| RNF144B | SPCS3 |
| PGAP1 | SPDEF |
| PYGO1 | SPEN |
| RNU6-37P | ERLIN1 |
| KLC3 | SPART |
| ONECUT2 | SPHK1 |
| ARID1B | SPIN1 |
| MAP7 | SPTAN1 |
| FLRT2 | SPTBN1 |
| ZNF341 | SPOP |
| SLC32A1 | SPPL3 |
| LINC01120 | SPRED1 |
| INSM2 | SPRED2 |
| ANKRD20A11P | SPRY1 |
| CNNM3 | SQLE |
| ZNF654 | TOM1L1 |
| HR | SREBF2 |
| TMEM65 | SRGAP1 |
| C6orf132 | SRGAP2 |
| CALML3-AS1 | SRGAP3 |
| ZNF440 | SRPK2 |
| AK3P3 | SRPRB |

| | |
|---|---|
| PGAP1 | SSBP1 |
| ZNF654 | SSPN |
| ABHD4 | SSR1 |
| ERVK3-1 | ST13 |
| CHRM4 | ST3GAL1 |
| WFDC5 | ST3GAL5 |
| NRG2 | ST7L |
| ESPN | STAG1 |
| TMEM64 | STAG2 |
| PLEKHA7 | STAM2 |
| FGD4 | STARD4 |
| ZNF600 | STAT5A |
| KISS1R | STAT5B |
| LRPAP1 | STAT6 |
| MED23 | ELP2 |
| LMO7 | STC1 |
| ZNF420 | STK3 |
| KLRK1-AS1 | STOML2 |
| ZNF234 | STX12 |
| VWCE | STX17 |
| CABP7 | STX4 |
| ZNF283 | STX8 |
| FRRS1 | SUCLG2 |
| C1QTNF12 | ZNF280C |
| WDFY2 | ZNF280D |
| GNA14 | SULT2B1 |
| PRAP1 | SUOX |
| CNTNAP3 | SUV39H1 |
| BLNK | KMT5B |
| PGAP1 | SUZ12 |
| INSYN2A | SVIL |
| ADAM23 | SYNCRIP |
| SHISA8 | TACC2 |
| ZNF850 | TADA3 |
| OTULINL | TAOK1 |
| MTUS2 | TARS1 |
| ELF4 | TARS2 |
| CHST3 | TBC1D17 |
| JAG2 | TBC1D19 |
| ZNRD2 | TBCE |
| GDPD5 | TBL1X |

| | |
|---|---|
| ARAP1 | TBL1XR1 |
| CACNB3 | TBL2 |
| MCF2L | TAF8 |
| LDB1 | TBRG4 |
| ARHGAP19 | TCERG1 |
| HBEGF | TCF12 |
| HIP1R | TCF19 |
| AQP3 | TCF3 |
| AQP3 | TCF4 |
| GAK | TCF7L2 |
| GNA11 | TFAP4 |
| FBXO41 | GRHL2 |
| SDHAF1 | TCF3 |
| ABHD6 | VPS72 |
| NAA60 | TCOF1 |
| CANT1 | TCP1 |
| LZTS1 | TCP11 |
| WHRN | DYNLT3 |
| STRADA | DYNLT1 |
| SHISAL1 | TDP1 |
| MARK4 | TNS2 |
| ELMO2 | DCUN1D1 |
| GNA11 | ZFAND3 |
| LDLRAP1 | TFB1M |
| ZGPAT | TFRC |
| ARHGEF40 | TGFB2 |
| CX3CL1 | TGFBR3 |
| | THOP1 |
| | ZBTB7B |
| | THRA |
| | THRAP3 |
| | MED16 |
| | THRSP |
| | THSD1 |
| | ADAMTSL5 |
| | TIA1 |
| | TIMELESS |
| | TIMM10 |
| | TIMM44 |
| | TIMM9 |
| | TIMP3 |

TK1
TLE2
TLE6
TLR6
GPR137B
CEMIP2
TMEM39A
TMLHE
TMPO
STEAP4
TNFSF13
TNNC2
TNPO3
MED12
TNRC6A
TNRC6B
TOM1
TOMM40
TOP2A
TPBG
TPD52L2
TPK1
TPM3
TPR
PDSS1
TRAF2
TRAF6
TF
TRIB1
TRIB2
TRIB3
TRIM16
TRIM2
TRIM44
TRIO
TRIP11
TRIP13
TRIP4
TRPC2
TRPM7
TRPS1

TRUB2
TSC22D4
CEP41
CRLF2
TSPAN15
TSPAN2
TSPAN5
TTC14
TTC3
TTC5
TTC8
TTYH2
TUBA1A
TUBB
TXNIP
TXNRD3
TYMS
U2AF1L4
UBAP2L
UBA3
UBE2C
UBE2D3
UBE2E2
UBE2E3
UBE2J1
UBE2S
UBE3A
UBE4B
UBL4A
SAE1
UBA2
UBR1
UBR2
UBXN6
USP1
USP21
USP22
USP34
USP36
USP4
USP47

USP9X
UTRN
VAMP3
VANGL1
VASP
VAV3
VCAM1
VPS54
VRK3
VTI1A
VTI1B
WAC
WASL
RCC1L
METTL27
WDFY3
IFT122
DCAF5
WDR33
WDR4
DCAF8
BRWD1
WEE1
NELFA
CCN5
WNT5A
WWOX
XBP1
XPA
XPO1
XPO4
XPO7
XPR1
XRCC4
XRCC5
YEATS4
YES1
YTHDF3
YWHAH
YWHAZ
ZBED3

ZBTB12
ZBTB20
ZCCHC14
ZCCHC7
ZCCHC8
ZDHHC3
ZDHHC5
ZDHHC6
ZEB2
ZNF638
ZNF124
ZNF142
ZNF148
ZBTB14
ZSCAN26
ZNF207
ZNF212
ZMYM3
ZNF277
ZNF282
ZNF292
ZBTB22
ZKSCAN4
ZNF318
ZNF322
ZNF519
ZNF398
ZNF22
ZNF445
ZNF467
ZNF521
ZNF18
ZNF644
ZFP82
ZNF565
ZNF28
ZFP90
ZFP91
ZNF235
ZSCAN12
ZFPM1

ZSCAN21
POLR1H
